# A pragmatic approach to produce theoretical syntheses in ecology

**DOI:** 10.1101/344200

**Authors:** Bruno Travassos-Britto, Renata Pardini, Charbel N. El-Hani, Paulo I. Prado

## Abstract

Theoretical syntheses have the role of describing and guiding knowledge generation, and are usually done by enunciating the conceptual bases that guide research in a given field. In fields that develop axiomatically, the conceptual basis can be easily identified in the set of axioms guiding model building. However, ecology does not develop axiomatically but rather pragmatically, i.e., ecologists do not build models based on a predefined set of assumptions (axioms). They rather resort to any information that seems useful to learn about ecological phenomena. Therefore, a theoretical synthesis in ecology cannot rely on the enunciation of axioms; instead, it requires identifying what information and knowledge ecologists use (i.e., what they decide is useful to learn). Here we present an approach for producing theoretical syntheses based on the information/knowledge most frequently used to learn about the world. The approach consists of (i) defining a phenomenon of interest; (ii) defining a collective of scientists studying the phenomenon; (iii) surveying the scientific studies about the phenomenon published by this collective; (iv) identifying the most relevant publications used in these studies; (v) identifying how the studies refer to the most relevant publications; (vi) synthesizing what is being used by this collective to learn about the phenomenon. We implemented the approach in a case study on the phenomenon of ecological succession, defining the collective as the scientists currently studying succession. We identified three propositions that synthesize the views of the defined collective about succession. The theoretical synthesis revealed that there is no clear division between “classical” and “contemporary” succession models, and that neutral models are being used to explain successional patterns alongside with models based on niche assumptions.By implementing the pragmatic approach in a case study, we show that it can be successfully used to produce syntheses describing the conceptual bases of a field, which have the potential to guide knowledge generation. As such, these syntheses fulfil the roles ascribed to scientific theories in the epistemological literature.

## Introduction

Theoretical syntheses describe the conceptual bases and guide knowledge generation in a field of study [1, 2]. However, the conceptual bases of knowledge generation in ecology might not be so easy to identify. The discussions about theory in ecology led to no clearly agreed conceptual bases unifying all scientific activity in this science [3–5] and attempts to synthesize ecological theory [6–8] were not generally taken into account in subsequent work (as one would expect from syntheses meant to describe and guide knowledge generation in a field). Regardless of this lack of consensus and reference to theoretical syntheses, ecology has undergone intense development as a science for more than a century of history [5, 9–11]. Hence, what is guiding the development of ecology and how can we make meaningful theoretical syntheses in this field?

The traditional format of a theoretical synthesis is based on the enunciation of axioms — statements assumed to be self-evident in a specific context [2] — followed by immediate theorems — deductions from these axioms that can be used to both support the truth of the axioms and make predictions about the world [12]. This kind of synthesis is aligned with axiomatic views of how a theory describes and explains phenomena and guides knowledge generation. In the most recent understandings of axiomatic views, if theorems deduced from the axioms represent accurately a phenomenon, this theorem will be regarded as a model of the theory [2]. If a phenomenon is adequately represented by the model and, therefore, behaves like the model, the axioms from which the models were deduced can be regarded as true of the world [12]. Therefore, a synthesis enunciating a set of axioms and how they can be combined to generate new models will be very useful to describe the conceptual basis of a field of study and should help guide scientific activity by clearly enunciating the accepted rules to build new models. In the natural sciences, the axioms of a successful theory are presented as laws of nature, which sometimes are viewed as exceptionless rules according to which nature behaves and sometimes as general rules that help us identify major tendencies in nature [11, 13, 14]. Either way, if these laws are assumed as true and models to learn about the world are deduced from them, these laws will be working as axioms of a theory. The adoption of such views of theory might explain why some authors interested in making theoretical syntheses in ecology are also interested in the existence and identification of laws in this science [15–18]. However, there are views of theory that do not assume that models used to learn about phenomena are, or at least should be, deduced from axioms [2].

In Travassos-Britto et al. [19], we proposed that knowledge generation in ecology is often not axiomatic but rather pragmatic. Under the pragmatic view of theories, scientific research involves using models to learn about phenomena, but there is no concern in following a set of rules enunciated in axioms. Rather, models are built freely by resorting to any knowledge available to the modeler such as previously proposed models, propositions, methods, definitions [20]. There are many different criteria for a model to be considered useful, including cognitive criteria (e.g., predictive efficacy, explanatory success, increased understanding, adequacy-for-purpose) [21–23] and social criteria (e.g., prominence of its proponents) [24]. The final decision on which model is useful and which is not is made by the agents (in this case, practicing ecologists) when carrying out their studies. As a consequence, the set of models used to learn about a phenomenon in a field of study is defined by the decisions of a collective of agents trying to learn about the phenomenon, not by their deductive relations with axioms. In this scenario, a theoretical synthesis based on the collective decisions of a community should better reflect the conceptual bases of a field of study than a set of axioms and deduced theorems. In this scenario, the question that remains is how to access the decisions of such a collective.

Scientists report their research mostly in written articles where they make propositions about how the world is [25]. Propositions made in past studies are often referred to in subsequent studies to inform readers of which views about the world the research is based upon. Therefore, by accessing the article reporting a scientific study it is possible to trace which propositions are being used as conceptual basis for that study and how they are being used [26]. The same applies for a collective of scientists studying a phenomenon. By accessing the articles describing the scientific studies of a collective, one can discover which and how propositions are being used as conceptual bases to learn about the world from the point of view of that collective. Propositions about the world that are frequently referred to by a scientific community may reveal, among other things, that some propositions seem particularly relevant to learn about the phenomenon and other propositions do not.

Here, we propose a methodological approach to identify and describe the most referred propositions in the scientific research intended to learn more about a certain phenomenon. We provide an example of the implementation of the approach used to identify the current conceptual bases of the studies about ecological succession and describe the specific methodological decisions we made to implement each step as well as the results obtained. We end up by discussing in what way our proposed approach to produce a theoretical synthesis is different from other approaches and how syntheses produced by using our approach can fulfill the role of a clearly enunciated theory.

### The Pragmatic Approach to Theories

The notion that models validity corresponds to how much a model is used in a given context is based on the pragmatic view of scientific theories [2]. Therefore, we dubbed our approach Pragmatic Approach to Theories (PATh). We outlined PATh as a set of steps that can be executed by adopting different methods. Here, we describe this set of steps by pointing out what each is supposed to achieve. Afterwards, we will describe how we specifically implemented each step in a case study about ecological succession.

The approach we are proposing consists of the following steps (see also Fig 1):

**Fig 1.**
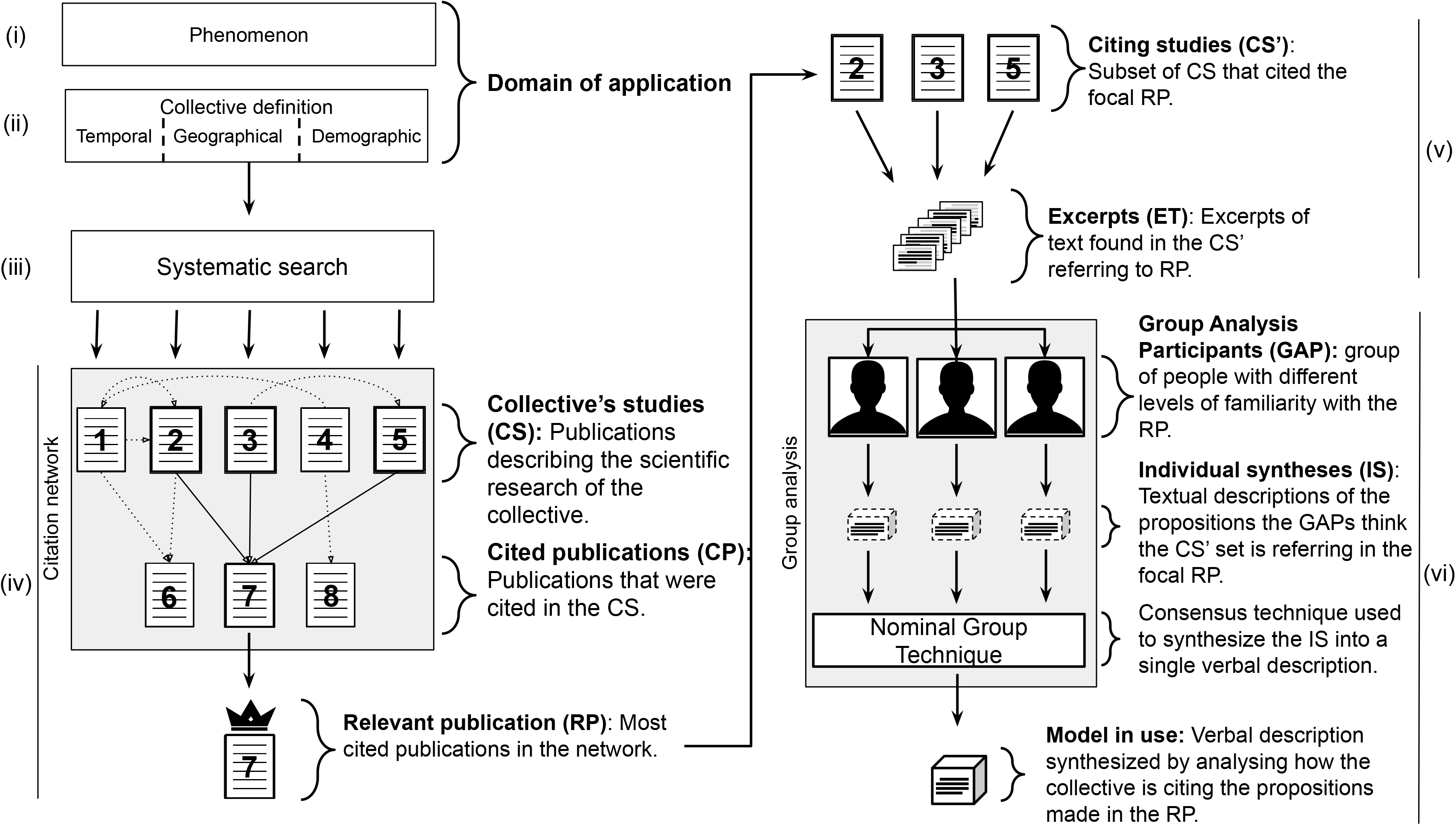
PATh Workflow. Workflow depicting how the information obtained in each of the steps of the PATh (roman numerals) is used in the following step aiming to go from the definition of the domain of application to the final models in use. Note that the steps (v) and (vi) are carried out for each relevant publication in the set defined in step (iv).

#### I Definition of the phenomenon of interest

The goal is to define the object of the theory. In the natural sciences, theories are about natural phenomena. Defining the phenomenon of interest should specify which concepts and corresponding terms to look for in the scientific literature. The output of this step should be a set of terms that circumscribe which studies are regarded to be about the

#### II Definition of the phenomenon of interest

The goal is to define from whose point of view one wants to make a theoretical synthesis about the phenomenon. Defining the collective of agents will give hints on where to look and how to filter all the activities aimed at learning about the phenomenon of interest. The output of this step should be a set of search parameters related to time scope, geographic, demographic and academic profile of the focal collective of agents studying the phenomenon of interest.

#### III Survey of the scientific activity

The goal of this step is to take a picture of the activity of the focal collective investigating the phenomenon of study. This collective may present their activities in different ways. Currently, the most used way to report a scientific study is in academic journals. With the parameters resulting from the previous step, one will be able to conduct a search aimed at recovering the reports of the studies of the focal collective about the phenomenon of interest. The output of this step is a set of publications that should be a meaningful sample of the publications of the collective aimed at learning about the phenomenon.

#### IV Identification of relevant publications

The goal of this step is to identify the publications that were most referred to due to their conceptual contribution for the scientific activity of the collective of scientists defined in step (iii). The output of this step should be a set of most referred publications among those identified in step (iii).

#### V Excerpting

The goal of this step is to identify, among all the information contained in the relevant publications, what is effectively being referred to by the community in their scientific activity and how it is being cited. The result of this step is a set of excerpts of text containing the citation of the relevant publications as described in the studies carried out by the collective of agents.

#### VI Content analysis of excerpts

The goal is to rebuild self-consistent descriptions of the focal phenomena or other related phenomena based on the content of the citations of the relevant publications. Common statements among the citations are used to rebuild the self-consistent descriptions, which we refer to as “models in use”. The result of this step is a set of statements that synthesize the main contributions of the relevant publications used by the collective to learn about the phenomenon.

The approach assumes that science works as a decentralized system of information exchange [27]. This system has been described in many ways, e.g., as cycles of normal science followed by paradigm shifts [28], as heterogeneous networks of actors [29], or as distributed cognitive systems [30]. Our approach is agnostic on the details of the dynamics of scientific information exchange systems, provided that such dynamics includes the use of models (as defined above).

The steps included in PATh were thought to give access to the activities of the collective of agents studying a phenomenon while dealing with issues associated with assuming that this decentralized system of information exchange can be traced in the network of citations among publications. The first issue with that assumption is that among the publications that are cited in a network of citations there is a continuum that goes from the most cited to the least cited ones. Therefore, there needs to be cut-off criteria to define what is a relevant publication and what is not. A second issue is that the citation index alone does not necessarily reflect theoretical links between citing and cited publication [26, 29, 31]. Therefore, it is necessary to differentiate citations that reveal theoretical links from other types of citations. The third issue is that even when a publication is cited due to its theoretical relevance in a field of study, it is most likely that only a particular part of it is being considered relevant and not all the propositions contained in it. Therefore, it is necessary to identify what part of the relevant publications is actually being referred to by the studies being carried out by the collective.

The steps (iv), (v) and (vi) were designed to deal with these issues and in doing so they differentiate this approach from a simple analysis based on a systematic search. In these steps, one should define criteria to reach a finite set of relevant publications, to differentiate theoretically relevant citations from non-relevant ones, and to identify what the collective of agents is actually referring to in the relevant publications. In the following section we will describe specifically how we implemented each step, taking ecological succession as a case study.

## Methods

### Defining the phenomenon of interest

We have chosen succession as a case study because it is a phenomenon that is referred to by a single term (“succession” itself), with a small degree of ambiguity or polysemy, compared to other ecological concepts [32]. We realize that this choice leads to a very inclusive criterion about what is ecological succession, but despite that, it points to a well-circumscribed research field in ecology. Inclusive definitions are not problematic or uncommon in theoretical synthesis, as exemplified by Pickett, Meiners and Cadenasso’s (p. 187) [33] definition of succession, regarded by the authors themselves as very inclusive: “changes in structure or compositions of a group of organisms of different species at a site through time”. Our approach is, however, robust to other meanings ascribed to the term, as we will see in the following section.

### Defining the collective of agents

At this step we chose to limit our case study to a collective of scientists currently publishing their results in venues that are part of a bibliographic database with broad coverage of English-written papers. What is considered as “current” is rather arbitrary and, therefore, we used a common option in literature surveys, which typically include a range of 10 years. We decided that the present decade (at the time of the analysis) was an adequate scope of time, considering the relatively long time a new discovery about the world takes to change a field of study [34].

We did not restrict the geographical or demographic scope to a specific group within this collective beyond the coverage provided by the database, and we did not take additional measures to control for publication or citation biases that may exist in the database [35]. Therefore, the results we obtained are as geographically and demographically biased as the publications found in the database used in the analysis.

### Surveying the scientific activities

This step was accomplished by searching all publications that contained the term “succession” or “successional” in their title or abstract and were considered as ecological studies in the database in the past ten years from the date of the study. To do so we carried out a systematic search in the ISI Web of Science™ database. This database was chosen because it has a set tools for systematic search as well as output files that can be promptly used to make citation network analyses (see next section). We used the keyword “successi*” in the “Topic” field of the search engine of the database, which searches for the term in title and abstracts, and filtered the publications that were in the “Ecology” category. We filtered the search for publications dated from January 2007 to July 2017. This search resulted in 5,536 publications, which represent a sample of the documentation of the scientific activity related to studies on succession in the field of ecology during the past ten years.

### Identifying relevant publications

To identify the relevant publications, we built a citation network including the 5,536 publications representing the studies of the focal collective plus all publications directly cited by these studies using the software CitNetEplorer™ [36]. The publications reporting current studies of ecological succession were dated from 2007 to 2017, but the publications cited by the retrieved documents could be from any year. The whole network included a total of 29,398 publications of different types (articles, books, chapters, proceedings papers, etc.), with 245,210 connections among publications.

We excluded highly cited publications that described statistical tools or approaches used in the studies, because they did not describe any properties of the phenomenon of interest or the natural world.

To select the smaller set of the most cited publications that were referred to in the studies of the collective of agents (the 5,536 works identified in the previous step) we created a cut-off criterion based on the citation index and representativeness of publication. We added publications to the set of relevant publications sequentially, beginning with the most cited paper and then adding the following most cited, while checking the percentage of studies by the focal collective that cited this new set. The proportion of the 5,536 studies that cited at least one publication in the set tended to stabilize at 80% when the 25^th^ most cited publication was included in the set.

We repeated this procedure considering only direct citations, considering direct and 2nd-degree citations (citations of citing articles), and considering direct, 2nd and 3rd degree citations. In all cases, there seemed to be a threshold tending to stabilization with 25 publications in the set of relevant publications (Fig 2). This analysis showed that about 20% of the studies that reflect the scientific activities of the focal collective were not citing the 100 most cited papers in the network.

**Fig 2.**
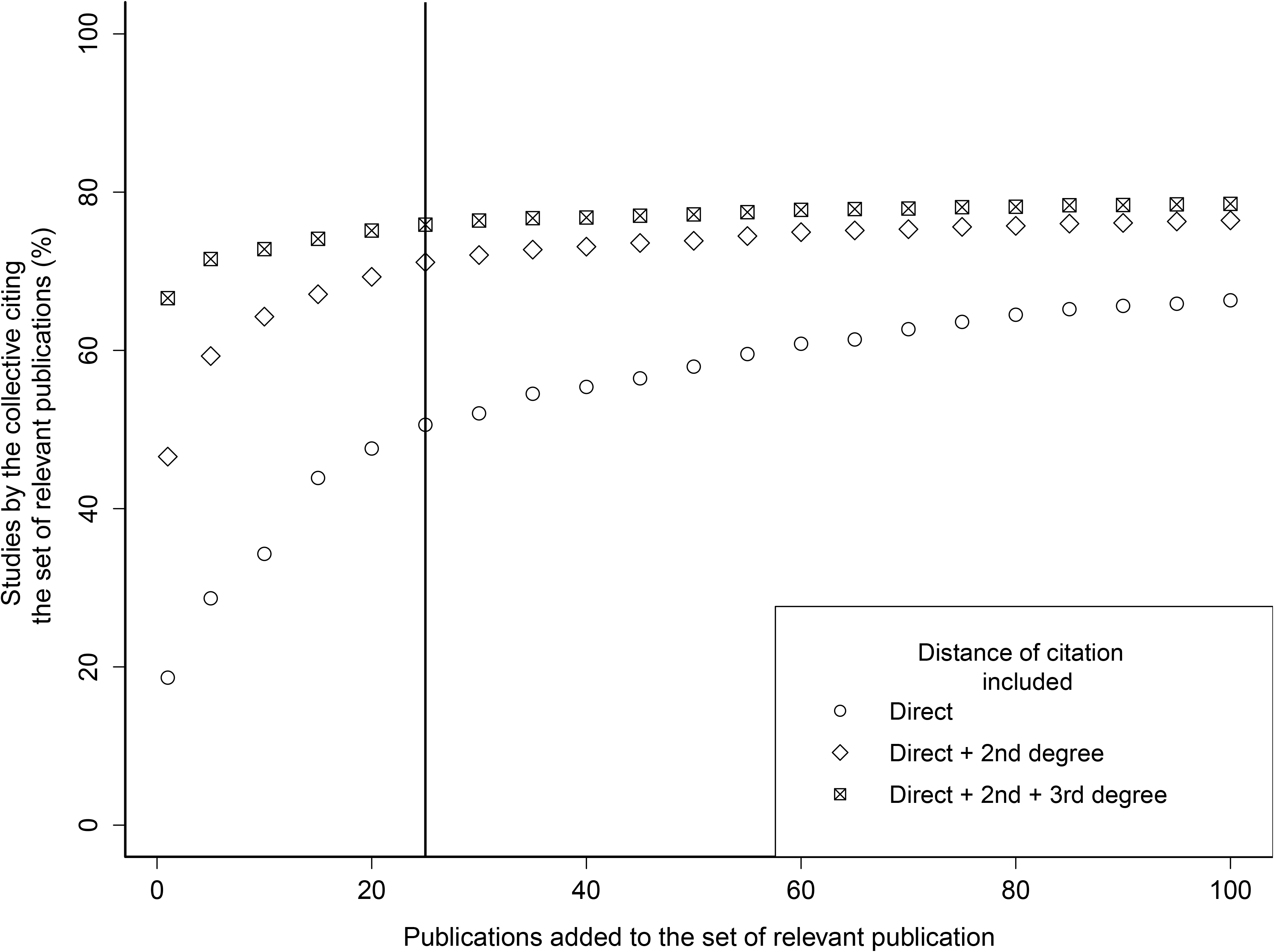
Stabilization of the relationship between the size of the relevant publication set and the percentage of studies by the selected collective citing the set. Stabilization of the relationship between the size of the relevant publication set and the percentage of studies by the selected collective citing the set. Direct citation distance considers only studies that cite the relevant publication directly. 2nd degree citations consider studies that directly cited the relevant publication and also studies that cited the studies that cited the relevant publications directly, the same logic applies to the 3rd degree citation. The vertical line indicates the threshold above which the percentage of studies of the collective that cite the set of relevant publications tends to stabilize, meaning that the addition of more relevant publications to the set does not aggregate more citations from the collective.

We checked if this fraction (20%) of the studies could be hiding a subcommunity cohesively citing another set of relevant publications, not related to the ones most cited by the 80% of the collective. We created, then, a second network containing only those 20% publications that did not cite the 100 most cited in the first network, amounting to a total of 954 publications. Afterwards, we executed the same procedure to identify the most cited publications by these 20% publications. The most cited publications in this smaller network were cited by only 6% of the 954 publications network. This result indicated that this fraction of 20% of the studies sampled at the previous step that does not cite the 100 most cited publications is not forming a divergent subcommunity that has an alternative cohesive view about succession. Most likely, these publications are individually referring to lesser-known models in the field of study without any convergence among them. Although we may technically consider these 20% as part of the collective of agents studying succession, we cannot guarantee that their views about succession are contemplated by the 25 relevant publications selected, since they did not cite any of them. At the same time, because these studies do not refer to a well-defined alternative view of succession, it does not seem that including their views in the analysis will be useful to understand major tendencies of the focal collective of agents.

The results of this step indicate that a representative fraction of the publications reporting the scientific activities of the focal collective refers to at least one proposition made in the 25 most cited publications (Table 1). We adopted, thus, the 25 most cited publications as the ones containing the conceptual bases for the studies on ecological succession by the defined community. The next step is to identify what propositions made in these publications are used by the collective of agents as conceptual bases for studying succession.

**Table 1.** List of relevant publications used in this worksorted by number of citations

| Publication | Number of citations |
| --- | --- |
| Connell and Slatyer (1977) [37] | 780 |
| Grime (1979) [38] | 468 |
| Connell (1978) [39] | 400 |
| Odum (1969) [40] | 374 |
| Harper (1977) [41] | 343 |
| Pickett and White (1985) [42] | 306 |
| Grubb (1977) [43] | 294 |
| Tilman (1988) [44] | 279 |
| Egler (1954) [45] | 271 |
| Cowles (1899) [46] | 266 |
| MacArthur and Wilson (1963) [47] | 264 |
| Bazzaz (1979) [48] | 207 |
| Huston (1979) [49] | 203 |
| Huston and Smith (1987) [50] | 200 |
| Grime (1977) [51] | 198 |
| Hubbell (2001) [52] | 189 |
| Pickett et al. (1987) [53] | 183 |
| Tilman (1987) [54] | 178 |
| Watt (1947) [55] | 178 |
| Huston and DeAngelis (1994) [56] | 177 |
| Grime (1988) [57] | 176 |
| Noble and Slatyer (1980) [58] | 171 |
| Tilman (1985) [59] | 163 |
| Shugart et al. (1984) [60] | 160 |
| Guariguata and Ostertag (2001) [61] | 160 |

### Excerpting

In this step, we selected a focal relevant publication and identified all studies of the collective of agents that cited it. Then, we randomly selected one of these studies and searched for the excerpts of text in which the focal relevant publication was cited. We disregarded excerpts in which the relevant publication was cited along with other publications, to ensure that the excerpt of text was specifically referring to the content of the focal relevant publication. We stopped the search once we reached 50 excerpts of text that fitted the criteria. Some of the sampled citing studies had more than one excerpt of text that fitted the criteria and others had none. We did this procedure to all 25 relevant publications, meaning that, at the end of this step, we had 50 excerpts of text for each of the 25 relevant publications. In a pilot analysis, participants declared that 50 excerpts of text were enough to execute the next step (content analysis) efficiently.

### Analyzing the content of citations

The 50 excerpts of text citing each relevant publication from the previous step were then submitted to a content analysis aimed to identify the main reasons for each relevant publication to be cited. This analysis was carried out by asking the graduate students and researchers in ecology (thereafter ‘ecologists’) to propose what are the main statements made in the relevant publications by only reading the excerpts of citation text. We used this approach due to the understanding that the interpretation of a single person creating such a synthesis might introduce biases in the final result. To deal with this issue, we used an approach relying on intersubjectivity as a key characteristic of the social processes of building scientific knowledge [62].

For each set of 50 excerpts citing each relevant publication, the participants were then asked to synthesize these statements in a concise, self-consistent set of propositions about the ecological phenomenon at stake. These sets were taken as the models from each relevant publication in use by the research collective in order to learn about succession.

The main characteristic of this approach is that more than one individual synthesized the models from the same excerpts and their results were compared in order to reach a consensus. Each set of 50 excerpts of text citing a single relevant publication was presented to three different ecologists, who formed a trio. Each trio was responsible for a set of three different relevant publications and synthesized the main contribution(s) of each. Therefore, to execute the synthesis using all 25 relevant publications we counted with the aid of 25 participants. All the 25 participants had a degree in biology, among which eight had a PhD in ecology (two professors and six post-doctoral fellows of the department of ecology in the university where the study was carried out) and 17 were graduate students (six PhD candidates, and 11 master’s degree candidates in ecology from the same university). The participants had different levels of familiarity with the set of relevant publications, ranging from knowing, citing, discussing and teaching about the publication to no familiarity at all. Nevertheless, at the moment of the analysis, the participants were not informed about what publication the set of excerpts of text were referring to (the actual citations in the excerpts of text were replaced by a “[Relevant Publication]” tag, in order to inform where in the sentence the relevant publication was referred to).

Even though we excluded the publications that were obviously cited by some statistical support, the participants in the content analysis could find citations that did not establish a conceptual link between relevant publication and the citing study. The citation might be just referring to a tool or method (in which case it was named ‘operational’), it might not be used to structure an argument (‘perfunctory’), or it might be used as an example of something wrong (‘negational’) [31]. The participants in our study were asked to disregard citations that were considered by them as operational, perfunctory or negational. Some of these three excluded categories of citation could help to identify the main contribution of a relevant publication. However, if a proposition is frequently cited in a domain, for example, in a negational way, it is unlikely that it is being used as a conceptual basis to create a model to learn about the phenomenon of interest.

After each participant of a trio had synthesized textually what they thought to be the model or models in use from each relevant publication, the three syntheses were combined in a consensus activity adapted from the “Nominal Group Technique (NGT)” [63]. The goal of this activity was to synthesize a verbal model or set of models that the three participants agreed upon as being what the citations of a relevant publication were referring to (see S2 Appendix for methodological details of this step). For the 25 relevant publications, 29 textual models about the natural world were identified. The following text is an example of a model synthesized from the citations of the most cited relevant publication [37].

> The mechanisms of succession are interactions between individuals who colonize the environment first with those who colonize the environment after. These mechanisms are of three types: (i) facilitation: in which species that colonize a place modify the environment increasing the chances of colonization by other species; (ii) tolerance: where species that colonize a site do not affect the chances of establishing other species; (iii) inhibition: in which species that colonize a site modify the environment reducing the chances of colonization by other species. The relative importance of these mechanisms may vary over time due to changes in environmental conditions. The general functioning of these mechanisms is of a priority effect: the chances of colonization of a site by a species are affected by the species that colonized before this site. The model does not consider the routes and mechanisms of the arrival of the initial species (i.e., why that initial species is the initial species and not another).

All models in use that were identified are shown in S1 Appendix.

## Results

We present this case study using ecological succession to illustrate how the results obtained through PATh can help us better understand ecology as a field of study. Through PATh we were able to identify 29 verbal models about succession that were frequently and widely referred to in studies about ecological succession in the past decade. We can call them the “central models”. The reason why each of these models is frequently referred to while attempting to learn about ecological succession may reveal much about the theory of this collective of scientists about this phenomenon. The conclusions we drew from analyzing the set of models in use should illustrate the kind of information one can obtain by using PATh to produce a theoretical synthesis.

### Defining ecological succession

The models we identified make different statements about what ecological succession is and focus on different properties of the phenomenon. We are not interested, however, in particular views within the domain, but in how we can define the phenomenon of succession in a way that encompasses any of the models retrieved by PATh. To do so we analyzed a single description of succession coming from one of the central models and tried to identify the necessary and sufficient conditions for succession to occur, according to this description. Then, we analyzed a second model and checked if the conditions identified for the first description of succession were also necessary and sufficient for succession as described in the second central model. If not, we tried to describe the conditions in more general terms in a way they could also be considered necessary and sufficient for succession as described in the second central model. If this was not possible, we discarded that condition as being too specific for the general description of succession. We did this until all descriptions of succession found in the central models were analyzed. We reached two propositions [25] that informed the conditions that seemed necessary and sufficient for any phenomenon of succession described in the set of central models to occur. However, some of the central models describe ecological processes that are different in many aspects. We then analyzed what was the primordial cause of the differences among these processes and conceived a third proposition that can explain why succession is such a diverse phenomenon, which may manifest in different ways.

The propositions expressing the necessary and sufficient conditions for delimiting the phenomenon of succession, as viewed in the set of models identified through PATh, are:

#### Proposition 1

At every moment in time, there is the possibility that resources will be available for use.

#### Proposition 2

Organisms from different species or at different ontogenetic stages have different probabilities of taking a fraction of the total available resource units. This difference can be due to (a) differential probabilities of site colonization, or (b) different probabilities that the individuals at the site or their propagules will take resource units.

#### Proposition 3

The dynamics of the resource and the probabilities of the species taking resource units are contingent on the abundance of species in the community and other environmental settings where the communities are changing.

Assuming that propositions 1 and 2 are necessary and sufficient for succession to occur means that in any situation in which these conditions are true, succession can occur, and if succession is occurring the two propositions hold. Even though the central models themselves may not have been conceived with these propositions explicitly in sight, our analyses show retrospectively that they underlie the construction of models within this field of study and, also, that by adopting them as true one could have conceived these models. Therefore, these propositions can be seen as fundamental principles of the domain. This means that any model that assumes these propositions as true is akin to the identified models and should be considered as a model within the domain of ecological succession as delimited by us. This information is not only useful to understand the field of study in which PATh was applied. It helps ecologists decide if their phenomenon of study is succession or not, according to the focal collective considered in this study. This clarity can be used to avoid spurious debate, and can also reveal more straightforward ways to propose changes to how we conceive the phenomenon.

### Neutral models in succession theory

Five of the 29 identified models did not include the word “succession” in their description. Three of these models came from references about the competition-colonization trade-off and its relationship to disturbance regimes (Models 4, 16, and 22, described originally by Connell [39]; Huston [49]; Watt [55], respectively. See S1 Appendix). In the same vein, the two other models that did not mention succession explicitly were the models of Island Biogeography and the Neutral Theory, which describe the role of colonization and extinction in community assembly in general (Models 14 and 19, described by MacArthur and Wilson [47]; and Hubbell [52]. See S1 Appendix).

Even though these five cited relevant publications do not focus on describing or explaining the phenomenon of succession per se, they describe important concepts that are currently being used to make new propositions about what succession is and how it works. Considering that the domain of succession has been grounded in niche theory for many years [7], it is somewhat surprising that Hubbell’s book [52] was the 16^th^ most referred by the collective studying the phenomenon currently (being, in fact, more referred than some papers considered as classical references for studies on succession). An overview of the citing excerpts reveals that the neutral model is frequently used to explain successional patterns at landscape or global scales, meaning that this model is actually being used to learn about succession. Similarly, the model of island biogeography is frequently used to explain why different patterns of succession emerge in fragments at varying distances from other fragments (see S3 Appendix). The fact that models assuming niche differentiation [37, 38] and models assuming competitive equivalence [47, 52] are being combined to learn about ecological succession, despite the incompatibility of such assumptions, strengthens our argument that theoretical syntheses based on axioms would not suffice to make an adequate description of this field of study [19]. An axiomatic synthesis of this field would have to either enunciate axioms that are not compatible with one another, something that would contradict the definition of axioms, or deliberately disregard some models that are in fact being used, neglecting how scientific research is actually conducted. This could lead to discussions about the factuality of each contradicting axiom, as it has happened for these two kinds of models in the past, when we have evidence that both seem useful to learn about succession [64–66].

### Classical and contemporary views on succession

We also observed that the division between “classical” and “contemporary” views on succession is not as clear for this collective as it is depicted in some publications, including textbooks [33, 67–69]. Concepts classified as being from the classical view in these studies are still used today to develop new models and concepts classified as contemporary seems to be used in the present less frequently than one would expect. For example, in the “contemporary” view succession is regarded to be individualistic, while in the “classical” view succession is treated as supra-individualistic.

Individualistic succession means that succession is essentially the exchange of individuals, in the sense that any pattern observed is just the result of interactions among these individuals. Supra-individualistic succession assumes that the agents of succession can be communities, functional groups or other supra-individual entities, and, therefore, succession can occur as these entities change in time [70]. The high frequency of citation of models considering supra-individual entities as the agents of succession (models 6 [40] and 29 [61]) shows that such models of succession are used in the present, and not just rarely.

Odum’s article [40] was the 5th most cited publication in a network of more than five thousand items representing the scientific activity of the focal collective. This paper is frequently cited in order to support the idea that succession is the change in ecosystem properties through time. Hence, there can be little doubt that the scientific collective studying succession still adopts supra-individual approaches to this phenomenon as a possible way to understand it. Similarly, the model proposed by Guariguata and Ostertag [61] is used to give support to the claim that succession can be viewed as changes in functional groups, which are also supra-individual entities.

In the classical view, succession has been considered a dynamic intrinsic to communities of primary producers, while in the contemporary view succession it is considered a dynamic of communities at any trophic levels [33, 68]. What our findings show is that the most referred models in this field are still about succession of plant communities. Most models are explicit about it and only a few propose mechanisms that could be applied to heterotrophic organisms. Therefore, it seems that succession ecology is still focused on plants and if there is a change towards a more multitrophic understanding of succession, this change is yet not noticeable enough to have surfaced with the most referred models.

In synthesis, some models classified as belonging to the “classical view” have been still used often as a conceptual basis for learning more about succession in the last decade, while some others classified as belonging to the “contemporary view” have not been so frequently used. This result leads to the question of whether this classification of concepts as belonging to classical and contemporary views on succession is indeed a description of how the collective understands the theoretical development of the field or is rather some sort of rational reconstruction found in textbooks that does not really correspond to the way the collective pragmatically uses models, or, yet, a prescription of how the field should be seen. These observations highlight some insights that PATh allowed us to reach, in this case concerning the theoretical structure of the succession domain.

## Discussion

We showed that PATh allows for a theoretical synthesis that assesses the conceptual bases of a field of study by describing the views of a defined community about an ecological phenomenon. Therefore, such a synthesis does fulfill one of the roles of a scientific theory, namely, the description of conceptual bases [1]. However, the way in which PATh allows for such a synthesis gives it some distinctive characteristics.

The first distinctive feature is that a synthesis made by PATh has an explicitly descriptive character, in the sense that this synthesis intends to provide a summary description of how scientists are employing the knowledge available to them. This description arises, thus, from a pragmatic perspective on theories and models.

Pickett, Meiners and Cadenasso [33] provided a literature overview and a general theoretical synthesis for ecological succession that can be compared with the one produced by using PATh. They also mention that this synthesis is to be used as a mechanistic reference in future studies of succession. They started from published papers to propose a synthesis around fundamental propositions about ecological succession and then described some central models in the field, just like we did. Some of their propositions are in accordance with the ones derived from PATh in some respects, but in disagreement with others. It is important to note, however, that their synthesis was made by using an “expert opinion” approach which is highly dependent on the proponents’ own views and in which it is not clear what procedures were carried out to reach the set of propositions and models. These procedures are important because without knowing them it is not possible to be critical about the results and methodological criticism is one of the cornerstones of the construction of scientific knowledge [71].

The main difference of using PATh to produce theoretical syntheses instead of expert opinion is that a detailed methodological description can be made to explain how propositions and models are directly linked to the actual scientific research made in a field. This process allows for a more direct and efficient process of criticism and re-evaluation of the synthesis. For example, one might argue that the third step was not conducted properly, resulting in a biased survey of the publications reflecting the activities of the focal collective of scientists. This could entail a set of relevant publications that do not contain conceptually important propositions about the phenomenon. This statement could then be tested, first, by evaluating if the parameters of the search may result in biased outcomes and then adjusting them to avoid the detected biases. Finally, we could check if the final set of central models obtained was in fact different from the one resulting from the previous search.

The second distinctive characteristic of syntheses made with PATh is related to the assumption that a theory about a phenomenon can be seen as the views of a specific collective of scientists about a specific phenomenon or class of phenomena. This assumption implies that different collectives of scientists studying a phenomenon may have different theories about it. Because the definition of who is part of this collective is a methodological step in the PATh, multiple instances of the approach can be applied to different collectives and the theoretical syntheses resulting from these different instances can be compared (or, perhaps, even combined to reach a more overarching approach). For example, an application of the PATh to different time scopes can reveal how some change may have happened through disuse of models that were highly cited in the past and/or increase in the use of models that are new to the field. Scientific practices can also change from place to place [72, 73], as well as depending on gender [74], age, academic position and other social and cultural characteristics [75]. Different applications of PATh could be used to answer if these different time, geographic and demographic profiles could lead to different views about a specific phenomenon. These social aspects of scientific work have been neglected for many decades in the natural sciences [76], and the use of the PATh may offer, therefore, a tool for dealing with these aspects that may raise the interest of academics to consider them more closely.

The syntheses allowed by the PATh are explicitly descriptive in the sense that the enunciation of the central models identified is not a recommendation of the best models to learn about the focal phenomenon. These models merely summarize the conceptual basis of research conducted in the field. That, however, does not preclude syntheses produced by the PATh to be used to guide scientific activity in the field, although in a non-axiomatic way [19]. For example, we detected that many scientists in the field of ecological succession accept that succession can be modelled as a supra-individualistic process, while others propose that succession be understood as an individualistic process [33]. This conflict revealed by the produced synthesis can alert either side to the need to express their viewpoints more specifically, either by presenting evidence supporting the claim that we should abandon models assuming supra-individualistic views of succession or arguing why there is no reason for that. If this knowledge is not yet available, it might be the case to invest in research to resolve this dispute. Such conduct should help avoid spurious debates within the field and, as a consequence, make the field more efficient in generating knowledge about the phenomenon. Therefore, we can use the information obtained by accessing and analyzing the views of a collective of scientists about a phenomenon using the PATh to make more informed decisions about what needs to change and how it needs to change. This change is assumed to be an integral part of theory development.

In the axiomatic view of theory, a scientist knows exactly the assumptions their models need to have in order to pertain to a given theory. If the model does not assume the axioms as true, it is not part of that theory [12]. Therefore, a theory is an *a priori* static entity defined by a set of axioms. If there are grounds to believe that the axioms are not true of the world, the entire theory must be cast aside. A synthesis made by using the PATh will show, in turn, which are the central models under use in a field of study and, therefore, this theoretical synthesis is an *a posteriori* entity. This is a considerable change of perspective because there is more space to tinker with theories and models, merge them, reinterpret them, and so forth. If one assumes that the conceptual basis of a field is grounded on a collective view of a collective and that collective change, it will become practically a necessity to assume that theories are ever-changing entities [30].

## Conclusion

Ecologists have shown that they can produce knowledge about the world without the restraints and guidance of a single, axiomatic theory. Some even disrecommended pursuing systematization of knowledge into theories on the grounds that theories are reins that restrain scientific development [77]. Meanwhile, the number of studies proposing conceptual unification, disambiguation of concepts, conceptual cleaning and calls to training more theoreticians in ecology [5, 78–82] indicates that there is a demand for some kind of systematized way to represent the knowledge generated in ecology. Here we presented an approach that could help ecologists create theoretical syntheses without the necessity of identifying axioms and fitting knowledge development into some predefined structure. These syntheses can be used to make more informed decisions about how to approach a phenomenon, for example, by deciding to investigate more thoroughly a model that is gaining attention quickly, by investing in propositions that are being neglected by a community or by advising against the use of models that have been shown to be flawed.

To the extent that the relevance of models within a field can be gauged by how much they are used by a collective of scientists, the relevance of a theoretical synthesis can also be gauged by how much a collective of scientists uses it. If ecologists do in fact use the syntheses made by using the PATh to guide their scientific activities, these syntheses will fulfill both roles of a clearly enunciated theory, namely, describing and guiding knowledge generation [1]. Furthermore, since these syntheses are made *a posteriori* their guidance will work much more like consulting maps than restraining reins.

## Supporting information

**S1 Appendix. Central models** Verbal models synthesized for each relevant publication.

**S2 Appendix. Description of consensus technique** Detailed description of how the consensus technique was carried out to reach the central models.

**S3 Appendix. Excerpts of text citing relevant publications** All excerpts containing the multiple citations for each relevant publication and that were subject to content analysis in order to make the model syntheses.

## Acknowledgments

We thank all 22 non-author participants that volunteered to the group-analysis (A. Palaoro, C. Hohlenwerger, D. Muniz, D. Bertuol, F. E. Mendes, F. D’Albertas, G. Bispo, G. Pitta, I. Romitelli, J. Menezes, L. Carneiro, L. Souza, L. Teixeira, M. Leite, N. H. Azevedo, R. Ourofino, R. Pelinson, R. Leporoni, R. Quesada, S. Iop, S. Mortara, V. Caldart). We also thank D. Scarpa and M.E. Prestes for their comments on the initial version of this manuscript.

BTB was supported by scholarships from CAPES (grant number 88882.315632/2019-01) during the time of this study. CNPq (grant number 465767/2014-1) and CAPES (grant number 23038.000776/2017-54) supported INCT IN-TREE, which the current study is part of. CE and PP received research fellowships from CNPq (grant number, and 311051/2018-9, respectively). CE was also funded by CAPES and UFBA for Senior Visiting Researcher Grant included in the CAPES-PRINT Program, which funds his stay in the Centre for Social Studies, University of Coimbra, Portugal (grant number 88887.465540/2019-00).

