## Supplementary material for "A pragmatic approach to produce theoretical syntheses in ecology": S1 Appendix

### 1 Supplementary material I – Identified models

The Models described here are direct conclusions of the contributors in step 4 of the PATh. As described in the methods, each description was made by a different trio contributors.

Model 1 - Described in Connell and Slatyer (1977)

The mechanisms of succession are interactions between individuals who colonize the environment first with those who colonize the environment after. These mechanisms are of three types: (i) facilitation: in which species that colonize a place modify the environment increasing the chances of colonization by other species; (ii) tolerance: where species that colonize a site do not affect the chances of establishing other species; (iii) inhibition: in which species that colonize a site modify the environment reducing the chances of colonization by other species. The relative importance of these mechanisms may vary over time due to changes in environmental conditions. The general functioning of these mechanisms is of a priority effect: the chances of colonization of a site by a species are affected by the species that colonized before this site. The model does not consider the routes and mechanisms of the arrival of the initial species (i. e. why that initial species is the initial species and not another).

Model 2 – Described in Grime (1979)

The process of ecological succession involves a change in the relative dominance of each type of ecological strategy of plants over time to a point of equilibrium, being determined by the biotic and abiotic conditions of the initial environment. The three main types of ecological strategies of plants are competitive, ruderal and stress-tolerant. These types of ecological strategies are generated by the trade-offs of investing in competition or survival and investing in growth or competitive ability. In an environmental gradient between productivity and stress, given enough time for the community to reach equilibrium, good competing species dominate at the productivity end and stress-tolerant species dominate at the extreme of stress. This generates a greater diversity in the mean conditions of the gradient, where the two types of species occur. Stress and disturbance are different mechanisms that influence the diversity of species in the community, respectively limiting the rate of productivity or removing the biomass of plants in the environment.

Model 3 – Described in Grime (1979)

Disturbances that remove biomass allow the arrival of ruderal species, delaying the effect of competitive exclusion and, therefore, the time to reach the balance. In situations with low disturbance frequency, there is a tendency for competitive exclusion and few species to dominate. In situations with a high frequency of disturbance, ruderal colonizing species would dominate. In situations with an intermediate frequency of disturbances, there is a mixture of species of the different strategies, and therefore, greater diversity is expected.

Model 4 – Described in Connell (1978)

Environments with intermediate frequencies/intensities of disturbance should present greater diversity in relation to environments with low and high frequencies/intensity of disturbance. This is explained by a trade-off between competitive skill and colonization. Environments with intense and/or frequent disturbances are dominated by species that are good colonizers (therefore, bad competitors). When there is little disturbance, the environment is dominated by the good competitors, who exclude the colonizers. Therefore, in intermediate levels of disturbance, there will be both types of species and the wealth will be maximum. Thus, at intermediate frequencies/intensities of disturbance, there would be the coexistence of pioneer species with species of the climax community.

Model 5 – Described in Connell (1978)

Species with good post-disturbance colonization capability are different from the efficient competing species. Therefore, species diversity will be low in the early stages of succession, when pioneers dominate. Diversity will have its maximum in intermediate stages when there are still pioneers but the secondary and late species are already beginning to establish themselves, and diversity will again fall into the final stages when the pioneers are eliminated.

Model 6 – Described in Odum (1969)

The model proposed in this publication aims to identify predictive patterns of forest ecosystem development (ecological succession), based on physical/chemical characteristics (e.g. soil nutrients, CO stock) and forest biological characteristics (e.g. species composition, biomass) as the forest matures, also underscoring the bidirectional interaction between the physical and chemical parts of an ecosystem.

The model proposes a directional process that begins in an event of environmental disturbance and reaches a state of equilibrium over time. Operational variables of the model can be gross primary production, respiration, species diversity (and consequently the change in alpha diversity), carbon storage, the net rate of ecosystem productivity.

The model predicts that in the initial stages communities are dominated by rapidly growing colonizing species, whereas in the late stages the communities are dominated by competing species, whose growth is slow. Diversity increases throughout the succession by migration but decreases over time through competition.

Regarding the physicochemical characteristics, the model also predicts a directional trajectory in which the initial stages have a lower carbon stock and higher primary production due to the high respiration rates of the colonizing species, whereas late successional stages reach a maximum stock of carbon and primary production.

Therefore, there must be a gradual accumulation of species throughout the succession process, culminating in a stable community in terms of richness/composition and also in primary productivity. On the other hand, resilience to disturbances in late communities tends to be lower than in early communities, since in late communities the species are highly specialized, and in early communities, productivity is maximized.

Model 7 – Described in Harper (1977)

The colonization of an area by seedlings is the most sensitive stage of the plant life cycle (because it is the stage most subject to biotic competition) and depends on the availability of safe sites, that are sites with soil conditions or opportunities suitable for seedling recruitment of the existing seed bank (or propagules bank), which are all viable seeds (both dormant and active) contained in the soil of an area.

Model 8 – Described in Pickett and White (1985)

The natural disturbances ("any discrete event in time that disrupts the ecosystem, community or population structure and changes resources, substrate availability, or the physical environment") generate environmental heterogeneity, mainly by modifying the availability of limiting resources, favouring reorganization of the community in structure and species composition. With this, communities are prevented from reaching a stable state, called by the author of climax. The effects of disturbances can be measured by loss of biomass, structure, function, and species richness. Ecosystem responses to disturbances will depend on the quality and quantity of abiotic resources that influence productivity, land use history and life characteristics of species (such as competitive ability). The Patch Dynamics Theory explains how the greater diversity of plants occurs in intermediate disturbance regimes, which create species patches in distinct successional stages within the same community. By keeping communities in transient states, disturbances also favour communities in different successional stages coexisting on a larger spatial scale.

Model 9 – Described in Grubb (1977)

The niche of species changes with the stage of life. The regeneration niche (i.e. the biotic and abiotic requirement to replace an adult individual) differs among them, and given the micro-scale environmental heterogeneity, favours coexistence between species. It is also the niche of regeneration that defines the spatial distribution of adult individuals, so that frequent disturbances, altering the biotic and abiotic conditions, alter which species will be established in relation to the already established species, modifying the successional trajectory.

Model 10 – Described in Tilman (1988)

The publication is about the dynamics between environmental factors and ecological succession/plant development. In this publication, the resource ratio hypothesis is proposed, which stipulates the effect of the existing resources in a site on the plant community and the inter-specific competition. In primary stages of succession, light is available but few resources in the soil, which causes intense underground competition. In later stages of succession in which accumulation of nutrients occurred, the competition is for luminosity. Plants that are good acquires of resource in poor soils (colonizing plants) invest more in roots and are bad competitors in environments with abundant resources.

Model 11 – Described in Egler (1954)

The model states that initial conditions and stochastic processes guide the course of succession. The model says that colonization is by early and late species simultaneously. Both pioneers and late remain throughout the succession and their dominance occur at different times. Stochastic processes act mainly in the late stages of succession when biomass is concentrated in fewer individuals. This model goes against the Gradual Succession model, in which species are replaced by others along the succession.

Model 12 – Described in Cowles (1899)

Throughout the succession, changes in species composition can alter soil conditions, promoting the establishment of different species. This process leads to the reduction of annual and increase of grasses and especially woody, and the increase of the richness of plant species.

Model 13 – Described in Cowles (1899)

The species that form dunes are grasses, shrubs or trees. However, succession in the dunes leads to a progressive and directional change of grasses to the mesophytic forest, through changes in luminosity and soil caused by each successive dominant species (which facilitates the establishment of different species). However, the successional trajectory depends on the topographic location in the dunes.

Model 14 – Descrito em MacArthur and Wilson (1963)

In islands, the balance between rates of extinction and immigration determines the richness of species. Extinction rates are higher the lower the area of the island, while immigration rates are higher the lower their isolation from the mainland. Thus, the richness of species on islands is positively influenced by the size of the island and negatively by its isolation. Fragments of habitats surrounded by hostile matrices function ecologically as islands and are therefore also affected by the above processes.

Model 15 – Described in Bazzaz (1979)

This model assumes that the life history traits of vascular plant species change as the successional stage progresses. These traits may be the ability to tolerate light, competitive and dispersive ability, defence against herbivores, growth time and lifespan.

The model predicts that different successional stages should present communities with different ecophysiological properties and/or strategies to maximize resource use. When light is limiting, differentiation in light capture strategy is an important mechanism for successional changes and species coexistence.

Pioneer plants are inferior competitors that need to grow and consume resources quickly to disperse continuously to new open habitats, and thus their size is smaller and lifespan is shorter to that of late stage plants, although their dispersal capacity is superior. Late stage plants, on the other hand, are more efficient at using lower light intensities, but they do so by lower photosynthetic rates. As the succession progresses, the forest community increases both in the number of herbivorous defence structures and in biomass and density.

Model 16 – Described in Huston (1979)

The model states that the intensity of the disturbance and the competitive exclusion are fundamental processes to explain the diversity of species. In high productivity environments, dominant species will tend to exclude lower species, leading to low diversity and dominance of long-lived species. On the other hand, intense or recurrent disturbances may reduce competitive exclusion, since species of rapid growth and rapid recolonization will be favoured.

Once:

1. Competitive exclusion is a process that reduces diversity,
2. Competitive exclusion is more likely to occur in stable and productive environments (later stages of succession) and
3. Intense disturbances reduce diversity by eliminating late species incapable of recolonization.

Thus, intermediate disturbances should promote greater diversity of species than extreme disturbances or absence of disturbance. Therefore, a dynamic equilibrium must occur in which inferior competitors will be able to persist after a disturbance until resource levels increase or decrease, and then be excluded by higher competitors as the succession approaches the climax.

Model 17 – Described in Huston and Smith (1987)

The succession – sequential change in relative abundance of dominant species – is governed by trade-offs between different characteristics. Better dispersing or breeding species (i.e. with high growth rates, maturing earlier or with shorter generations) are adapted to early stages (after disturbance) and are replaced by better competing species, adapted to the conditions created by previous species (e.g. low light). Thus, species substitution is faster in the earliest stages and competition increases throughout the succession.

Model 18 – Described in Grime (1977)

In plants, there is a "three-way trade-off" between being good at competing for resources, resisting disturbance, or tolerating stress. This trade-off establishes three functional types of plants: ruderal, with high productivity, high dispersion/colonization and low competitive capacity; competitive, with high productivity, low dispersion and high competitive capacity and; resistant, with low productivity and high tolerance to unfavourable conditions. Throughout the successional process, environmental conditions change and therefore there is a substitution in the type of plant favoured, as it favours different functional types of plants.

Model 19 – described in Hubbell (2001)

In the neutral model, the species are all demographically equivalent, that is, they have equal chances of dying, reproducing and colonizing a place. In this context, the attributes (traits) of the species are not important, and therefore have no relation to their success or abundance in a community. Thus, the variations of abundance in time and space are stochastic processes defined by the rates of dispersion, speciation and extinction. Therefore, limitation to the dispersion and initial abundance of each species in the local and the landscape are relevant for the assembly of the communities and therefore also for the succession.

Model 20 – Described in Pickett et al. (1987)

There are three general causes that determine a succession: species availability, space availability, and species performance. That is, for colonization by a species to occur, there must be propagules of this species, a place where they can settle, and the species must have a performance that guarantees its survival for a certain time.

Model 21 – Described in Pickett et al. (1987)

Disturbances are important causes of changes in ecological succession. Tolerance, facilitation and inhibition can act in ecological succession at the same time by the same species.

Model 22 – Described in Watt (1947)

Plant communities are dynamic, in the sense that species composition changes over time. These changes are governed by disruption and competitive exclusion. Competitive exclusion occurs when there is a high availability of resources (or productivity) which causes low diversity (in the long term) because the most competitive species exclude all others. Recurrent disturbances can prevent competitive exclusion by the fact that there is a trade-off between recolonization capacity and competitive capacity. In rich/productive environments plant diversity will be maximal at intermediate levels of disturbance. In turn, high plant diversity favours high animal diversity (or at least phytophagous insects).

---

Model 23 – Described in Tilman (1987)

Nutrient limitation shapes the structure of plant communities via the competition process. Both very limited quantities and nitrogen enrichment can lead to reduced diversity due to competitive exclusions. Pioneer species are competitively superior in nitrogen-poor environments, while later-stage succession plants are competitively superior in nitrogen-rich environments. There is an interaction between constraining resources shaping the community structure. For example, prairies are limited in nitrogen in wet years and by water in dry years.

---

Model 24 – Described in Huston and DeAngelis (1994)

Diversity patterns vary widely and are influenced by environmental gradients and the life history of the organisms involved. Environmental gradients may be regulators (which directly influence plant growth, such as precipitation and temperature) or complexes (which influence other resources and regulatory gradients, such as altitude). Regarding life history, differences in environmental diversity can be related to the competitive capacity of species (which determines the competitive exclusion rates). Disturbance levels also affect diversity: localized disturbances appear to increase species richness and maintain ecosystem resilience by redistributing resources and reducing competition, while very intense disturbances tend to decrease richness of species. In general, environments with low productivity and low competitive exclusion should have high turnover rates and high diversity, while productive environments with high competitive exclusion should have low turnover and, consequently, low diversity. It is important to highlight that different life forms may present different responses to the productivity gradient. On the other hand, exotic and native species seem to respond in the same way, since they present similarity of attributes.

Model 25 – Described in Grime (1988)

Plant species can be classified into three functional strategies: ruderal, stress-tolerant or competitor, or intermediaries between two of these strategies. Plant strategies are related to the availability of resources in the environment, the occurrence of disturbances and species characteristics. For this classification of strategies, a key is needed based on a broad list of relevant ecological information, such as reproductive effort, seed characteristics, and tolerance to disturbances.

Model 26 – Described in Noble and Slatyer (1980)

Ecological succession after a disturbance can be separated into two main phases: the regeneration phase, in which competition is low and abundance of individuals is determined by regenerative processes; and the competition phase, in which regeneration is over and competition is high and determines the abundance of plants. During the regeneration phase, plant species that establish themselves first are those that: (1) have the capacity to vegetatively regrow, (2) have seed banks in the soil, or (3) have propagules that can disperse rapidly from other sources. The frequency of disturbances determines which species will be favoured throughout the process of ecological succession. Which species will be favoured for a given frequency of disturbances depends on the functional attributes of the plants (e.g. age of first sowing, seed longevity in soil seed bank, dispersion strategy, life history, etc.). The functional attributes of the plants present in these environments determine the way in which the plants colonize the environment and may have been selected by the disturbances themselves. Thus, through these functional attributes, it is possible to predict which species will resist the disturbances and settle in the regenerative phase of ecological succession.

Model 27 – Described in Tilman (1985)

The competition-based resource-ratio hypothesis (RRH) of plant succession proposes that species are efficient to use one resource, but inefficient to use another (which is your limiting resource). So, RRH predicts that a trade-off between light availability and soil nutrient shall predict the plant species community composition because plant species cannot enhance abilities to utilize both nutrients and light efficiently. Therefore, RRH determines whether competing species can coexist and, if not, which species will exclude others. It also predicts: 1) the highest diversity of competing species should occur at intermediate resource availability; 2) species composition in a community should change whenever the relative availability of limiting resources changes.

Model 28 – Described in Shugart et al. (1984)

The responses of individuals to environmental conditions influence the composition and structure of the community explaining how it changes over time. These individual responses are influenced by the growth rate, allometry, mortality and recruitment of the species, by characteristics of the functional group of the species, and by competition with its neighbours. Based on these individual responses, it is possible to propose models to predict the ecological succession pathway after a disturbance. This type of model used to analyze forest successions, describing the growth and mortality of tree individuals and/or simulating tree responses to the environment, climate and disturbances are called a gap-model.

Model 29 – Described in Guariguata and Ostertag (2001)

In the ecological succession of secondary rainforests, there is a sequence of phases characterized by the type of plants found in the area: A) colonization by herbaceous and shrubby plants; B) substitution by pioneer plants intolerant to shade; C) fast-growing and higher and long-lived plants are established and make up the bulk of the biomass. Throughout these phases, other changes also occur: 1) an initial investment focused on the acquisition of resources is replaced by an investment in structural materials; 2) biomass is progressively increasing; 3) the basal area increases more gradually than the vegetation cover; 4) taxonomic and functional diversity increase; 5) increase in the number of species with larger individuals, leading to forest stratification.
