## Supplementary material for "A pragmatic approach to produce theoretical syntheses in ecology": S2 Appendix

### 2 Supplementary material II – Application of the consensus group technique

The application of the adapted NGT proceeded as follows:

1. The proposed synthesis of each person of the trio was presented to the entire trio, without each person knowing the authors of the synthesis, except their own proposed synthesis. The proposed synthesis was the description of how many and which were the most relevant models described in the focal relevant publication. Each model is then numbered.

2. After reading the proposition of their peers, each person in the trio reevaluates their own proposed synthesis and make a new proposal. This new proposal is solely the combination of which of the numbered models should be used to synthesize the relevant models of the focal relevant publication. The participants in the activity should consider that some of the models presented in the first round might be combined in a single model, others might be kept separated (if one thinks the literature is referring to more than one model in this publication), and other might be discarded as non-relevant mention in the literature.

3. The combination proposals are then presented to the trio, again without each person knowing the author of each combination proposal, except by their own.

4. The combination proposals are then ranked by the participants in a scale from the most adequate to least adequate according to their individual views. The combination proposal with the higher rank considering the ranking of all three participants is selected. If the combination proposal states that the literature refers to the focal relevant publication by more than one model, the next rounds are applied to each individual model.

5. The model or models are selected to be synthesized textually for each one of the participants individually. This time the participants should consider only the models numbered in the first round.

6. For each model re-synthesized, the three synthesis made for each participant are presented, again without each person knowing the authors of each synthesis. Then, the newly proposed synthesis are ranked as in the previous round and the synthesis with the higher rank is selected.

7. The selected synthesis is submitted to an open evaluation by the participants that propose modifications to it if they think necessary. The modifications are compared to the synthesis selected in the $6^{th}$ turn and re-ranked.

8. The process described in the turn 6 and 7 are repeated until consensus is reached.
