## Supplementary material for "A pragmatic approach to produce theoretical syntheses in ecology": S3 Appendix

Excerpts of text citing Connel and Slatyer (1977)

A typical approach to restoration has been to plant the suite of species found in an intact community relict. However, it is possible that there are necessary but fully transient community states that occur during succession (Connel and Slatyer 1977).

This phenomenon improves micro-site conditions, which allow occupation of the same species, or others and thus influences species richness (Connel and Slatyer 1977).

Connel and Slatyer (1977) proposed three models of succession: (1) facilitation, in which early species modify the environment to make it more suitable for later colonizers; (2) tolerance, in which, as the environment changes, established species exhibit a progressive tolerance of invading species; and (3) inhibition, in which early colonizers restrict the invasion of later colonizers.

Therefore BSCs enhance the probability of colonization and survival of later successional species according to the facilitation model of Connel and Slatyer (1977).

The shift could thus be due to facilitation of N-demanding species (Connel and Slatyer 1977).

The classic facilitation, tolerance, and inhibitory models of succession proposed by Connel and Slatyer (1977), based on the net effect of early colonists on later ones, have been widely used to classify successional patterns in terrestrial and aquatic systems (Bertness 1991, De Steven 1991, Farrell 1991).

This implied that, regardless of dominant exotic species, all patch types were roughly equivalent at inhibiting native grass recruitment from added seed (Connel and Slatyer 1977).

Connel and Slatyer (1977) identied three models of succession during community development in which early successional species can have either a positive (facilitation), neutral (tolerance), or negative (inhibition) effect on the establishment of later species.

Plant-plant interactions, including succession processes (Connel and Slatyer 1977), have been shown to have a marked inuence on plant distribution and diversity patterns (Bruno et al. 2003).

This species occurs especially on steep north-facing slopes (Richard 1990) with two possible scenarios, i.e. the establishment arboreal communities or long standing, dense, competitive shrub cover (Anthelme et al. 2002), according to the inhibition model described by Connel and Slatyer (1977).

Facilitation has long been recognized to potentially operate during the course of succession (Connel and Slatyer 1977).

We neither detected strong effects of environmental variables on herb layer species diversity, nor did we nd any evidence for facilitation or inhibition among species (sensu **?**).

Three studies that have experimentally examined the net effect of BSCs on later successional vascular plant species (Chapin et al. 1994; Elmarsdottir et al. 2003; Hawkes 2004) indicate that the effect generally ts Connel and Slatyer (1977)s facilitation model, wherein the BSC enhances the probability of colonization and survival of members of later seres.

Whether or not BSCplant interactions in succession follow Connel and Slatyer (1977) facilitation or inhibition models, we should be aware of the nature of this interaction because properly manipulating it could result in more successful rehabilitation on the ground.

Large-scale species coexistence through a successional mosaic is the product of two potential mechanisms, only one of which explicitly depends upon the competition-colonization trade-off (Amarasekare 2003). First is succession through facilitation (Connel and Slatyer 1977).

According to the facilitation model of Connel and Slatyer (1977), the micro-crust should enhance the probability of colonization for bryophytes and vascular plant species.

Connel and Slatyer (1977) propusieron tres modelos para explicar la influencia de plantas pioneras sobre la invasin y crecimiento de especies ms tardas en la sucesin. Estos son: facilitacin, inhibicin y tolerancia. Especies sucesionales tempranas podran tener efectos positivos (facilitacin), negativos (inhibicin) o no tener efecto (tolerancia) sobre el establecimiento de otras especies.

En el hemisferio norte, los estudios de colonizacin de plntulas de rboles y arbustos en praderas abandonadas despus de la eliminacin del bosque (De Steven 1991a, 1991b, Gill & Marks 1991, Callaway 1992) apuntan, en su mayora, a evaluar los mecanismos de sucesin propuestos por Connel and Slatyer (1977).

At early stages, the succession of fouling communities from Baha La Herradura appears to follow at least partly the tolerance model proposed by Connel and Slatyer (1977).

As a result of this modifying activity, established vegetation can facilitate the incorporation of new individuals into the community, according to a model of succession by facilitation (Connel and Slatyer 1977).

It is also possible that once earlier colonists are established, they arrest the incorporation of new individuals according to a model of succession by inhibition (Connel and Slatyer 1977).

Two main processes of competitive interaction between plant species induced by the modication of environmental conditions are facilitation and inhibition (Connel and Slatyer 1977).

The concept of ecological succession refers to more or less deterministic (rule-based) developments in the composition or structure of an ecological community after disturbance (Connel and Slatyer 1977)

The increased stochasticity of compositional development seems not to be related to invasion niches on bare ground and subsequent colonisation events (Connel and Slatyer 1977)

It is well accepted that environmental conditions do not remain constant over succession, altering the strength and selectivity of abiotic and biotic ltering processes over time (Connel and Slatyer 1977).

At early successional stage, the strong environmental lters may initially select good dispersers or disturbance-tolerant species from several clades of closely related species (Walker & Chapin 1987; Helmus et al. 2010), and communities should show signicant clustering. However, these early colonists could modify the environment that facilitates the colonisation of other species (Connel and Slatyer 1977)

This class has received the most attention and therefore contains the greatest diversity of potential successional drivers. It covers Tilmans competition-based resource ratio hypothesis (Tilman 1985), the interaction-based views of Connel and Slatyer (1977) and even Clements (1916), as well as the majority of trait-based sorting suggested by Grime (2001).

Connel and Slatyer (1977) produced a revolutionary paper illuminating the mechanisms of succession in natural communities.

Classical succession has the key assumption that species are replaced because they modify the environment so that it becomes more suitable for other species but less suitable for themselves. This was termed the facilitation model of succession by Connel and Slatyer (1977), who also coined the term inhibition model for Eglers (1954) hypothesis of initial oristic composition that followed from Gleasons (1917) initial questioning of the view of Clements (1916). In this model, species arriving rst at any site persist by excluding or suppressing all other species, with no species competitively superior to others, so that short-lived species are replaced by longlived species. Connel and Slatyer (1977) developed a tolerance model between these two where some species are competitively superior to others and come to dominate because of a greater tolerance to limiting resources.

The model was derived from, and built upon, the successional mechanisms of facilitation, tolerance and inhibition of Connel and Slatyer (1977).

They then increased in their importance as Pinus declined, essentially following the tolerance model of Connel and Slatyer (1977).

It is worth noting that the facilitation model (Connel and Slatyer 1977) might also operate under Pinus forests in some Mediterranean ecosystems

| |
|---|
| To investigate which successional mechanism is more important, facilitation or tolerance (sensu Connel and Slatyer (1977)) |
| Facilitative interactions, together with posterior competition, are among the proposed mechanisms triggering succession Connel and Slatyer (1977) |
| Connel and Slatyer (1977) contributed to the demise of the exclusive facilitation view by proposing the alternatives of inhibition and tolerance. |
| Hence, vegetation dynamics is no longer assumed to require directionality or determinism, and is no longer seen to be driven only by the net effect of facilitation (Connel and Slatyer 1977). |
| The recognition of facilitation as a mechanism of succession was originally proposed by Connel and Slatyer (1977). |
| This suggests that, at our site, dense Festuca paniculata dominated grasslands may be an arrested successional stage (or para-climax **?**) |
| Compositional changes in connected trophic levels (herbivores, predators, decomposers) in such successional ecosystems have long been assumed to be the derivative of such plant-driven succession (Connel and Slatyer 1977) |
| Naturalised shmb species facilitate (sensu **?**) the establishment of taller native vegetation in one region, but they may inhibit establishment in another. |
| Some argue that succession depends on context and therefore follows no consistent patterns and no general theory (e.g., McIntosh 1999; Walker and del Moral 2003). Others argue that consistent and identiable mechanisms operate during succession (e.g., Connel and Slatyer (1977)) |
| The successional models of Connel and Slatyer (1977), for example, might predict that predators will alter succession of prey, but these models do not address how that succession of prey will in turn inhibit particular predators. |
| Under such conditions secondary successions are likely to be very slow, if not halted altogether (i.e. inhibition sensu Connel and Slatyer (1977)). |
| On the north-facing ridges at Awana regeneration is 'inhibited' (sensu Connel and Slatyer (1977)) with recurrent fires further deteriorating soils and so favouring flammable species such as manuka, Hakea spp. and Erica spp. |
| Facilitation is a classic mechanism for succession (Connel and Slatyer 1977). |
| Oldeld succession has long been used to understand the mechanisms that determine long-term plant population dynamics (Connel and Slatyer 1977). |
| Thus, the identity and abundance of species that colonize rst may affect the colonization success of laterarriving species. Such priority effects may be positive (facilitative) or negative (inhibitory) (Connel and Slatyer 1977). |
| In the classic paper on conceptual models for ecological succession, Connel and Slatyer (1977) predicted that their facilitation model would apply to heterotrophic succession. |
| Connel and Slatyer (1977) point out that strong facilitation can lead to fully transitory communities that are nevertheless necessary to reach certain community states (see also Law and Morton 1996). |

Excerpts of text citing Grime (1979)

| |
|---|
| It is useful to summarize environmental gradients into a productivitystress gradient (Grime, 1979) to which various structural and functional characters of ecosystems are related and which largely determine the preference for either spontaneous succession or technical restoration. |
| It is generally accepted that diversity is highest at a moderate level of stress or productivity, as the number of species able to grow is limited physiologically or by competition, respectively, toward the ends of the stressproductivity gradient (Grime, 1979). |
| We used Grime (1979)'s CSR (competitor, stress-tolerant, ruderal) classification of plant strategies (Grime, 1979). |
| Native seedling establishment could be prevented by competition with non-native species, particularly annual grasses such as Avena spp., that are dominant in the landscape (Hobbs 2001) and are likely to be represented in the old-field soil seed banks (Grime, 1979). |
| To examine the potential importance of biological soil crust for ecological rehabilitation, we need a fundamental distinction between high and low abiotic stress systems (Grime, 1979). |
| Looking at Grime (1979)'s strategies, the annual stage was dominated by ruderals and competitive species. |
| In an environment characterised by repeated disturbance and infertility, it is possible that these species constitute a stress-tolerant, disturbance-tolerant group (sensu Grime 1979) |
| According to Grime (1979), all five internally transported annuals reaching fruiting stage on faeces deposits in this study are stress-tolerant ruderals (Klotz et al., 2005) that can endure moderate intensities of stress and disturbance. This strategy is typical for winter annuals (Grime, 1979). |
| The major distinguishing feature of many herbivores is selective disturbance (sensu Grime, 1979) of primary production through diet preference; |
| Differences in initial abiotic conditions (provided they are large enough) may result in different but internally consistent trajectories of ecosystems in trait space during succession. This has been suggested by Grime (1979) in terms of competitors, stressors and ruderals (CSR). |
| The two species common in the retrogressive stages of the Waitutu Chronosequence were tolerant of waterlogging, but were atypical of stress-tolerators (sensu Grime 1979). |
| The pasture sites Cr0 and Ma 1 did show higher gain rates than loss rates of plants and species, paralleling with chronosequence predictions (Fig. 3). Such results are consistent with the hypothesized input of colonizing species, with high recruitment and growth rates, at the beginning of succession (e.g., Grime 1979). |
| The functional groups of the weed species were classified using Grime (1979) C-S-R (competitor, stress-tolerant, ruderal) strategy types (Grime, 1979). |
| The dual nature of environmental filters is particularly evident along stressproductivity gradients in which plant recruitment is limited physiologically at one end and by competition at the other, respectively (Grime, 1979). |
| Our results support the hypothesis that species in this forest trade off the ability to be an effective exploitative competitor, that is, to grow quickly and thereby usurp relatively more available resources, with the ability to avoid mortality when resources are less plentiful (Grime, 1979). |
| (1) perennial grasses and sedges of grasslands and edges (e.g. Brachypodium rupestre, Bromus erectus, Carex flacca and Phleum pratense); (2) annual, small-sized grasses and herbs of dry habitats (e.g. Catapodium rigidum and Sherardia arvensis); and (3) ruderal (sensu Grime 1979) grasses and herbs of disturbed, nitrogen-rich habitats (e.g. Dactylis glomerata and Sonchus oleraceus). |

| |
|---|
| According to Grime (1979)'s theory (Grime, 1979), there are essential differences between the traits of ruderal, competitive and stress-tolerant dominants, with important implications for diversity. Grime (1979) suggested that dominant species with competitive and ruderal strategies have stronger negative impacts on diversity than stress-tolerant species. Fast growing species with the capacity for clonal expansion and dynamic foraging have the highest chance to monopolize resources and to reduce the opportunity of other subordinate species. In a survey of the Central European flora, Prach & Pyek (1999) demonstrated that the most successful species appearing as dominants in man-made successional habitats have the traits predicted by Grime (1979). |
| The competitor life strategy (according to Grime, 1979) was the most typical (but with various transitions to S and R strategy). |
| The life strategies (CSR) are according to Grime (1979)'s system (Grime, 1979). |
| According to Grime (1979), stress includes all constraints limiting community productivity, whereas disturbance is a mechanism limiting plant biomass by causing its partial or total destruction (see White & Jentsch 2001 for a review). |
| Species traits have been classified in many ways that relate to disturbance (Noble & Slatyer 1980; Pavlovic 1994; Walker et al. 1999). For example, the r/K continuum (Grime, 1979) and r/K selection (McArthur & Wilson 1967) suggest that r-traits, which are typical of pioneer species, would be advantageous early after disturbance, and K-traits, which are typical of successor species, would be advantageous later in community assembly. |
| Irregular disturbances which affect species abundances can also decrease the effect of competition on the structure of communities (Grime, 1979). |
| As Villger et al. (2008) pointed out, the competitive, stress-tolerant and ruderal (CSR) strategies according to Grime (1979) are characterized by highly correlated traits. |
| Many fen plant species, such as small sedges, orchids or specialised herbs, are stress-tolerators (Grime, 1979). |
| The following parameters or traits were studied: ecological strategy type (according to Grime 1979)... |
| Mean cover values of affected plant traits after pooling into treatment groups. (a) Growth height 3 (¿50 cm); (b) Hemicryptophytes; (c) Ecological strategy type csr sensu Grime (1979); |
| This abiotic stress predominantly affected annuals and therefore species of the ecological strategy types sr and r (sensu Grime 1979). |
| Species were classified into weed and non-weed species groups according to Grime (1979)'s CSR strategy types (Grime, 1979). |
| Prach (1997) found that participation of C-strategists significantly increased during the first 10 years of succession, while R-strategists decreased. The same results were obtained by Grime (1979). |
| Differences in colonization along the gradient are driven by low moisture or nutrients at low productivity and competitive suppression at high productivity (Grime, 1979). |
| Niche differentiation (Giller 1984), species competition (Huston and DeAngelis 1979, Tilman 1984, Wilson and Tilman 1993) and disturbance (Grubb 1977, Denslow 1980, 1987, Tilman and Pacala 1993) have been proposed as driving forces for high species diversity; an unifying model was hypothesized by Grime (1979). |
| As defined by Grime (1979), stress represents the degree to which the abiotic environmental conditions of a site limit the production of plant biomass, while disturbance represents the degree to which the environmental conditions of a site remove or destroy pre-existing living plant biomass. |
| We obtained a simple measurement model of two latent environmental filters (stress and disturbance) that both successfully captured the observed patterns of covariance between the measured environmental variables and that agreed with the original definitions of Grime (1979). |
| Pioneer species generally remained conceptualized as species that were replaced by post-pioneer species, mainly because of poor competitive abilities (Grime, 1979). |

| |
|---|
| Flows of energy leading to characteristic trophic structure and material cycles within the system were considered to be regulated by abioticbiotic feedback mechanisms. These feedback mechanisms initiated and maintained the succession, in addition to biotic autogenic processes (e.g. competition). The resulting effects of feedbacks were viewed as being controlled by the relative importance of stress and disturbance subjected partly to the activities of and competition between organisms (Grime, 1979). |
| The return of pioneer species is quite easy, because they are adapted to long periods of dormancy until the appropriate conditions appear (Grime, 1979). |
| Theories predict that species adapted for surviving and/or competing well in unproductive soils (i.e., stress-tolerators of Grime (1979); belowground competitors of Tilman [1988]) are at a competitive disadvantage in productive soils. |
| The predictions of general theories are complicated further by the fact that some species adapted to nutrient-poor soils are more resistant to disturbance than other species (Grime, 1979). |
| Forest disturbance is typically characterized by biomass removal (Grime, 1979). |
| It is still unclear how these species interactions change along an environmental gradient consisting of multiple (often interacting) abiotic stressors and biotic stressors (herbivory or consumer pressure, i.e. disturbance sensu Grime 1979). |
| Dense woody canopy induces environmental stress (sensu Grime 1979) for ant communities in general (and for grassland ants in particular) by negatively affecting surface temperature, and thus it can negatively affect ant community structure (Gmez et al. 2003). |
| We quantified disturbance using Grime (1979)'s definition, as the mechanisms which limit the plant biomass by causing its partial or complete destruction. |
| Life-history traits that lead to stress tolerance typically reduce competitive ability and vice versa (Grime, 1979). |
| An explanation for this phenomenon is the intermediate disturbance hypothesis (Grime, 1979). "Grime (1979) reported that spontaneous vegetation of disturbed grassland starts with the invasion of pioneer or opportunistic species. " |
| Hence, plants on the upper shore are considered to be better competitors (sensu Grime 1979) than species on the lower shore and where their niches overlap the higher shore species eventually displace lower shore ones (Bertness 1991; but see Bockelmann & Neuhaus 1999). |
| The strong positive correlation between mean tree growth and crowding importance for surviving trees conformed to hypotheses suggesting that crowding (or competition) importance increases when productivity is higher (Grime, 1979), at least for those trees that survive. |
| Thirty percent out of 27 plant species exhibited a ruderal Grime (1979) strategy (Grime, 1979). |
| High representation of ruderal plant species in traps was expected (Grime, 1979). |
| The cutting and removal of biomass has enabled the persistence of small plant species (now often rare or red-listed) with a low ability to compete for light but that are resistant to mechanical stress (Grime, 1979). |

Excerpts of text citing Connell (1978)

| |
|---|
| The intermediate disturbance hypothesis forecasts highest species diversity at sites subject to intermediate rates and intensities of disturbance because there will be a mixture of pioneer and climax species (Connell, 1978). |
| One possible explanation may be the intermediate disturbance hypothesis Connell (1978), stating that environments with intermediate rate of disturbance display highest diversity. |
| Perhaps the most common and intuitive prediction is that diversity should peak at an intermediate level of disturbance, the so-called intermediate disturbance hypothesis (IDH, Connell, 1978). |
| The widespread perception that older stages do not harbor a high richness as suggested by intermediate disturbance hypothesis of Connell (1978). |
| The verbal model of the IDH assumes a uniform trade-off from the best colonists to the best competitors, such that very few species immediately invade newly opened patches, while very few species ultimately limit all others over long time scales (Connell, 1978). |
| Another outcome, unexplored for bats to date, is that under an intermediate level of disturbance, species richness will be higher than in an undisturbed habitat, as set forth in the intermediate disturbance hypothesis (Connell, 1978). |
| There is little consensus in how species richness and diversity change over successions (Huston and DeAngelis 1994; Rosenzweig 1995), with various models predicting richness and diversity peaking at either early (e.g. the initial floristic composition model of Egler, 1954) or intermediate (e.g. the intermediate disturbance hypothesis of Connell, 1978). |
| We postulated at the start of this study that, based on the intermediate-disturbance hypothesis (sensu Connell, 1978), we would observe the highest levels of observed species richness at sites subject to intermediate numbers of fires (23 fires since 1972) and an intermediate period of time since fire (1216 years). |
| This hump-shaped distribution of diversity as a function of time since disturbance matches the predictions of the intermediate disturbance hypothesis (Connell, 1978), in which community diversity increases over time as new microhabitats are created. |
| Moderate disturbances (Connell, 1978) may also decrease the strong competitive interactions that may be present at high productivity (Huston 1979; Wilson & Keddy 1986; Campbell & Grime 1992; but see Wilson & Tilman 1991). |
| Connell (1978) proposed in his intermediate-disturbance hypothesis that intermediate-disturbance frequencies produce the highest diversity. |
| One of the more popular contemporary hypotheses is the intermediate disturbance hypothesis which suggests that biodiversity is maximized where the frequency and/or intensity of disturbance occurs at an intermediate level (Connell, 1978). |

The intermediate disturbance hypothesis is limited to the patch scale. Indeed, Connell (1978) excluded the landscape scale, stating that the hypothesis would not hold for geographical gradients.

A hump-backed curve of species richness has been described for disturbance gradients as the "intermediate disturbance hypothesis" (Connell, 1978).

A pattern with highest abundance and diversity at intermediate successional stages was found in a moth recovery study in the Neotropics (Hilt & Fiedler 2005), obeying the intermediate disturbance hypothesis (IDH; Connell 1978). According to this hypothesis, the diversity of early successional stages is low, as only few pioneer species have been able to disperse there. The diversity peaks at the intermediate stages, as more species invade the area. Finally, in the late stages of succession, diversity decreases again, due to competitive exclusion.

Prominent disturbance theories include the intermediate disturbance hypothesis (IDH; Connell 1978), which predicts the occurrence of highest species richness at intermediate levels of disturbance.

As predicted by the IDH, peak richness was expected to occur at the mid-point of forest succession where pioneer and climax species co-exist (Fig. 1a; Connell, 1978).

Post-fire changes in species richness did not follow the bell-shaped curve as proposed by the IDH (Connell, 1978).

The first derives from the intermediate disturbance hypothesis (IDH), which predicts that species diversities will be highest in areas that are subject to moderate levels of disturbance (Connell, 1978).

These habitats are characterised by intermediate levels of environmental and disturbance gradients, confirming the intermediate disturbance theory (Connell, 1978).

Although Connell (1978) emphasised the chance aspect, the core of this mechanism is the similarity in interference ability, i.e. the neutrality.

The necessary conditions are: (1) there is disturbance of patches at a scale smaller than the one we are considering for the paradox; (2) this disturbance occurs with a frequency such that there will a mixture of patches of different time since disturbance (this is the Intermediate Disturbance Hypothesis of Connell (1978); successional niche has been used; this may be another aspect of the rather general term regeneration niche).

I do not know of any theory directed at this. P.W. Richards suggested that in tropical rain forests, normally diverse, mono-dominance was to be found in unfavourable habitats (Salisbury 1931), but Connell (1978) suggested that mono-dominance is the climax situation, and the iconic species-rich tropical rain forest communities are recovering from disturbance.

Examples of theoretical concepts include the intermediate disturbance hypothesis (Connell, 1978), general successional theory (Luken 1990; Johnson and Miyanishi 2008), habitat accommodation theory (Fox 1982), the initial conditions hypothesis (Egler, 1954), ecological thresholds (Groffman et al. 2006), resilience thinking (Walker and Salt 2004), and biological legacies (Franklin et al. 2000).

| |
|---|
| Diversity may peak at an intermediate stage, when both short-lived colonists and longer-lived, but disturbance-sensitive, species are present (i.e., the intermediate disturbance hypothesis; Connell,1978). |
| The effect of disturbance depends on the frequency and intensity of the disturbance events (Connell, 1978). |
| Species richness was significantly higher in blowouts as compared to controls pointing to the effect of gaps making room for more species early in the establishment phase (Connell, 1978). |
| Ecological theory and empirical studies have suggested that biological diversity is greatest in environments subject to intermediate levels of disturbance (Connell, 1978). |
| Since the intermediate disturbance hypothesis was popularized by Connell (1978). |
| The first states that biodiversity increases at localities subjected to intermediate rates of disturbance (Connell, 1978). |
| Disturbance has long been proposed as a mechanism enabling the coexistence of species, with a unimodal relationship between diversity and the level of disturbance predicted by the intermediate disturbance hypothesis (IDH; Connell, 1978). |
| Because different sets of species are affiliated within recently disturbed and long undisturbed forests, nonequilibrium maintenance of species diversity is hypothesized to be favored by intermediate frequencies of disturbance (the intermediate disturbance hypothesis Connell, 1978). |
| This pattern fits the prediction of the intermediate disturbance hypothesis proposed by Connell (1978), according to which an intermediate incidence of disturbance is often translated into higher diversity of organisms. |
| The intermediate disturbance hypothesis describes how frequency, magnitude and time since disturbance may affect plant diversity, and predicts that the highest number of coexisting species should be found at intermediate magnitudes of these processes (Connell, 1978) |
| Natural and cleared sites were more similar to each other than to alien sites, which confirmed that clearing initiated recovery and did not exacerbate disturbance, in accordance with the intermediate disturbance hypothesis (Connell, 1978) that predicts maximal species diversity between the extremes of high stress (alien invasion) and complete stability (natural sites). |
| The obtained data were best fitted by a humpshaped curve, as suggested by the intermediate disturbance hypothesis (IDH; Connell, 1978). |
| Connell (1978) hypothesized that the highest diversity in tropical forests should occur under a disturbance regime with intermediate frequency and intensity. Too little disturbance results in the loss of light-demanding species in late successional forests through competitive exclusion, and too much disturbance results in the loss of species that compete well in full shade. |

This last result may suggest an intermediate gradient hypothesis (similar to the intermediate disturbance hypothesis of Connell 1978) where species from both the dry and very wet forest can exist in the wet forest and add to its diversity.

If disturbance is defined as removal of biomass (sensu Grime) then the intermediate disturbance model (Connell, 1978) will also predict higher richness at intermediate biomass levels, i.e. intermediate succession phases.

Different hypotheses included equilibrium mechanisms through niche partitioning (Tilman 1994); non-equilibrium coexistence dynamics (Huston 1994) related to disturbance (Connell, 1978), predation and biotic interactions (Janzen 1970, Wills et al. 1997); fluctuations of environmental conditions (Chesson 2000); and balance between immigration/speciation and extinction (McArthur and Wilson 1967, Hubbell 2001).

Indeed, its integration with other frameworks such as the intermediate-disturbance hypothesis (Connell, 1978) and the immigration/extinction balance theory (McArthur and Wilson 1967), does produce an improved theoretical background to understand biodiversity

However, despite the importance of evenness in explanations for the maintenance of species richness in tropical forests (Connell, 1978), changes in species evenness with succession are poorly explained.

The most prominent hypothesis of how disturbances shape diversity is the intermediate disturbance hypothesis (IDH; Connell 1978). The IDH predicts that species richness is maximized at intermediate levels of disturbance, because competitively dominant species exclude other species at low levels of disturbance, whereas at high disturbance levels only the most resistant species subsist.

This hypothesis is generally based on the idea that a trade-off in species life history traits prohibits species with poor colonization traits (such as seed number) at high disturbance, excludes less competitive species in older undisturbed communities, but allows a mix of both groups of species at intermediate disturbance (Connell, 1978).

From this hypothesis, we derived the predictions that succession develops as a net increase in the number of taxa (e.g. Connell 1978) and that community structure progressively changes during succession from one structure to a different one.

The IDH posits that biodiversity is highest in communities when disturbance is neither too rare nor too frequent, i.e., in communities with intermediate levels of disturbance (Connell, 1978).

Peaked diversity patterns are consistent with the intermediate disturbance hypothesis, when species coexistence is maximized between disturbance-tolerant but competitively less dominant species and disturbance-intolerant but competitively dominant species (Connell, 1978).

Our results also fit the intermediate disturbance hypothesis (Connell, 1978), which predicts a greater species richness in communities subjected to moderate levels of disturbance.

| |
|---|
| These results agree, at least qualitatively, with the Connell (1978) hypothesis, which states that intense and long disturbance events favor the development of a few phytoplankton groups, leading to a population with low diversity, while disturbances of intermediate frequency would lead to higher diversity. |
| The frequency of natural disturbance (sand movement) in these habitats is intermediate (Martnez et al. 2001) which thus supports the intermediate disturbance theory of Connell (1978). |

| Excerpts of text citing Odum (1969) |
|---|
| Odum (1969) proposed trends associated with ecosystem development and the Hubbard Brook studies of ecosystem response to disturbance. |
| Odum (1969)s strategy of ecosystem development helped set the stage for hypothesized functional dynamics with time since disturbance. |
| Community stability increases over the course of succession (e.g., Odum 1969). |
| Over time, plant growth and biomass accumulation increase, and the input of new litter causes the detrital pools to build up, resulting in an extended period of carbon accumulation and a positive NEP (net ecosystem production). Finally, the buildup of Clive, Cforest floor, and CCWD (coarse woody debris) cause mortality and decomposition losses to accelerate, which moves the pools toward steady state and causes NEP to approach zero in the oldest stands (Odum, 1969). |
| Nonetheless, the NEP and carbon storage efficiency (net ecosystem production/total net primary production) observations support the hypothesis that old stands approach steady state (Fig. 1b and e; Odum 1969). |
| Decreasing vegetation quality (Van der Wal et al. 2000b) and increasing structural complexity (increasing amount of dead organic material and standing biomass, and increasing vegetation height) are typical for many successional sequences (Odum, 1969). |
| Future data will refine our current understanding of the impacts of altered hydrology on vegetation succession in the Everglades and increases our ability to apply succession theory to resolve restoration issues (Odum, 1969). |
| Odum (1969) hypothesized that forest stands experience an initial postdisturbance reduction in carbon storage due to higher respiration rates, with NEP (net ecosystem production) increasing to a maximum as canopy assimilation peaks and declining thereafter. |
| Annual carbon storage in our forest chronosequence followed the successional trajectory predicted by Odum (1969), increasing to a maximum and then gradually declining as the forest matured. |
| Succession provides a temporal framework in which to understand ecological processes such as species assembly (Walker & del Moral 2008), vegetation dynamics (Marrs & Bradshaw 1993) and ecosystem development (Odum, 1969). |
| Our results are consistent with this, since after the hydroseeded stage (20042005) important changes in species composition occurred; there was an influx of widespread native species (2006) followed by a decline in turnover with time, thus increasing the community stability (e.g. Odum 1969). |

| |
|---|
| "The measured flux results are discussed first in the context of the three hypotheses. We then place the results in the broader context of classic and contemporary ecological ideas regarding:<br>(E1) the role of assimilation and respiration in controlling the net C flux of terrestrial ecosystems (Valentini et al., 2000; Reichstein et al., 2007); (E2) C exchange along ecological succession, with a focus on the Strategy of Ecosystem Succession of E. P. Odum (1969); and correspondingly (E3) the role of ecosystem resistance and resilience to disturbances such as droughts and ice storms in maintaining the ecosystem service of C sequestration." |
| These results largely followed expectations based on the Strategy of Ecosystem Development of Odum (1969), which hypothesizes that GPP (gross primary production) rapidly increases with ecological succession, then decreases as forests age, while RE (respiration) increases monotonically due to the increase in autotrophic biomass. |
| Modernizing the Strategy of Ecosystem Development (Odum, 1969) to take into account the relationship between GEP and RE may improve understanding of biosphereatmosphere C exchange over successionary time scales and, combining our results with others, may help dispel the prevailing idea that mature forest ecosystems with large C pools must be small C sinks (see Carey et al., 2001; Rser et al., 2002; Knohl et al., 2003; Zhou et al., 2006; Urbanski et al., 2007; Baldocchi, 2008). |
| Odum (1969) additionally hypothesized that early successional ecosystems maximize productivity, while late successional ecosystems maximize protection against (i.e. resistance to) environmental variation. |
| Over relatively short time-scales (decades to centuries), ecosystem properties such as net primary productivity increase and reach maximum values (Odum, 1969). |
| Succession generally refers to the biological changes that occur in an ecosystem after a clearing or exposure of an area, often resulting in predictable sequences of species composition shifts Odum (1969). |
| Diversity is hypothesized to increase as more species accumulate through migration but then decrease through successional time as less successful competitors are eliminated (Odum, 1969). |
| Based on Odum (1969)'s theory of ecosystem succession (Odum, 1969), this index declines both during succession and following recovery from disturbance, because equilibrium conditions are approached and the soil microbial community becomes more carbon efficient (Anderson and Domsch 1985, Wardle and Ghani 1995). |
| Odum (1969) hypothesized that net ecosystem production (NEP) increases during early succession before beginning a gradual decline to near zero in older stands, due in large part to declining net primary production (NPP). |
| Overall, Odum (1969) suggested that variable nutrient supplies in later succession would favour larger animals, which further confirms our observations. |
| First, if species turnover is low and colonization is continuous, diversity may increase over time, peaking late in succession (Odum, 1969). |

| |
|---|
| Trends in diversity on mounds did not support predictions of the intermediate disturbance hypothesis (Hobbs and Hobbs, 1987; Huntly and Inouye, 1988), but rather, a model of gradual species accumulation (Odum, 1969). |
| The most commonly acknowledged hypothesis, originally proposed by Kira and Shidei (1967) and popularized by Odum (1969), is that total photosynthetic assimilation (that is, gross primary productivity) of trees is increasingly offset by higher autotrophic and heterotrophic respiration as stands age. |
| Diversity can increase asymptotically through succession (Odum, 1969). |
| The idea of increasing stability during ecological succession was first introduced by Odum (1969). |
| Despite Lugos conclusions that mangroves are steady-state ecosystems he also suggested that mangroves may satisfy many of the criteria for succession developed by Odum (1969), which included characteristics of community energetics and structure, life history, nutrient cycling, and overall homeostasis. |
| We tested the following hypotheses: (1) mangroves are youngest on the seaward fringe and oldest on the landward edge similar to patterns described by Fromard and others (1998); (2) mangroves follow succession patterns described by Odum (1969); (3) C stocks in soils increase with age consistent with the model of Chen and Twilley (1998); (4) soil respiration varies with forest age, declining as forest productivity declines (Alongi and others 2008); and (5) N limitation to growth increases with forest age consistent with observations in terrestrial forest chronosequences (Vitousek and others 1993). |
| Tests of differences among characteristics of forests of different ages were used to assess successional patterns. Successional criteria were derived from Odum (1969), listed below) and estimated from data gathered at the site. |
| Using the forests of differing ages we were able to assess changes in forest and soil characteristics as they aged to test whether processes that occur during the maturation of mangrove forests are concordant with predicted successional patterns (compare Odum 1969). |
| Mangroves do not transition from linear grazing food webs to detritus based food webs as proposed by Odum (1969), but instead are dominated by detritivores throughout their development. |
| Here we propose an additional level of reflection which extends the idea that the succession process results from modifications of the physical environment driven by the communities (succession sensu Odum 1969). |
| Odum (1969) adopted a more integrated view by considering ecological succession as a developmental process at the scale of ecosystems. Odum (1969)'s concept originated from the early idea of succession proposed by Cowles (1899), which realized that biological and physical components of ecosystems co-adjust in time and space. The ecosystem was considered from Odum (1969)'s point of view as a system where communities of organisms in a specific area interact bi-directionally with the physical and chemical environments. |

Immature marshes with low biomass can be expected to have relatively low GPP (gross primary productivity) and R (respiration) compared to mature marshes with high biomass. As created marshes age, GPP, R, and NEE (net ecosystem exgenge) should increase to levels that meet or exceed those characteristic of mature ecosystems (Odum, 1969).

”Thus, the appropriate comparisons for assessing change in gas exchange rates over time were between created and reference marsh pairs, and these data suggest that either gas exchange attributes develop very rapidly (i.e., ¿ 3 years) or the model proposed by Odum (1969) does not apply to created Marshes.”

To assess early and intermediate successional patterns in field CO2 exchange as described by Odum (1969), future research should focus on following young (15 years old) created marshes with frequent sampling.

This is in agreement with Odum (1969), who argued that increased control of the physical environment is a mechanism that allows the ecosystem to achieve maximum protection from perturbations.

In a balanced ecosystem, as Odum (1969) suggested, the ecosystem-level ratio between production and respiration (P/R) approaches 1.

Odum (1969) further suggested various ecosystem attributes between developmental and mature stages: mature stages are characterized by complex, well-organized, low-growth, and high-information structures.

Also typical features of the dominating organisms may be used to characterize ecosystem states: the longer complexifying dynamics have been affecting the system at a site, the higher should be the probability to find organisms with long life spans, high body size and body mass, and with an increasing degree of specialization (Odum, 1969).

To describe the development of eco-physiological features, already Odum (1969) has proposed to focus on certain ecosystem attributes which are regulated by the interactions between the habitat’s organisms.

Communities in early succession are usually dominated by fast-growing species (colonisers), while in later succession by slowgrowing species (competitor) (Odum, 1969).

Succession has been classically described from terrestrial systems as an orderly and directional process resulting in a stable climax community (Odum, 1969).

Organic matter (OM) generally accumulates in ecosystems as succession proceeds (Odum, 1969).

The metabolic quotients were high in young soils and low in mature ones (Odum, 1969).

During succession, resistance increases and resilience decreases (Odum, 1969).

Previous studies on the impacts of forestry drainage and restoration show that peatlands tend to conform to the theory of Odum (1969): The nutrient-poor, late-successional bogs appear to be resistant systems with slow and often partial secondary succession following drainage (Laine et al. 1995), and slow regeneration after restoration (Jauhiainen et al. 2002; Laine et al. 2011).

| |
|---|
| Forests tend to conserve more energy with succession, that is a reduced ratio of maintenance to structure (Odum, 1969). |
| The specific RH (heterotrophic respisration) and specific RA (autotrophic respiration) were used to revisit Odum (1969)'s theory of ecosystem succession (Odum, 1969). We hypothesized that both specific RH and specific RA would decline with forest succession based on Odum (1969)'s theory. |
| Odum (1969)'s theory of ecosystem development predicts that the ratios of respiration to biomass decrease with the forest approaching a climax stage of development since the biomass becomes more energy efficient (Odum, 1969). |

Excerpts of text citing Harper (1977)

| |
|---|
| During the first stage (pioneer colonization, ¡ 20 years; Fig. 8: Stage 1), a limited number of moss and lichen species establish on the newly-emplaced lava, forming patches in small surface irregularities (¡ 102 m in scale; N.A. Cutler et al., unpubl. data) that function as safe sites (Harper, 1977). |
| Seed and seedling mortality from random and non-random factors is immense (Harper, 1977). |
| Second species colonization from the surrounding species pool, which was more prevalent in the south and flat sites, might have been a biotic feedback effect with seedling establishment of autochthonous species reduced through competition from the greater abundance of hydroseeded species (Matesanz et al., 2006 ; Gonzlez-Alday and Martnez-Ruiz, 2007). Colonizing seedlings and juveniles are very sensitive to such biotic competition (Harper, 1977). |
| It is known that there is spatial variability in the distribution of seeds in the seed banks of sand dunes (Zhang and Maun 1994; Baptista and Shumway 1998; Cheplick 2006), and that they can often accumulate in clumps or safe sites (Harper, 1977). |
| Sprouting may result in several individuals (clones) originating from one parent plant (Silvertown 1987). Although sexual reproduction will provide greater genetic variability (Harper, 1977), vegetative propagation through ramets may result in the survival of resistant individuals (Miller & Kauffman 1998). |
| Four of the 13 species that reproduce both by seed and sprout are in different successional stages (Appendix). These species invest heavily in sexual and asexual reproduction, which may be characteristics of pioneer species (Harper, 1977). |
| Replacing vertebrate grazing with annual cutting can also lead to large changes in vegetation structure (Harper, 1977), which might modify the effect of invertebrate herbivores. |
| The expression seed bank can be defined as all the viable (dormant as well as ready to germinate) seeds contained in the soil in a given area (Harper, 1977). |
| Sometimes it may also be useful to differentiate between the currently germinable proportion of the seed bank (the active seed bank) and the part that will become germinable in the future (the dormant seed bank, Harper 1977). |
| A review of the results obtained at one Calluna heath and seven meadow sites confirms the special importance of the top layer of soil (Harper, 1977). |
| Soil surface microhabitats are a harsh environment for seeds of many vascular plant species (Harper, 1977). |
| Population ecologists, influenced by Harper (1977), study demographic processes, population fluctuations and spatial distributions. |
| Although the seed input from herbivores is low in relation to the total seed rain, it may nevertheless have implications for community structure, as seeds are dispersed and concentrated in localised patches of dung (Harper, 1977), which may facilitate local establishment and stand species enrichment. |
| The species composition is itself affected by successional change during the restoration process, as the processes affecting the propagule bank are dynamic (Harper, 1977). |
| In fire-prone ecosystems, outcrops may act as safe sites (sensu Harper 1977). |
| The succession of plant communities is basically governed by adult decline and seedling establishment (Harper, 1977). |

| |
|---|
| The combination of germination requirements (Barik et al. 1996), spatial availability of safe sites (Harper, 1977), and timing of dispersal all influence germination and establishment success. |
| However, during decomposition, both the abundance and the activity of phytotoxic compounds continuously change over time by their sorption and polymerisation on soil organic matter and clay minerals (Makino et al. 1996), and because of the chemical transformation by microorganisms (Blum et al. 1999). This was explicitly assessed by Harper (1977) in a review of allelopathy, who pointed out that plant-produced phytotoxic compounds, being rapidly degraded by the soil microbial activity into non-toxic molecules, should have a limited expected impact on plant population dynamics. |
| Our results support the view that investment in reproduction trades off with investment in growth (Harper, 1977). |
| The establishment of seedlings is known to be a crucial life stage in plant population ecology (Harper, 1977). |
| We hypothesised that fire enhances S. bracteolatus recruitment because of an increase in abundance of suitable microsites (safe sites sensu Harper 1977) after fire. |
| For long-lived woody plants, seedling emergence, establishment and growth are probably the most critical phases of their life cycles (Harper, 1977), and are the most susceptible to competition with grassland species and damage by herbivores. |
| Fine-scale heterogeneity may result from physical variability, microtopography, or population processes such as seed dispersal patterns that leave seeds more or less concentrated around the parent plant (Harper, 1977). |
| The abiotic factors and the availability of safe sites (sensu Harper 1977) may be more important for the species diversity and composition than the biotic factors (Billings and Mooney 1968, Grime 1977). |
| Despite the relatively high environmental severity at alpine Finse, biotic interactions, such as competition from the resident vegetation appeared to be more important for the establishment of species and for the community diversity than the availability of safe sites (sensu Harper 1977) and experimental warming. |
| Plant density in mixtures was assessed by counting all plants grown from seeds (i.e., genets in the terminology of Harper 1977). |
| This correlation was strong at the site level but virtually non-existent at the quadrat (local) level, the scale at which I would expect resource competition to occur (Harper, 1977). |
| The actual mechanisms by which plants interact with each other and the environment occur at small spatial scales (Harper, 1977). Stand-scale studies can only provide inferences about mechanisms based on correlations with disturbance type, environment, or time since disturbance. |
| Rather, our results suggest that preferential safe sites (sensu Harper 1977) are shared by many colonizing species, but that species interactions reduce the establishment success of these early colonists. |
| Competition studies conducted with the replacement series methodology have shown that the RY total of some species mixtures (RYT; the sum of relative yields of component species) varies with species abundances (e.g., Harper 1977), implying that the average RY of species in mixtures of certain plants may change as required for a change in the complementarity effect. |
| Moderated microenvironmental conditions may potentially serve as microscale refugia or safe sites (sensu Harper 1977) for tree seedlings in the context of global warming during the 21st century (IPCC 2007) within a climatically sensitive ecotonal region of the aspen parkland. |
| Harper (1977) reported an average value of reproductive efforts in annuals ranging from 15% to 30%. |
| At the individual plant level, absence of microsites (Harper, 1977) for germination has been proposed as a determining factor limiting establishment of plants. |
| Density-dependent mortality occurred, which is generally considered a symptom of competition (Harper, 1977). |
| Much less known is the population structure of clonal species, characterized by a multi-shoot architecture of genets, longevity of individuals, as well as genetative and vegetative propagation (Harper, 1977). |
| Soil seed banks are main constituents of ecosystems (Harper, 1977). |

| |
|---|
| The seed bank, defined as all viable seeds contained in the soil of a given area (Harper, 1977), holds a record of previous vegetation, as well as potential permutations for future plant communities. |
| The term safe site refers to a site with edaphic conditions or opportunities suitable for successful seedling recruitment (Harper, 1977). |
| In the rain shadow of the Andes in Argentina, western sites are much more humid than eastern ones. Because cypress reproduces almost exclusively by seed, regeneration is contingent upon the chance of seeds finding safe sites (sensu Harper 1977) to successfully germinate and become established. |
| Grassland plants have two ways to regenerate their populations: vegetative shoots which originate from bud banks and seedlings from seed banks. The input and output of bud banks (Harper, 1977) via vegetative reproduction by rhizomes, or other perennial organs, are of great importance to the recruitment and regeneration of the population in subsequent seasons and years (B enson et al. 2004, Klimeov and Klime 2007). |
| Herbivory and animal trampling can also affect species composition at the seedling stages by creating gaps inside forest patches, thereby providing safe sites for some species (Harper, 1977). |
| The bud bank consists of dormant meristems that are comprised of bulbs, bulbils, and buds on rhizomes, corms, and tubers (Harper, 1977). |
| A seed bank is defined as the soil seed reservoir, and seeds from it are capable of germination when conditions are favourable (Harper, 1977). |
| A seed bank is defined as the soil seed reservoir, and seeds from it are capable of germination when conditions are favourable (Harper, 1977). |
| Further, the belowground bud bank (sensu Harper 1977) contributes to re-sprouting in response to seasonal climates and disturbance (Rogers and Hartnett 2001; Klimeov and Klime 2007; Carter et al. 2012). |
| The dispersal curve, which indicates the change in expected seed density with increasing distance from the parent tree, is one of the traditional models of seed dispersal (Harper, 1977). |
| A higher final germination percentage may be beneficial by increasing the number of potential seedlings that can establish and survive to subsequent life stages, hence increasing plant fitness (Harper, 1977). |
| Seeds disperse after ripening and then form a seed rain, which is defined as the sum of all the seeds from mother plants in a given time and area (Harper, 1977). |
| In grassland plant communities, where most species are clonal (van Groenendael & de Kroon 1990) and where sexual recruitment is low (Harper, 1977), space pre-emption mostly depends on clonal traits (Herben et al. 1994; Gough et al. 2001). |
| Each clonal fragment was composed of one mature ramet (i.e. an erect shoot, its leaves and roots; Harper 1977) with one connected spacer for species from G and I clonal groups, and of three joined ramets forming a tuft for species from P group (i.e. spacers were almost nonexistent). |

Excerpts of text citing Pickett & White (1985)

---

Although small patch or gap disturbances were recognized as sources of spatial heterogeneity (Pickett & White, 1985), the occurrence of large catastrophic disturbances raised the specter of extensive areas being homogenized and even destroyed.

---

Pickett & White, (1985)'s book, Natural Disturbance and Patch Dynamics, ushered in a period of concerted attention to natural disturbances in a wide range of systems and emphasized spatial heterogeneity in ecosystems.

---

Describing the five components of natural disturbances quantitatively requires interdisciplinary collaboration. A distinction is made between external (allogenic) forces that converge on ecosystems and the internal (endogenic) processes of the system (Pickett & White, 1985).

---

Formal studies of plant succession have been conducted since 1895 (Warming 1895) and much has been learned about how ecosystems respond to a dynamic physical environment (Pickett & White, 1985).

---

As long recognised since at least Cowles (1899), many real communities are in a transient, not stable, state, because disturbance keeps communities from reaching a stable state (reviewed in Pickett & White, (1985)).

---

It is important to also consider the effect of disturbance frequency and magnitude on transient community states (Pickett & White, 1985), which we did not explicitly examine in this paper.

---

Disturbances not only remove vegetation from suitable sites, creating unoccupied habitats that plants must colonize from seed banks or outside sources, they also cause changes in the abiotic environment that may favor the establishment of certain species (Pickett & White, 1985).

---

Disturbance was well established as a focal topic of ecological research by the late 1980s. Concepts such as patch dynamics (Pickett & White, 1985), the shifting mosaic steady state (Bormann and Likens 1979; Paine and Levin 1981), landscape equilibrium (Romme 1982), landscape heterogeneity (Turner 1989), and nutrient loss and retention (Vitousek and others 1979) were receiving considerable attention.

---

Although the relative success of juvenile trees is only one phase of forest development, individuals in the sapling bank typically have the greatest opportunities of attaining canopy dominance following the death of a single or several canopy trees (Pickett & White, 1985).

---

Different sorts of disturbances have different effects (e.g. Pickett & White, (1985)).

---

The deficiencies in that form of succession model do not apply to models that use the definition of succession common in the non-savanna literature, i.e., a definition that does not assume reversibility or a controlling role for grazing (Pickett & White, 1985).

---

It is critical to understand the consequences of disturbances because they are ubiquitous and because they are major determinants of ecosystem patterns and processes (Pickett & White, 1985).

---

The structure and specific composition of plant communities can be strongly determined by spatial and temporal patterns of plant recruitment, particularly after dramatic disturbances such as fire (Pickett & White, 1985).

---

Thus, plant communities will likely reflect a complex synergism of disturbance characteristics that affect plant performance directly by releasing limiting resources (Pickett & White, 1985) and indirectly by modifying herbivore foraging patterns (Stuth 1991).

---

The landscape-scale coexistence of different communities representing the successional states depends on disturbances that prevent the persistence of a climax state (Pickett & White, 1985) and on the possibility of re-invasion into early successional patch states from neighbouring patches.

---

Ecologically, a disturbance rapidly changes resource availability, which provides opportunities for reorganization in biological communities (Pickett & White, 1985).

---

The highest plant species diversity and species density was found around plot 3 at an intermediate age of approximately 150 years. This finding is consistent with the general theory of plant succession, which suggests that intermediate successional stages (Grime 2001) or intermediate disturbance regimes often have the highest plant diversity (Pickett & White, 1985).

---

In addition, natural landscapes are themselves dynamic, with local communities continually being influenced by disturbance (Pickett & White, 1985). The pervasive roles of disturbance and succession for community assembly suggest that temporal dynamics may play a key role in observed LSRRSR relationships.

---

While the importance of the frequency and intensity of small-scale disturbances in rocky-shore communities (Sousa 1979a, b, 1984), temperate and tropical forests (Schaetzl et al. 1988, McCarthy 2001), and elsewhere (Pickett & White, 1985) is well appreciated, this study is the first to our knowledge to explicitly test how the species richness or composition of the surrounding community acts in concert with variation in consumers to impact recovery.

---

The turnover time of a tree population was calculated as the inversion of its mortality rate (Pickett & White, 1985).

---

With the same disturbance rate in different forest types, as suggested in Pickett & White, (1985), the forest structure will depend on the growth rate, limited by directly available resources, generating sparse forests in the boreal regions and dense forests in tropical rain forests.

---

Disturbance has been acknowledged as an important factor affecting community composition and biodiversity in natural ecosystems (Pickett & White, 1985) with highest diversity predicted at intermediate disturbance (Connell, 1978; Hobbs and Huenneke, 1992 ; Huston, 1979).

---

The composition of post-disturbance vegetation will also depend upon how well the disturbed species recover and compete with new arrivals, which is typically a function of their life history traits (Pickett & White, 1985).

---

Agricultural land uses can be considered ecological disturbance regimes (sensu Pickett & White, (1985)) with the potential to affect the speed and extent of forest regeneration in abandoned fields (Guariguata & Ostertag 2001; Myster 2004; Chazdon et al. 2009).

---

| |
|---|
| The EDI index, built with information provided by farmers and landowners, represents an advance for quantifying disturbance regimes (sensu Pickett & White, (1985)) associated with agricultural practices. |
| Our Ecological Disturbance Index (EDI) was derived from the analysis of agricultural land uses as ecological disturbance regimes (sensu Pickett & White, (1985)). |
| As the disturbance creates patches that are more exposed to light and high temperatures, they imposed negative effects in the dry year. This is in accord with the results from Ryser (1990) who found that under very dry conditions, mortality in gaps may be higher than in closed vegetation. The finding also supports the conclusion made by Pickett & White, (1985) that disturbance may interact with stress, resulting in different physical effects of a disturbance at a given site. |
| In real landscapes, however, disturbance and succession drive spatial and temporal variability in patch turnover and suitability (Pickett & White, 1985), which are not adequately incorporated in most models. |
| Disturbance plays an important role in determining the structure and functioning of ecosystems (Pickett & White, 1985), and is also important in maintaining species diversity (Denslow et al. 1998). |
| As a nutrient conservative species with low leaf nutrient and high nutrient resorption efficiency (Yan et al. 2006; Huang et al. 2007), S. superba could persist in soils depleted by soil erosion which often accompanies intense disturbance (Pickett & White, 1985). |
| Variation in the frequency, extent, and intensity of disturbance events has a profound effect on the nature of landscapes (see Pickett & White, (1985)). |
| The theory of disturbance (Pickett et al. 1989; Walker and Willig 1999; Willig and McGinley 1999; Willig and Walker 1999) has matured considerably since the seminal publication of Pickett & White, (1985). |
| Variable disturbance regimes can help maintain diversity by facilitating the coexistence of species from different successional stages (Pickett & White, 1985). |
| A definition of ecological disturbance was provided by Pickett & White, (1985) as any discrete event in time that disrupts ecosystem, community or population structure and changes resources, substrate availability, or the physical environment (i.e. alters niche opportunities for the species capable of living in a given setting). |
| Both natural and anthropogenic disturbances can be characterized through their severity or through the amount of damage they cause. Damage can be measured as a loss of biomass, structure or function (Pickett & White, 1985). |
| The quality and quantity of abiotic inputs, namely, materials, water, and solar energy, influence productivity and the response of the ecosystem to disturbance (Pickett & White, 1985). |
| Disturbances produce spatial and temporal heterogeneity in biological communities by facilitating individual and species turnover (Pickett & White, 1985). |

Animals that modify their habitats act as natural disturbance regimes in ecosystems. Though varying in type, intensity, frequency and extent, disturbance regimes are typical of every ecosystem (Pickett & White, 1985).

---

Abiotic and biotic disturbances play a major role in shaping and sustaining forest composition (Pickett & White, 1985)

---

In many ecological systems, disturbance, defined by Pickett & White, (1985) as any relatively discrete event in time that disrupts ecosystem, community or population structure and changes resources, substrate availability or the physical environment, varies spatially among habitat patches (Caswell 1978; Tilman 1982; Townsend 1989).

---

The results of our experiment indicate that patchy disturbances create heterogeneity in the structure and temporal dynamics of attached algal communities, one of the key elements of patch dynamics theory (Pickett & White, 1985).

---

Changes after disturbance of the vegetation can be revealed through changes in structural parameters such as stem density, biovolume and species richness (Pickett & White, 1985).

---

Narrowing gaps (education, distribution, food production, holistic research, resource management, value) indicate culture-sustaining vitality. Education, income, and value gaps have widened in the past several decades (Odum and Barrett 2005). Books and classical papers have been written on the field and process of gap dynamics (Pickett & White, 1985).

---

A commonly used definition of disturbance is the one by Pickett & White, (1985): any relatively discrete event in time that disrupts ecosystem, community, or population structure and changes resources, substrate availability, or the physical environment.

---

Disturbance is a ubiquitous force in ecological systems that shapes patterns and dynamics across a range of spatial and temporal scales (White and Pickett 1985).

---

The dynamic nature of ecosystems has been studied in many ecological contexts, including: ecosystem change and vegetation succession (Pickett and Cadenasso, 2005); forest dynamics modeling (Loucks, 1981; Whitmore, 1982; Shugart, 1984); landscape mosaics (Levin and Paine, 1974; Pickett and Cadenasso, 2005), etc. In most cases these ecological studies focused on natural systems and natural disturbances (Pickett & White, 1985), with little or no reference to human intervention or impact.

---

Disturbances are one of the major causes of fluctuation in ecosystem structure and functioning because they act in promoting environmental heterogeneity and resources releasing (Pickett & White, 1985).

---

Intensity or timing of treatments, as well as land-use history and initial stand composition, all of which can influence how plant communities respond to disturbance (Pickett & White, 1985).

---

Ecological disturbance usually is understood as sustained disruption of ecosystem structure and function for periods longer than a single growing season (Pickett & White, 1985).

---

| |
|---|
| Many mature lots in the low BD areas offered similar moderate disturbance, especially if portions of the lots were neglected or had little maintenance. This variable patchwork offers Rich Patch Dynamics (sensu Pickett & White, (1985)). |
| Local controls and exchanges across patch boundaries may fundamentally influence processes taking place in adjacent ecological systems (Naiman et al. 1988) and dictate patch dynamics (Pickett & White, 1985). |

Excerpts of text citing Grubb (1977)

One important lesson from successional studies is that each species has a range of responses to the environment, depending on its life history stage (seed, seedling, juvenile, and reproductive adult) and whether the plant is colonizing, establishing, growing, or senescent (Grubb, 1977).

Grubb (1977) defined the regeneration niche as the biotic and abiotic species requirements to replace a mature individual. This concept encompasses most processes in the life cycle, from flowering to establishment and onward growth. The regeneration niche is defined here, in line with common usage, as the environmental conditions in the early phase in the life cycle of a plant (seedling and sapling stage).

Grubb (1977) suggested that the regeneration niche of a species may differ strikingly from its adult niche and that species differ in their adaptations to those niches.

It is expected that the importance of competition increases with succession because resources become limited as stand biomass increases (Grubb, 1977). Such increased competitive pressure will lead to differentiated strategies to obtain the increasingly scarce resources, with concomitant different functional traits, a process leading to increasingly limited trait similarity.

The regeneration niche theory (Grubb, 1977), which states that seedling establishment requirements differ among individual species, means that different availability of safe sites (sensu Harper et al. 1965) could determine establishment of individual species.

The repeated periods of grazing and treading by cattle, on the other hand, result in repeated periods of gap formation, which provide space and favorable light conditions for germination and seedling recruitment (Grubb, 1977).

More generally, coexistence could be maintained by a wide range of gap-phase related processes, including the generation of a mosaic of small-scale patches at different phases in succession (Jones 1946, Watt 1947, Forcier 1975, Connell 1978), species sorting along a gradient of gap size (Kohyama 1993, Busing and White 1997), within gap partitioning (Ricklefs 1977, Denslow 1980), the regeneration niche (Grubb, 1977), or different temporal strategies to access the canopy (Canham 1990, Poulson and Platt 1996, Messier et al. 1999).

Grubb (1977) proposed the concept of a regeneration niche, indicating that the diversity of strategies in the species regeneration patterns favors coexistence in plant communities.

Grubb (1977) reiterated the importance of the regeneration niche in community assembly.

This links with the regeneration niche concept (Grubb, 1977) which stresses the importance of plant establishment requirements to the control of the whole community dynamics.

Niche breadth is therefore affected by a species phenology and its regenerative niche (Grubb, 1977).

Species are at their most susceptible to climatic variation in their regeneration niche (Grubb, 1977).

Third, once established, trees have a greater range of environmental tolerance than seedlings (regeneration niche, Grubb, 1977).

---

Although germinant and seedling densities did not differ in their broad-scale patterns relative to the forest edge, seedling regeneration niche appeared to gradually shift as seedling fine-scale patterns and environmental associations progressively diverged with increasing size-class. Such changes in regeneration niche over time (Grubb, 1977) may lead to discordant spatial patterns of tree size-classes structured by different regeneration bottlenecks (Doviak et al. 2001, 2003; Gratzer and Raib 2004).

---

A central goal of ecology is to understand the mechanisms responsible for the coexistence of species, and niche differentiation during regeneration has long been regarded as an important explanation for the diversity of plant communities (Grubb, 1977).

---

Loss of coupling might be due to the differential effect of experimental treatment on the vegetation and diaspore bank. We can hypothesize three possible reasons for this decoupling: (1) the presence of perennial vegetation not dependent on seed germination, e.g., Calluna vulgaris and Vaccinium myrtillus; (2) creation of few establishment niches (sensu Grubb, 1977); (3) with time the effects of one-off treatments will diminish.

---

Gopher mounds may also serve as safe sites (Grubb, 1977) for species that are poor competitors in the undisturbed community.

---

There is good evidence in the literature that certain habitat qualities are necessary for seedling recruitment. The theory of a safe site, or regeneration niche (Grubb, 1977), is widely accepted. But as seen here, microsite quality plays an important role even one step earlier, by affecting the spatial pattern of seed distribution.

---

Absence of most seed bank annuals from the established vegetation may reflect unsuitable conditions for germination or establishment (regeneration niche, Grubb, 1977).

---

Grubb (1977) emphasised the regeneration niche as one of the important components of a plants niche. The significance of young life cycle stages for species dynamics and coexistence has been stressed by several authors in subsequent studies (Fenner and Thompson 2005; Leck et al. 2008).

---

Germination requirements are important factors that characterise the regeneration niche (Grubb, 1977).

---

The subtle differences detected in the preferential micro-habitats for germination and establishment of each species suggested a differentiation in their regeneration niche (Grubb, 1977).

---

Biotic and abiotic variables can act on plant functional traits to constrain species to a particular set of environmental conditionstheir regeneration niche (Grubb, 1977).

---

Our objective is to understand the mechanisms by which the first generation of trees regenerating in successional TDF (referred to hereafter as established trees) influences the regeneration of woody plants from the local species pool, and therefore secondary succession. We focus on the early and critical stages of the process of regeneration (sensu Grubb, 1977)seed dispersal, survival of seeds, germination, and seedling establishment (Poorter 2007) to answer the following questions.

---

"Exotic, invasive species frequently exploit this window of opportunity in the regeneration niche (Grubb, 1977)."

---

Although canopy removal initially increases light availability and decreases competition with trees, plant species richness is positively affected (Grubb, 1977).

---

These features are typical of disturbed areas, favouring the establishment of new species within a community or the renewal of the same community (Grubb, 1977).

---

The best microsite for S. bracteolatus seedling establishment during summer was in the shade of tussocks, but the optimal S. bracteolatus micro-environment changed as plants grew, as in many species as they mature (Grubb, 1977).

---

While these studies provide insight into detailed mechanisms of succession in established communities, they risk overlooking some fundamental processes of species colonization and establishment, which also structure communities (Grubb, 1977). Disturbances often create abiotic and biotic conditions that interfere with the re-entry of pre-disturbance dominant species by pre-empting the regeneration niche creating a priority effect and modify the successional trajectory.

---

Those plant adaptations include (1) seed structures to escape predation both from terrestrial mammals and from fish (Junk 1989) in order to germinate, (2) seedling strategies (Grubb, 1977) either of growing fast in order to have their leaves above the next high water level or of being able to endure extended submersion, and (3) special root structures, such as aerenchyma tissue, to facilitate gas exchange under water (Junk 1989, Lopez & Kursar 1999, Parolin et al. 2004a, Parolin 2009)

---

Regeneration is a central process of forest ecosystem dynamics (Grubb, 1977), and sustainable forest restoration is only possible if adequate information on regeneration of species is available.

---

The higher effectiveness of repeated treatments in driving compositional shifts may be due to the more intense disturbance produced by annual cutting, which suppresses the dominant P. aquilinum (Stewart et al. 2008) and creates regeneration niches after disturbance (Grubb, 1977).

---

Thus, we propose an extension of the nucleation model, the matrix discontinuity hypothesis, which predicts that the rate and pattern of succession will be determined not just by isolated trees, but rather by distinct and observable habitat patches or areas that provide clear regeneration niches (sensu Grubb, 1977).

---

Different disturbance regimes influence vegetation composition and characteristics by modifying the abundance of some existing species and providing establishment opportunities for others in different regeneration niches (Grubb, 1977), with different resource requirements, or with different life histories (McIntyre et al.1995; Moloney and Levin 1996).

---

The responses of plant communities to disturbances and their resulting changes over time in this study may be explained by the dynamic systems perspective (Roberts 1987; McCook 1994), regeneration niches (Grubb, 1977) and resource availability (McIntyre et al.1995; Moloney and Levin 1996) whereby changed environmental conditions following disturbance allow species most suitable to the altered environment to colonise and occupy the site.

---

The high level of small-scale heterogeneity in rain forests contributes to the local coexistence of species that share the same ecological niches in the adult stage but have different regeneration niches (Grubb, 1977).

Differences among plant species can contribute to coexistence and the occurrence of species-rich communities, and many important differences reside in how plants reproduce, i.e., their regeneration niche (Grubb, 1977).

Disturbances can create bare substrate in otherwise closed vegetation (Klinkhamer & de Jong 1988; Peart 1989), thereby creating potential regeneration niches for C. canescens (Grubb, 1977).

This is interesting as, apparently, the Corynephorus site, which was suitable for seedling emergence was not as suitable for seedling establishment, whereas the pioneer forest sites favoured seedling establishment to a much greater extent than emergence. Evidently, emergence and establishment requirements were quite different, highlighting the importance of the regeneration niche (Grubb, 1977).

In the dense vegetation of lowland communities, the importance of gaps for seedling recruitment has been experimentally shown: gaps reduce competition (Silvertown 1981; Silvertown and Smith 1988; Callaway et al. 2002) and provide regeneration niches (Grubb, 1977).

Indeed, organisms requirements and sensitivity to environmental factors change during their lifecycle (Grubb, 1977). Grubb (1977) and Young et al. (2005) suggested analysing ecological niches according to critical life-history stages of species (e.g. reproduction, dispersal, growth). Ontogenetic niche shifts are therefore a crucial issue (Werner and Gilliam 1984, Miriti 2006).

A number of authors have suggested that the ecological niche of early life stages is the main factor defining the species niche space and distribution (Grubb, 1977).

The seedling recruitment stage is a key component of the regeneration niche (Grubb, 1977) that determines the initial success or failure of a species to establish (Gibson and Good 1987).

Although the soil chemistry of most sand steppes in this region has been investigated (Olsson et al. 2009), this may not reflect the conditions required for the regeneration of sand steppe species as the regeneration niche may differ from that of species growing in established vegetation (Grubb, 1977).

The environmental conditions that allow new individuals to colonize an area successfully and become established in the local population has been defined as an individual's recruitment niche (Young et al. 2005), similar to Grubb (1977)'s regeneration niche. If the recruitment niche of an organism is parameterized according to its multiple environmental influences, trade-offs can be quantified and recruitment success predicted under multiple scenarios of biophysical forcing.

A regeneration niche, i.e. a competition-free environment, is necessary for seedling recruitment and establishment for species with diverse life-history tactics (Grubb, 1977).

The realized niche of ssp. alpini might be more restricted than its fundamental niche, since it survived on sites lower in the marsh than where it can be found in the field, suggesting possible exclusion by interspecific competition. However, it did not flower at the lower site and the experiment, by virtue of transplanting vegetative clumps, did not take into account the regeneration niches (Grubb, 1977) of either subspecies.

A variety of mechanisms underlying the productivity-richness relationship have been proposed and debated, including competitive exclusion (Grime 1973, 1979) or displacement (Huston 1979, Tilman 1982), colonization and recruitment of individuals (Grubb, 1977), niche differentiation and facilitation, niche complementarity, evolutionary adaptations (Silvertown et al. 1999, Cornwell and Grubb 2003), dispersal limitation (Prtel and Zobel 2007), and a no-interaction model solely based on plant density (Oksanen 1996).

The mechanisms by which epiphytic bryophyte communities form are poorly understood at least in part because of uncertainty about the applicability of successional theory derived from vascular plant systems. Individual investigations of the four aspects of the regeneration niche (dispersal, establishment, disturbance, and competition after Grubb, 1977) have shed light on notable dispersal limitations (Jonsson 1993, Ross-Davis and Frego 2004), challenges to successful establishment (Longton and Miles 1982, Kimmerer 1991, 2005), compositional shifts following substrate changes or disturbances (Barkman 1958, Studlar 1982, Sderstrm 1988a, Rambo and Muir 1998, der and van Hees 2004), and the apparent relative unimportance of interspecific competition for mature bryophyte gametophytes (Slack 1982, During and van Tooren 1987, Sderstrm 1988b).

Interspecific interactions such as competition and facilitation may reduce or increase the fundamental niche so that the realised niche (Grubb, 1977) differs from the fundamental niche.

According to Grubb (1977)'s definition, the realised niche of E. spinalba is an environmental space (with aspect and elevation as main axes) constrained by biotic interaction, here grazing pressure (herbivory).

Excerpts of text citing Tilman (1988)

| |
|---|
| The tradeoff between light and soil nutrient demand has been generalized as a governing principle of vegetation dynamics change in north temperate environments (Tilman 1988). |
| The dynamics of succession is strongly influenced by soil dynamics (Tilman 1988). |
| Secondary succession usually starts with relatively high soil organic matter content and nutrient availability, a well-developed seed bank and soil food web, and fast-growing, weedy plant species that are more limited by light than by nutrients (Tilman 1988). |
| Because of poor soil moisture and nutrients in the shoulder, the dominant plants of this microhabitat are expected to allocate more energy to below ground parts for acquisition of moisture and nutrients (Tilman 1988). |
| Optimum moisture and nutrient content in side slopes and removal of tall shrubs and trees by brushwood cutting may affect the above- and below-ground architectural characteristics of dominant plants in the side slope. Removal of shade providing shrubs and trees encourage the dominant plants of the side slope to maximize their above-ground spread for capturing light resources (Tilman 1988). |
| High rootshoot ratio of plants indicates poor site conditions in terms of soil resources, as the plants allocated more energy to their below ground parts to ensure maximum acquisition and storage of soil resources (Tilman 1988). |
| Similar examples of lower light transmission on richer soils have been reported in other parts of the world (reviewed by Coomes & Grubb 2000), for example, along gradients of sandiness in central-northern United States (Tilman 1988) and in tropical rain forests on contrasting soil types in Malaysia (Palmiotto et al. 2004) and Amazonia (Coomes & Grubb 1996). |
| The resource-ratio hypothesis (Tilman 1988) provides a theoretical framework for understanding why competition for light is weak on nutrient-poor soils. Tilman (1988) argued that plants allocate a larger proportion of assimilate to foraging for nutrients when growing on poor soils and consequently produce relatively few leaves, resulting in a shift from above- to below-ground competition. |
| The dominance by exotic annuals and their replacement by native perennials appear to represent the transient dynamics that are common in temperate regions (Tilman 1988). |
| Species that are subordinate in natural communities might be able to colonize and grow faster than the corresponding dominant matrix species (cf. colonizationcompetition trade-off; Tilman 1988). |
| Contrasts of succession across entire severity gradients of nutrients or other resources (e.g. light or water availability, substrate stability) could provide helpful comparisons among disturbance types and test general principles of disturbance response (Tilman 1988). |
| In environments with scarce water and nutrients, pioneer species are highly tolerant to disturbance, but are poor competitors when resource availability is high (Tilman 1988). |

| |
|---|
| Vegetation is a critical habitat component for most animals, acting as a food resource for herbivores, and more generally as a source of shelter from predators and from the vagaries of weather (Partridge 1978; Huey 1991). It is also a very dynamic habitat component with rapid changes occurring, for instance, after a fire (Catling & Newsome 1981), while more gradual changes can occur through plant growth and succession (Tilman 1988). |
| Autonomous patch processes, such as the increase of intraspecific competition in denser patches (Tilman 1988), may influence the clonal diversity. |
| Traditional theories predict that in the absence of disturbances high, soil productivity should produce late-successional forests of shade-tolerant tree species (Tilman 1988). In addition, these theories predict that species adapted for surviving and/or competing well in unproductive soils (i.e., stress-tolerators of Grime [1979]; belowground competitors of Tilman 1988) are at a competitive disadvantage in productive soils. |
| The community dynamics we observed do not necessarily reflect equilibrium outcomes for competition among the species, nor do they reflect the influences of AMF (arbuscular mycorrhizal fungi) on species richness via effects on plant recruitment (van der Heijden 2004). Rather, we may be seeing transient dynamics that are strongly influenced by differential growth rates among species (Tilman 1988). |
| In general, fertilization increases biomass of grasslands (Tilman 1988). |
| This may be due to ongoing adaptations of species to the available resources when annual nutrient removal by mowing does not lead to a decrease in nutrient availability (Tilman 1988). |
| After disturbance, some communities can rapidly return to the original state, usually following the tolerance (i.e. fast growing and high dispersers are the first to colonize the disturbed zones but are replaced by slow growing competitively superior species over time; Tilman 1988). |
| The answer to the first question may be that high tall runner biomass was not sustained for long enough in the first part of the experiment to decrease non-tall runner biomass and species richness. Transient dynamics can occur whereby inferior competitors persist until resource levels equilibrate (Tilman 1988) or disturbance ceases (Huston 1979). |
| Competition results when plants share limited resources. Inter-specific competition is clearly influenced by geoclimatic conditions and plants can respond using two different strategies. Firstly, plants may choose faster growth to occupy more space and, as a result, improve their chances of obtaining new resources (Grime, 1977). On the other hand, another option is to survive with lower levels of resources (Tilman 1988). |
| Net Nitrogen mineralization rates vary depending on the changes of vegetation type, decreasing during natural secondary succession in old-fields [15], but increasing during secondary succession on infertile sandy soils Tilman (1988), and in nutrient-poor ecosystems [17]. |
| Changing resource supplies (light, but also soil resources) along a density gradient can also indirectly alter the competitive balance between species (Tilman 1990; Wagner & Radosevich 1998; Kaelke et al. 2001; Lusk 2004; Baraloto et al. 2005) as different resource environments may favour different competitive strategies or plant traits (Tilman (1988); Suding & Goldberg 2001; Baraloto et al. 2005). Examples include the hypothesized trade-offs between above-ground and below-ground competitive ability across fertility gradients (Tilman 1988). |

| |
|---|
| ”The energy gain of every time step is calculated as follows (see Tilman 1988):<br>Egain=min(lightgain , soil resourcegain)plant(modulerespirationmodule)<br>The resource availability in the ecological field is calculated at a 1 cm scale. There is no shade tolerance (Lavorel, 2001; Warren and Topping, 2001) incorporated in the model, all species have the same ability to use light.” |
| Plants are believed to allocate a relatively high proportion of biomass to leaves in nutrient-rich environments (Tilman 1988), and a low proportion of biomass to leaves in nutrient-poor environments (Chapin et al., 1987). |
| Long-term studies offer very valuable information when studying the effects of disturbance because of the transient nature of the initial species responses (Tilman 1988). |
| Tilman (1988)’s resource-ratio hypothesis proposes that vegetation dynamics over time is characterized by a change from competition for nutrients to competition for light. Four different hypotheses of classical and current models of species distribution and biotic interactions are addressed: (1) along a productivityclimatic gradient, the regeneration of Q. ilex (the xerophytic species in this study) varies only with competition, i.e. it can colonize sites where its direct competitors suffer from increasing drought (Tilman 1988); (2) shrub encroachment can facilitate Q. pyrenaica (the mesophytic species in this study) in its xeric margin of distribution (Sthultz et al. 2007); (3) traits specific to nurse shrubs can have different effects on the regeneration and relative proportion of oak trees (Chu et al. 2009; Padilla et al. 2009); (4) both the facilitative role of shrubs on Q. pyrenaica and their negative effects on Q. ilex could alter the relative proportions of these two species, locally modifying the location of the bioclimatic limit along the elevation gradient. |
| Further, the clear allocation trade-off in favor of clonal reproductive biomass at the expense of root growth in H. pilosella may suggest that in terms of carbon investment the AMF-dominated foraging strategy is cheaper than root growth (Fitter 1991; Schweiger et al. 1995) and therefore allows for enhanced Carbon allocation into vegetative reproduction in order to form the well-known closed clonal aggregations of H. pilosella (Bishop et al. 1978; Bishop and Davy 1994), which minimize interspecific competition (Tilman 1988) at the community scale. |
| Local increases in productivity produce a shift in the limiting factors for plant growth, from soil resources to light, thus increasing the intensity of aboveground competition (Tilman 1988). |
| Consistent with the view that a colonizationcompetition trade-off is core to succession processes (Tilman 1988) and results from many studies of succession (Tilman, 1990), species with early-successional traits (i.e. annual life cycle, reproduction by seeds, small seeds) dominated as colonists in species-poor newly established experimental communities, while mid-successional traits (perennial life cycle, taller growth, vegetative reproduction) characterized later colonists and those in communities sown with greater diversity (Roscher et al., 2015). |
| One of the many theoretical mechanisms for succession is the resource-ratio hypothesis, whereby the timing of establishment by certain plant species is dictated by their specific requirements for limiting resources (e.g., soil nutrients, water, light), the availability of which change over time (Tilman 1988). As these limiting resources fluctuate over time, total plant biomass increases; however, if the rate of accumulation of these resources is slow, succession occurs as a slowly shifting trajectory for plant communities, which is determined by the resource availability at that point in time (Tilman 1988). |
| This is consistent with the resource-ratio hypothesis of succession, whereby changes occur on a gradient over time relative to the changing availability of limiting resources at the soil surface (Tilman 1988). |

Excerpts of text citing Egler (1954)

The observed change in dominance in the first few years following abandonment was best explained by differential growth rates among species and likely some suppression by faster-growing species (Egler (1954) initial floristics composition model).

It is commonly known for long time that secondary succession is affected by the initial presence of propagules (Egler, 1954). Consequently, addition of propagules by sowing is an important management tool to affect the course of succession.

Sowing considerably affected the course of succession and the effect of sowing persisted for the whole experimental period. These results support Egler (1954)'s well established initial floristic composition hypothesis.

The second mechanism of differential species availability is the persistence of propagules in the soil or substrate of the disturbed site (Egler, 1954).

The Initial Floristic Composition (IFC) hypothesis of Egler (1954) is predicted to apply to tropical postagricultural secondary succession where previous land-use was of low intensity and seed sources are nearby (Gomez-Pompa & Vazquez-Yanes 1981, Finegan 1984). According to the IFC hypothesis, most species that will later dominate the community will colonize the site at the onset of succession. The proposed sequential physiognomic dominance of species, first light-demanding species of increasing longevity, and finally shade-tolerant species, unfolds largely due to differences in growth rates, longevity, and shade-tolerance among tree species that happen to colonize the site at abandonment (Gmez-Pompa
Vazquez-Yanes 1981, Finegan 1984).

We evaluated the prediction that the IFC hypothesis (Egler, 1954) applies to succession on abandoned agricultural fields with light use and proximity to old-growth forests patches (Gmez-Pompa & Vzquez-Yanes 1981; Finegan 1984, 1996).

Therefore, the initial floristic composition hypothesis (Egler, 1954), which predicts that early processes largely determine the outcome of restoration, may explain short-term changes following grassland restorations, but is not supported by longer-term experiments (Table 1).

Even so, we assumed that different rates and pathways of vegetation development on sites with and without species introduction will have a long-term influence on the composition of communities (Hypothesis 2), as species introduction will influence seed availability and dispersal stochasticity (e.g. Zobel et al. 2000), initial floristic composition (Egler, 1954) and the sequence of species arrivals (e.g. Drake 1991).

A continuous decrease in diversity during succession, caused by prevalence of extinction processes, is consistent with the hypothesis of initial floristic composition (Egler, 1954). At the beginning of succession a large number of species immigrate, but more and more species go extinct as competition for increasingly depleted resources, mainly light, increases over time. In addition, stochastic extinction increases when individuals increase in size and the overall density of individuals declines.

In their study of comparable forests in Zhejiang Province, Li et al. (1999) hypothesized that species composition in subtropical forests in China is primarily driven by initial floristic composition (Egler, 1954). Our results support the view that many woody species arrived early in succession. Similarly, the finding that there were only a few species specific to any particular successional stage lends support to the prevalence of initial floristic composition (Egler, 1954).

In accordance with Egler (1954)'s initial floristic composition hypothesis, individuals from both pioneer and mature forest species established at the onset of succession (cf. Pea-Claros, 2003); however, only the pioneer species were present in high numbers, and displayed rapid growth rates (Figs. 2df and 4e, g). The reduction in recruitment and the increase in mortality of pioneers occurred simultaneously, and correlated strongly, with their own structural development (Figs. 2 and 4, mid-panels).

| |
|---|
| The concept of initial floristic composition of Egler (1954) suggests the colonisation of early and late successional species in initial stages. The tree species that dominate today on our plots, above all Betula pendula, Salix caprea and Fraxinus excelsior, already germinated in the first year of abandonment and thus affirm this theory. |
| This pattern is reminiscent of the old concept of initial floristic composition (Egler, 1954), which states that all species are present at the start of the succession and are then differentially selected. An increase in trait similarity with stand age is generally interpreted as an effect of environmental filtering, with a progressive sorting of the species best adapted to the local abiotic conditions (Uriarte et al. 2010). Thus, succession in SDTFs (Seasonally dry tropical forests) may not follow the relay floristic model (Egler, 1954), which predicts a gradual substitution of pioneer by late species along forest recovery. Instead, these ecosystems may conform to the initial floristic composition model, with pioneer species remaining in advanced stages of succession (Egler, 1954). |
| We tested the following hypotheses about ecological succession in the studied SDTF: (i) successional dynamics is dominance controlled. In this case, tree diversity would be higher in intermediate stages of succession, due to the competitive exclusion of mid-successional species as the forest matures (Yodzis 1986; Begon et al. 2006). (ii) Succession pathways conform to the initial floristic composition model (Egler, 1954). |
| The relative importance of seed colonization and resprouting in forest recovery is controversial, originating opposing succession models. In the relay floristics model, a gradual species substitution is expected across time, whereas the initial floristic composition model predicts that pioneer species remain in advanced stages of succession (Egler, 1954). The successional gradient analysed in the present study corroborates the former model, since there are striking changes in tree community composition from early to intermediate and late stages. |
| Classical successional theory (Clements 1916) suggests that community composition should converge toward climax, as determined by environmental conditions (the monoclimax theory). In contrast, Egler (1954)s Initial Floristic Composition concept predicts that the course of succession is, to a large extent, determined by the species that were initially dominant; thus we would expect divergence. |
| As a consequence, we observed pronounced and relatively fast convergence within habitats, i.e. in individual habitats the similarity of communities was very high at the end of the study period. In contrast, the effect of time of succession initiation (both season and year) was very pronounced at the initial stages, but decreased with time. This implies that the initial floristic composition was, in our case, less important than suggested by Egler (1954)s initial floristic composition concept. |
| This class includes not only species characteristics, but also landscape context as a source of propagules and the characteristics of the dispersal vectors that move seeds. The importance of this broad category to constraining succession is highlighted by the concept of initial floristics where succession is determined by initial colonists (Egler, 1954) and is critical to neutral theory, which focuses on stochastic dispersal and colonization events (Hubbell 2001). |
| Some early conceptual models (Egler, 1954) and subsequent research demonstrated that dispersal limitation can be a key determinant of the rate of succession. The IFC hypothesis states that species representing all guilds colonize soon after disturbance but reach dominance at different times according to their growth rates and longevities (Egler, 1954). |
| A long-standing debate in the succession literature, that has primarily been tested in the temperate zone, is whether forests are initially colonized by a group of pioneer species that are subsequently replaced by later successional species (the relay floristics modelEgler, 1954) or whether the species that initially colonize the site remain later in the successional process (initial floristic composition) (Egler, 1954). |

| |
|---|
| We found that perennial species occurring with high cover already in young old-fields were more successful in late-succession old-fields in both regions than those detected only with low cover in young fields. This is well in accordance with Egler (1954)s theory of initial floristic composition and draws attention to the importance of the founder effect (Egler, 1954). |
| Many adaptations to fire (e.g. serotiny, resprouting, soil seed-banking, thick bark) exist among species in this ecosystem; therefore species diversity was expected to be high immediately following fire. Early post-fire species were expected to have a dominant effect on subsequent succession, supporting the initial floristics model (Egler, 1954). |
| Consistent with the initial floristics model (Egler, 1954) and findings in other coastal conifer forests in mediterranean climates (Capitanio<br>Carcaillet 2008; Franklin 2009), species diversity and richness peaked 1 and 2 yr following fire, respectively, due to the presence of early- and late-successional species. |
| Classical succession has the key assumption that species are replaced because they modify the environment so that it becomes more suitable for other species but less suitable for themselves. This was termed the facilitation model of succession by Connell and Slatyer (1977), who also coined the term inhibition model for Egler (1954)'s hypothesis of initial floristic composition that followed from Gleason's (1917) initial questioning of the view of Clements (1916) |
| Thus, most burned conifer forests quickly recovered prefire species composition and tree density, not through classic succession or relay floristics but via direct re-growth or initial floristic composition sensu Egler (1954). |
| Egler (1954) had proposed that many of the species that would come to predominate in later successional communities were in fact present right from the start. His Initial Floristic Composition hypothesis was in opposition to the dominant assumption that species arrived in succession in order of their dominance. |
| The initial floristic composition is a significant factor in old-field succession, since dominants of later stages usually arrive soon after abandonment and early dominants substantially influence the further vegetation development (Egler, 1954). |
| The observed gradient strongly supports the initial floristic composition model by Egler (1954). |
| Our findings clearly confirm the initial floristic composition model by Egler (1954). Differences caused by the initial treatments at the beginning of the experiment fundamentally determined the initial floristic composition and the subsequent development of vegetation on the plots. |
| We can examine whether species establish differentially along succession according to their life history attributes (cf.Clements 1916), or alternatively whether species from different functional groups colonize open spaces synchronously (cf. Initial Floristic Composition hypothesis, IFC; Egler, 1954). |
| The initial stages of primary succession are particularly important as they can have a profound influence on the subsequent development of the vegetation. The pre-emption of space by early colonists can have a lasting impact on species composition Egler (1954) |
| Other seedlings and saplings present in Euakafa20 in 2005 mainly comprised gap-establishing species, but also included some shade-tolerant species in low abundance. This is consistent with Egler (1954)'s initial floristic composition model of forest succession, as modified by Finegan (1996), who stated that most, but not all, forest species establish within a few years of disturbance (e.g. Uhl<br>Jordan 1984). |
| However, our results are more in line with the Initial Floristic Composition Theory (Egler, 1954): late-successional species were present in the area at the start of the succession and most of the species recorded had colonized early. |
| Priority effects or local conditions may be important and might promote continuing heterogeneity among communities (Egler, 1954). |

| |
|---|
| There is little consensus in how species richness and diversity change over successions (Huston 1994; Rosenzweig 1995), with various models predicting richness and diversity peaking at either early (e.g. the initial floristic composition model of Egler, 1954) or intermediate (e.g. the intermediate disturbance hypothesis of Connell 1978) stages of succession before declining as interspecific competition strength increases. |
| Thus, it is likely that some true forest species were already in place at the onset of succession, i.e. the initial floristic composition model of Egler (1954) holds relevance to these data, and that the only evident process during successional development is a loss in non-forest species richness. |
| Although the amelioration of adverse conditions ceased three years after community establishment, and all plots were subjected to the same environmental conditions thereafter, initial differences in species composition persisted in time. Such results are consistent with the hypothesis of initial floristic composition (Egler, 1954), which states that those species present at a particular site early in succession pre-empt the site and influence the course of succession on it for a long time. |
| Initial and relay floristics models of succession (Egler, 1954) are useful for understanding early succession in managed forests because these models (1) focus on patterns of initial regeneration that may be evident in the immediate post-disturbance phase of succession (Noble<br>Slatyer 1980), (2) predict measurable shifts in diversity and abundance, and (3) are sufficiently descriptive to be useful for developing forest management strategies (Egler 1954; Kimmins 1997). |
| The data support the initial floristic model of succession (Egler, 1954) among structural layers, shown by constant species richness per plot in upper structural layers coupled with a shift in the dominance of structural layers over time. |
| The potential for initial and relay floristics to act in concert is consistent with Egler (1954)'s concept of vegetation development, and is indicative of the importance of the disturbance regime in succession. |
| The species response curves presented here provide no evidence to support the hypothesis that late-successional (e.g. Betula spp.) or bog-forming species (Sphagnum spp.) will invade the moorlands if prescribed burning was not carried out. The results are consistent with the initial floristic model (Egler, 1954) and the tolerance models of succession working together through time (Connell & Slatyer 1977). |
| Alternative Successional Treajectory (AST) can come about in at least two ways: (1) through competitive effects of certain colonizers (initial floristic composition, sensu Egler, 1954) that inhibit the recruitment of other species and (2) through local extirpation of a key constituent of the normal successional trajectory. |
| The succession observed here did not involve complete replacement (relay floristics), but resulted at least in part from a shift in dominance (initial floristics sensu Egler, 1954), with the persistence of a group of fungi throughout the stand age sequence. |
| Examples of theoretical concepts include the intermediate disturbance hypothesis (Connell 1978), general successional theory (Luken 1990; Johnson and Miyanishi 2008), habitat accommodation theory (Fox 1982), the initial conditions hypothesis (Egler, 1954), ecological thresholds (Groffman et al. 2006), resilience thinking (Walker and Salt 2004), and biological legacies (Franklin et al. 2000). |
| Some plant communities are thought to change according to the initial floristic composition model of succession (Egler, 1954). Under this paradigm, diversity declines with increasing time since the last fire, as all (or at least the majority of) plant species present during the succession re-establish shortly after fire, re-establishment is relatively rapid, and changes over time are driven by plant life histories; growth rates, growth forms and survivorship (Collins et al. 1995; Capitanio & Carcaillet 2008). |
| The density of regenerating buds would, therefore, set a medium-term limit to shrub cover expansion as long as there was a limit to the cover and size of individual shrubs under the study conditions. The importance of the initial composition of the vegetation in determining vegetation dynamics is well established (Egler, 1954). |

| |
|---|
| The initial floristic composition (IFC) hypothesis (Egler, 1954, see also van Breugel et al. 2007) is applicable to tropical post-agricultural succession where previous land use is of low intensity and seed sources are available in nearby natural forests, conditions typical of abandoned forest villages. The hypothesis proposes a sequential floristic or life form dominance of species. Early after abandonment, light-demanding species will dominate, but will eventually be replaced by shade-tolerant species due to differences in growth rate, longevity, and shade-tolerance among tree species that colonize an abandoned site. |
| Since regeneration under trees in pastures constitutes a very early step in forest recovery, evaluating the importance of attributes of isolated trees for the structuring of the regeneration assemblage will enable assessment of the importance of the legacy of the initial tree composition (Egler, 1954), and of the deterministic and predictable character of succession. |

| |
|---|
| Direct, repeated observations (e.g. through historical photography or long-term plot studies; del Moral 2007) began formally with studies of dunes in Denmark (Warming 1895) and Michigan (USA; (Cowles, 1899)), and such observations provide the best source of evidence about temporal changes in plant and soil biological communities over years to decades. |
| In this paper, we review these classic examples of chronosequence-based succession and the studies that have tested the chronosequence assumptions and inferences. These classic examples are: Cowles (1899) study of sand dune succession, Dachnowskis (1912, 1926a) study of hydrarch succession, Coopers (1923a,b, 1931, 1939) and Crocker<br>Majors (1955) studies of succession on till substrate following glacial retreat, and Billings (1938) and Oostings (1942) studies of old-field succession. |
| Although preceded by earlier studies of vegetation succession on coastal dunes (e.g. Beck-Mannagetta 1890; Warming 1895), the classic study of dune succession widely cited in English-language (especially American) textbooks is that by Cowles (1899), who examined plant communities on sand dunes along the southern shore of Lake Michigan. As the post-glacial lake receded over time, it resulted in the formation of a sequence of sand dunes representing former beach ridges. |
| The vegetation sequence for dune succession generally presented in textbooks (e.g. Fig. 1) tends to show a simple linear successional sequence of annuals, sand-binding dune grasses, cottonwoods, pines, and oak, despite the fact that Cowles (1899) had emphasized that only perennial dune grasses, shrubs, and trees such as cottonwoods were dune-forming plants (with cottonwoods germinating in protected depressions on the upper beach, p. 182) and had described different successional pathways for different dune locations (e.g. windward vs. lee slopes). |
| However, not only Cowles (1899) but numerous other studies (Fuller 1912; Downing 1922; Weaver<br>Clements 1929; Olson 1958; Poulson 1999) have reported that the cottonwoods establish only on moist germination beds such as depressions on the beach, low pannes, swales, or recently in-filled runnels, all with surfaces close to the water table. This species does not successfully colonize dunes previously established by Ammophila. |
| Finally, the argument of facilitated dune succession leading to the directional progressive change from dune grasses to mesophytic forest was based on hypothesized changes to the light conditions and sandy soil brought about by each successive dominant species (Cowles, 1899). In particular, it was hypothesized that the plants changed soil properties (such as field capacity, pH, and base saturation) in a way that facilitated successful establishment by the next seral dominants. |
| Interestingly, a careful reading of Cowles (1899) original observations (not viewed through the lens of succession) suggests the actual processes and mechanisms determining species composition of the dunes. He noted that [p]erhaps no topographic form is more unstable than a dune and on the whole the physical forces of the present [italics ours] shape the floras as we find them (p. 96). |
| As long recognised since at least Cowles (1899), many real communities are in a transient, not stable, state, because disturbance keeps communities from reaching a stable state (reviewed in Pickett<br>White 1985). |
| Investigations of vegetation on sand dunes bordering Lake Michigan, USA (Cowles 1899) played an important role in introducing the ecological concept of succession, which was further developed by Clements (1916). |
| Studying succession with the chronosequence approach requires relatively short time, therefore it is a popular method, but may result in errors in the prediction of successional series. For example, a dune series at Lake Michigan suggested by Cowles (1899) has been disproved by Olson (1958) and Jackson et al. (1988) or the traditionally accepted prairie succession path was later confuted by Collins and Adams (1983) |
| Many examples of succession use species-abundance data from different locations that presumably represent different successional states, or chronosequences (e.g., shifting sand dunes, Cowles 1899; intertidal boulder fields, Sousa 1979; abandoned agricultural fields, Kardol et al. 2006). |

| |
|---|
| Some of our results concerning the influence of surface and substrate characteristics on vegetation structure (H2) match basic assumptions of successional theory such as the decrease of annuals as well as the increase of grasses and especially woody species in the course of time (Cowles 1899). |
| Odum's concept originated from the early idea of succession proposed by Cowles (1899), which realized that biological and physical components of ecosystems co-adjust in time and space. |
| Succession on various sand dunes has been studied in many parts of the world as one of the best examples of spontaneous primary succession (Doing 1985; Burrows 1990; Lichter 1998; Walker del Moral 2003). It was among the first successions described and successional theory has one of its roots in sand dunes (Cowles 1899). |
| The vegetation patterns and processes of coastal dunes have been of interest to ecologists for more than a century. In fact, as early as 1967, MacArthur & Wilson (1967) recognized the basic idea that plant communities of different ages in sand dunes reflected how the communities changed over time. |
| Primary succession often takes so long that it can only be estimated by comparing different-aged substrates, such as chronosequences of sand dunes. Cowles (1899) first suggested the relationship between the age of sand dunes and their overlying vegetation along the southern edge of Lake Michigan (Cowles 1899). |
| The functional role of dune vegetation as a management tool for altering the topography of a landscape has been well-documented by ecologists, beginning with Cowles (1899), and it is now well known that the formation of coastal dunes is closely associated with vegetation. |
| After an initial period of pioneer studies clearly influenced by Cowles (1899)s theory of ecological succession on sand dunes, Cooper (1936) was the first to report the distribution of 53 representative coastal plants from SW Alaska to northern Baja California. |
| Clonal taxa are important colonizers of man-made disturbed sites in Europe (Prach and Pysek 1994, 1999), of accreting shorelines of the Great Lakes in the United States (Cowles 1899), and of habitat exposed by glacial retreat (Stocklin and Baumler 1996) |
| . Great Lakes dune plant communities are classic models of primary succession (Cowles 1899). |
| More than a century ago, Cowles (1899) noted that the jack pine association was replaced overtime by black oak (Quercus velutina Lam.), an observation supported by later authors (e.g., Olson, 1958; Menges and Armentano, 1985; Lichter, 1998). |
| Nonetheless, the enhanced seedling survival near wet pannes in our experiment still guides us to explain the observations of Cowles (1899) of dense populations of jack pine in the intradunal depressions where the water table was high. |
| The increased dominance of black oak in the older dune ridge plots (Fig. 4, Table 3) supports the succession trajectory from jack pine to black oak that was predicted by Cowles (1899). |
| In a now classic study of sand dune succession, Cowles (1899) described the chronosequence, or space-for-time substitution of vegetation along sand dunes, moving from bare sand beach, to grasslands, to mature forests. From Cowles (1899)s and others studies of chronosequences over the past 100 y, ecologists have observed that plant species richness and diversity generally increase over successional time, especially for primary successional systems (e.g., Lichter, 1998), or early in secondary succession (e.g., Shafi and Yarranto, 1973; Cook et al., 2005). |
| Chronosequences have also been used extensively in ecology, and were in important in, e.g., development of some of the earliest and most influential ideas on ecological succession (Cowles 1899). |
| Sand dunes are pioneer sites for ecological succession [29] and they can be used as model systems to study the process of ecosystem development (Cowles 1899). |

Excerpts of text citing MacArthur & Wilson (1967)

| |
|---|
| Heggaton was the smallest of the locations ( Fig. 2), raising the possibility that reduced species richness may be a product of higher extinction rates on this smaller habitat island (MacArthur & Wilson 1967). |
| Larger rain-forest fragments have higher diversity of plants and animals than smaller ones (MacArthur & Wilson 1967) and thus should be greater source pools for recolonization of adjacent patches. |
| The theory of island biogeography predicts that species richness is a function of immigration and extinction rates, and that increasing isolation (i.e., distance away from sources of potential colonists) will decrease species richness via reduction in immigration rates (MacArthur & Wilson 1967). |
| While this is not what we expected to find, it does correspond with theory of island biogeography (MacArthur & Wilson 1967), which predicts an equilibrium species richness but nonequilibrium species composition due to local extinction and new immigration. |
| We note that the negative relationship between island area and species richness present in this study system is inconsistent with predictions of most neutral diversity models, such as the theory of island biogeography (MacArthur & Wilson 1967), which makes it possible to use this study system to investigate other mechanisms that may control diversity without confounding effects of island area. Further, dispersal limitation does not affect access of plant species to islands, and all plant species present in the system are present on even the smallest and most isolated islands. |
| ”The number of species was assumed to depend on the area of the grassland component of the old fields according to Arrhenius species-area relation:<br>$logS = c + z * logA$ (1)<br>where S is the number of species, A is the area and, c and z are constants (MacArthur & Wilson 1967).” |
| To explain this theoretical prediction, it helps to consider two contrasting models of community assembly. One is the classic theory of island biogeography (MacArthur & Wilson 1967), in which the species pool is a large stable reservoir of species unaffected by the dynamics of local communities. |
| Regional processes, on the other hand, can result from evolution through time (Jablonski and Sepkoski 1996, Ricklefs 2004), or at ecological time scales through dispersal to the local community from a larger region (MacArthur & Wilson 1967). |
| Stable plant communities are characterised by a balance between colonisation and extinction rates (MacArthur & Wilson 1967). |
| Underlying our hypotheses, H1H5 is the r/K selection theory (MacArthur & Wilson 1967) according to which species can be characterized as r selected (colonizers) or K selected (competitors) based on their life history strategies. The r selected species allocate a higher share of their resources to reproduction than K selected species, which are in turn often more long-lived and more combative. |
| The trade-off here resembles that of K and r strategies (MacArthur & Wilson 1967). An example of a species with K selected characteristics is P. nigrolimitatus, which produces fruit bodies only long after mycelial colonization. Those fruit bodies can be large, especially if the species has achieved a dominating position in the mycelial community, and they may remain reproductive for many years, possibly even for decades. The species Postia caesia exemplifies the r strategy. This common generalist species has small annual fruit bodies, and it never reaches a high total hymenophore area on a log. |
| Indeed, many of our most fundamental ecological models and concepts, such as island biogeography (MacArthur & Wilson 1967), metacommunity dynamics (Leibold et al. 2004), succession (Clements 1916), neutral models (Hubbell 2001), and community stability (Ives and Carpenter 2007) are based on temporally dynamic phenomena, yet they are frequently tested with spatial rather than temporal data. |

| |
|---|
| This conceptual approach of intact forest fragments immersed in a degraded matrix is based on the theory of island biogeography of MacArthur & Wilson (1967). We can also expect an important role of edge effect because of the relatively small areas of the spoil heaps, which is in concordance with the theory of island biogeography (MacArthur & Wilson 1967). |
| One reason why we found such an effect in this study may be that the urban environment constitutes an entirely hostile matrix, as assumed by metapopulation (Hanski, 1999) and theory of island biogeography (MacArthur & Wilson 1967), whereas in many other studies this may not be the case (Hanski and Pyry, 2007). |
| These bionomic traits are included among the descriptive traits which discriminate the ecological strategy adopted by organisms, according to the rK model introduced by MacArthur & Wilson (1967). Individuals of small size, high fecundity, high mortality rates, low structural organisation and investment in offspring often show a low competitive ability and therefore they have an ecological advantage in unoccupied or unstable environments (the r-strategy of colonisation). This type of organism dominates the earlier stages of an ecological succession, whereas later stages see the progressive achievement of highly competitive species, which mostly possess large individual size, longevity, high organisation level, efficient mechanisms of defence and predation (the K-strategists). |
| theory of island biogeography (MacArthur & Wilson 1967) and metapopulation dynamics (Hanski 1998) are key theoretical frameworks underpinning most ecological and conservation studies on species extinction (Ricketts 2001). |
| theory of island biogeography processes (MacArthur & Wilson 1967), involving stochastic gains and losses of species, would therefore also be expected to play an important role in determining species assemblages in clusters. |
| These results, and the increase in the number of species with cluster size, are consistent with interpreting clusters as biogeographic islands of different sizes (MacArthur & Wilson 1967). |
| When the distance between patches decreases, functional connectivity usually increases (Honnay et al. 2002) and thus equalizing forces on patch composition occur. Such processes have been extensively studied within the framework of island biogeography theory (MacArthur & Wilson 1967) and metapopulation dynamics (Hanski 1999), both assuming that suitable habitat patches are isolated from one another by hostile habitats (but see Laurance et al. 2007). |
| Also, study system is not directly comparable with the MacArthur & Wilson (1967)s dynamic equilibrium model developed in insular systems (e.g., Azeria 2004) because it was a closed system after the initial colonization phase and that local extinctions (if any) cannot be countered by recolonizations. |
| In recent years, island ecology has been dominated by theory of island biogeography based on MacArthur & Wilson (1967), which posits that the number of species on islands results from the balance between immigration and extinctions the former influenced by the distance from the mainland and the latter by the effect of island area acting on population size. |
| Size-related traits have been strictly associated with the ecological strategies adopted by organisms during the course of ecological successions in the classical rK model ( MacArthur & Wilson 1967). |
| Theory of island biogegraphy (e.g., MacArthur & Wilson 1967) attributes these phenotypes to environment-dependent selection that favors the greater productivity of weaker competitors in uncrowded, resource-rich sites opened by disturbance, but favors the greater efficiency with which stronger competitors convert resources to reproductive output as crowding and competition increase with succession. |
| MacArthur & Wilson (1967)'s theory of island biogeography was revolutionary, and also inspired the more recent unified neutral theory of biodiversity and biogeography. The unified neutral theory has the potential to make predictions about island biogeography that are not well studied. |
| Our model, in accord with the simplest version of MacArthur & Wilson (1967)'s theory, predicts linear immigration and extinction rates as functions of species richness at dynamic equilibrium. |

| |
|---|
| The central concept of the theory of island biogeography was a dynamic equilibrium in species richness where immigration of new species was balanced by local extinction. Under the theory of island biogeography, both immigration and extinction rates on islands are functions of island species richness, assumed linear in the simplest case, but allowed to be concave in other versions (MacArthur & Wilson, 1967). |
| The immigration curves are in accord with MacArthur & Wilson's theory, having a curved and decelerating shape for mainland species immigration probabilities following a difference log-series. This is caused by the species absent from the island tending to be the ones that have a lower chance of getting there, as proposed by MacArthur and Wilson. |
| The lower species richness at breast height on small islands could potentially result from small islands having fewer trunks of B. pubescens and thus a lower total area available for the colonization and establishment of lichens, as predicted by speciesarea theory (MacArthur & Wilson 1967). |
| The strong species-area associations observed suggest that clearcuts constitute habitat patches for the studied groups, situated in a matrix. This is consistent with the assumptions of the theory of island biogeography (MacArthur & Wilson 1967). |
| Life history traits are generally thought to be a zero-sum game of trade-offs (MacArthur & Wilson 1967) where the resource allocation to spore production, and the thereby attained high dispersal and colonization potential, are followed by a lowered competitive and persistence potential. |
| Small patch size and low connectivity act as dispersal filters as they strongly decrease colonization probabilities of species that have ineffective dispersal strategies under current landscape configuration and management (Kolb & Diekmann 2005; Lindborg 2007; Schleicher, Biedermann & Kleyer 2011). The lower propagule pressure on such patches should result in smaller species numbers (MacArthur & Wilson 1967). |
| During and after the process of deforestation, a forest remnant undergoes changes because it becomes, to some degree, an island, and being smaller and more isolated it cannot support all the species that it held as part of a larger habitat area (MacArthur & Wilson 1967). As extant species become locally extinct, diversity decreases through time toward a new equilibrium (qua theory of island biogeography; MacArthur & Wilson 1967) between local immigration and extinction. |
| As described by Trepl (1995), both the theory of island biogeography (MacArthur & Wilson 1967) and the metapopulation theory (Levins 1970; Opdam 1991) are presented in studies that try to explain how species abundance and richness are dependent on the habitat patterns in cities. |
| Many butterfly studies conducted at the landscape scale have focussed on the effect of patch size and isolation and have used the theory of island biogeography (MacArthur & Wilson 1967) or the metapopulation theory (Hanski 1999) to explain species richness or population dynamics, respectively (Thomas and Harrison 1992; Baguette et al. 2000; Steffan-Dewenter and Tscharntke 2000; Anthes et al. 2003). |
| For the Krakatau birds, MacArthur & Wilson (1967) concluded that birds approached equilibrium within only 2536 years of the explosion. |
| According to theory of island biogeography (MacArthur & Wilson 1967), the number of species in a defined area is determined by patch isolation (affecting immigrations) and area (affecting extinctions). |
| Since the spruce-fir habitats are often separated by large distances and surrounded by distinctive types of habitat, they can be considered functionally isolated. Accordingly, we hypothesized that carabid beetle diversity might be affected by the same processes that affect islands, as described in MacArthur & Wilson (1967) theory of island biogeography. Namely, we predicted that diversity would increase with area and decrease with isolation. |

Brown (1971) tested MacArthur & Wilson (1967) theory for the small-mammal communities (excluding bats) inhabiting the isolated peaks of the Rocky Mountains in Nevada. Brown found a steeper species-area curve than typical of insular biotas and did not find any relationship between the numbers of species and isolation. He concluded that colonization occurred during the Pleistocene and that the current communities are relicts. He also concluded that the extinction rate has been low and the immigration rate approaches zero (Brown 1971). In consequence, the study system is not at equilibrium (i.e., colonization does not equal extinction) as predicted by MacArthur & Wilson (1967).

MacArthur & Wilson (1967)s theory of island biogeography represents a milestone in understanding distribution patterns of species. Despite various criticisms, additions and revisions to their work, the central idea of island communities being shaped by complex dynamic interactions involving immigration, extinction and evolution is still the basic paradigm of recent island biogeography (Losos & Ricklefs, 2010).

MacArthur & Wilson (1967) applied their equilibrium theory to species numbers. Subsequently, several authors have indicated that processes driving island species richness may also drive changes in the genotypic diversity of their populations (reviewed by Vellend & Orrock, 2010).

theory of island biogeography (MacArthur & Wilson 1967) is often referenced to explain species diversity and use of these habitat fragments.

According to MacArthur & Wilson (1967)s theory of island biogeography (MacArthur & Wilson, 1967), the rate of new species entering a newly formed system by colonization falls as the number of species in the system increases because the chance that an immigrant is already present in the system increases. "The colonization process in the species number of each functional group of the protozoan communities was tested to determine whether the MacArthur and Wilson (1967 MacArthur, R and Wilson, EO. 1967.) model could be fit to it:
$St = Seq(1 - e - GT)$
where St is the number of species at time t; Seq is the estimated equilibrium number of species of ciliate colonization; G is the colonization rate constant; and T90% is the time to reach 90% Seq."

Previous studies have demonstrated that during the primary colonization process, the number of species generally increases and then equilibrates, following the MacArthur & Wilson (1967) equilibrium model equation (Wang et al. 1985; Xu et al. 2005, 2009).

" It should be emphasized that the classic MacArthur & Wilson (1967) model is not simply a saturating curve of accumulated species number; it also presupposes that the number of species is determined by a dynamic balance between immigration and extinction, the rates of which depend on the richness achieved to the moment."

According to rK model (MacArthur & Wilson 1967), an increasing dominance of large and complex individuals is predicted along ecological successions.

Probably the best-known spatial community concept is MacArthur & Wilson (1967)s theory of island biogeography, where the processes of extinction, immigration and emigration lead to an equilibrium number of species in an island community depending on the size of the island and the distance to the mainland.

The conservation of biological diversity in barrier island maritime forests requires the maintenance of constituent small populations. Relative to most mainland forests, maritime forests are limited in size and are also isolated. While fragmented forests may function ecologically as islands (sensu, MacArthur & Wilson 1967), those of actual islands also harbor unlikely species assemblages, composed of species that co-exist either because of vicarious colonization events over time or because they have had continuous occupancy since the Pleistocene (Bellis 1995).

Changes in environmental stress and/or disturbance regimes may have important conservation implications for isolated islands far from the mainland, which often support rare and endemic species and typically have limited opportunities for colonization from mainland sources (MacArthur & Wilson 1967).

As montane forest fragments represent isolated patches we tested whether liana species richness would increase linearly with area on a log-log scale as assumed by the theory of island biogeography (MacArthur & Wilson 1967).

Excerpts of text citing Bazzaz (1979)

| |
|---|
| In general, early successional species are typified by rapid growth, poor shade tolerance, early reproduction, and a short life span (Bazzaz 1979). |
| When light is limiting, differentiation in light capture strategy (Bazzaz 1979) is required for species to coexist (Aikio 2004; Kohyama & Takada 2009). |
| Colonization is critical to the development of vegetation after disturbance (Bazzaz 1979). |
| A number of studies have investigated shifts in species traits during succession until the maximum biomass stage (decades to centuries; in contrast to studies with chronosequences in the order of several thousand years, e.g. Wardle, Walker & Bardgett 2004), some of which compared physiological characteristics of early and late successional species (Bazzaz 1979). |
| Shade-intolerant, early successional species typically have a suite of morphological and ecophysiological traits that support high rates of carbon gain under high light, whereas shade-tolerant, late-successional species are characterized by traits that allow positive carbon gain at much lower light levels, but that limit their ability to maximize carbon gain under high light (Bazzaz 1979). |
| Indeed, structural defences which provide a direct physical barrier for herbivores (Johnson et al. 2010) are higher in mid- late-succession roots (Bazzaz 1979), where plants experience higher levels of herbivory (Figs 1 and 2). |
| The combination of sequestering resources and the potential to reach a taller maximum height can be thought of as late-successional strategy (Bazzaz 1979) a strategy that is successful in the absence of large-scale disturbance. |
| The ecophysiological properties of vascular plant species that comprise the plant community change as the community changes during succession (Bazzaz 1979). |
| Early successional plants are often competitively inferior, and in order to continuously disperse to new open habitats, they need to grow and consume resources quickly. In comparison to species of later successional stages, these early species have high photosynthetic and respiration rates. This faster metabolism is associated with higher stomatal and mesophyll conductance (Bazzaz (1979) and references therein). In many cases, species from the later successional stages have characteristics that are more typical of shade-tolerant plants; they are able to utilize low light intensities more effectively, but tend to have lower light saturation points and lower maximum photosynthetic capacities (e.g., Bazzaz 1979). |
| The subtle differences detected in the preferential micro-habitats for germination and establishment of each species suggested a differentiation in their regeneration niche (Grubb, 1977). These differences may imply that the chance of establishment and coexistence of species varies over time because of the changing environmental conditions within succession (Bazzaz 1979). |
| In closed-canopy forests, competition for light is the primary mechanism driving succession (Bazzaz 1979). Shade-adapted species have physiological and morphological traits that provide for a low whole-plant compensation point and thus low growth rates. |
| Both vegetation development and succession are used specifically to refer to the processes that govern plant growth, reproduction, and mortality across plant species and sizes (Bazzaz 1979). |
| At the community level, this threshold response is accompanied by an ontogenetic niche shift across community species, in which seedlings respond to windows of opportunity in temperature, light and water pulses (Bazzaz 1979). |
| Two hypotheses explicitly address this phenomenon. The first hypothesis suggests body size differences are due to lower food quality in late successional environments (where plants grow more slowly, Bazzaz 1979). |
| Pioneer plants are characterized by high dispersal capacity (Bazzaz 1979). |
| Photosynthetic properties differ between plants that are typical to different successional stages: maximum photosynthetic capacity, dark respiration and light compensation point generally decrease from early- to late-successional species (Bazzaz 1979). |

| |
|---|
| Low wood density and short maximum life span are features typical of early successional species (Bazzaz 1979) |
| The ecophysiology (physiological ecology) of plant succession was initially described by Bazzaz (1979). |
| Following a disturbance (i.e. storm, anthropic impact), the success of primary succession on a foredune (the success of foredune natural community restoration actions) will depend upon the availability of propagules and the potential for establishment (Bazzaz 1979). |
| Bazzaz (1979) reported, in a meta-analysis, that early successional species have high rates of photosynthesis. |
| Negative edge effects can be the result of shading by intact adjacent canopy species (Bazzaz 1979). |
| Hovenia dulcis combines rapid growth at high-light with intermediate survivorship at low-light conditions, traits that are typical of early and mid-successional species (Bazzaz 1979). |
| As succession advances from early to more advanced stages, a community experiences an increase in biomass and density (Bazzaz 1979). |
| Generally, the process of plant succession could accelerate nutrient cycling, and the roots of the pioneer plants could help to increase soil permeability and decrease soil BD (bulk density) (Bazzaz 1979). |
| Pioneer species that regenerate in forest gaps and open areas were hypothesized to have a higher plasticity than shade tolerant species, because they grow in a more variable environment (Bazzaz 1979). |
| In accordance to our results, a number of studies have shown an increase in seed mass during succession (Bazzaz 1979). |
| Drastic environmental changes associated with land-use change may intensify environmental filtering (e.g. forest clearing may increase moisture and heat stresses for ground vegetation; Bazzaz 1979) |
| The removal of established trees increases light availability at the soil surface, which tends to favour colonisation by light demanding species (Bazzaz 1979). Seagrass species possess a wide variety of life history traits and inhabit a continuum of successional capabilities, but can often be categorized as late or early successional species (sensu Bazzaz 1979). |
| Unlike late successional species, early successional (sensu Bazzaz 1979) seagrasses have adapted to disturbance through reliance on rapid disturbance recovery and expansion as opposed to resistance to disturbance itself (Unsworth et al. 2015). |
| In wet forests, decreasing light availability is typically regarded as a major driver of shifts in tree species composition during succession (Bazzaz 1979). |

Excerpts of text citing Huston (1979)

| |
|---|
| Increasing vegetation diversity is expected to increase the diversity of herbivores (i.e. orthopteroids, true bugs, leafhoppers, and butterflies and moths) (Huston 1979). |
| Coexistence between functionally similar species might also result from slower growth rates on nutrient-poor sites, reducing the pace of competition after disturbance (following the dynamic equilibrium theory of Huston (1979) and supported by experimental evidence; Rajaniemi 2003; Wardle & Zackrisson 2005). |
| The maintenance of taxonomic diversity (expressed as either species richness or species diversity) at the oldest sites may depend partly on the size-symmetric nature of nutrient competition, which makes competitive exclusion between species more difficult (Rajaniemi 2003; Wardle<br>Zackrisson 2005; Gundale et al. 2011). This will increase the likelihood of coexistence between species with similar resource-use niches (Huston 1979). |
| The correlations between species diversity, land-use intensity and above-ground standing crop demonstrated in this study support predictions of both the Intermediate Disturbance Hypothesis (Huston 1979) and the biomass-diversity model of Bakker (1989). |
| Such theories predict that high resource availability leads to lower richness because those species with the highest growth rates (Huston 1979) or highest resource acquisition capacities competitively exclude weaker competitors when resource availability is high (Grime 1979, Huston 1979, Grace 1999). |
| Recurring large scale disturbances can prevent competitive exclusion as well, hindering the system to reach the climax state (Huston 1979). |
| Low or reduced levels of disturbance will lead to low diversity through competitive exclusion and the dominance of long-lived species, while high or increased levels of disturbance will eliminate species incapable of rapid re-colonisation and growth (Huston 1979). |
| Traditional views of disturbance, however, assume that frequent disturbance is more or less equivalent to frequent density-independent mortality of all species (Huston 1979). |
| Regarding patterns in the vascular flora, our data showed that maximum species richness is co-incident with the peak of Kalmia cover (90%), which contradicts competition-based causes of plant community structuring such as a centrifugal organisation (Wisheu & Keddy 1992) and the mass-ratio hypothesis of species diversity (Huston 1979). |
| It can be argued that environmental stress is difficult to define and evaluate, given that the inevitable result of natural selection is to minimize the effect of such stress on an organism (Huston 1979). |
| Transient dynamics can occur whereby inferior competitors persist until resource levels equilibrate (Tilman 1988) or disturbance ceases (Huston 1979). |
| "DIVE model (Dynamics and Interactions of Vegetation) uses plant strategy parameters that emerge from functional relationships and climatic constraints, such as growth rate and seed production, that then reflect a populations strategy in being a coloniser or competitor. The performance of a PPS (populations of plant strategies) directly affects these abilities via e.g. the intrinsic growth rate (Huston 1979) or seed production (Angert et al., 2009). |
| The dynamic equilibrium model of Huston (1979) predicts that without disturbance, species diversity will be high on dry sites because of the low growth rates of vegetation (and hence, relatively low rates of competitive displacement). |
| This result is consistent with a combination of the humped-back model (Grime, 1974) and the intermediate disturbance hypothesis (Huston 1979). In these concepts, dominant species competitively exclude other species if dominance is not broken by either stress at low fertility levels or by disturbance. |
| At intermediate disturbance or stress levels, plant species diversity is supposed to be maximised (Huston 1979). |

| |
|---|
| In models formulated primarily to examine the interactive effects of abiotic disturbance and productivity, Huston (1979) (dynamic equilibrim model) and Kondoh (2001) posit that disturbance decreases diversity when productivity is low, is unimodally related to diversity at intermediate levels of productivity, and increases diversity when productivity is high. |
| Nutrient-rich systems are expected to be dominated by few species unless grazers prevent the development of such dominance (Huston 1979). |
| Productivity has been widely cited as a factor affecting ecosystem responses to grazing by large herbivores (Milchunas et al. 1988; Cingolani et al. 2005; Lunt et al. 2007) and to disturbance in general (Huston 1979). |
| According to Huston (1979)s theoretical model, disturbance intensity and competitive exclusion are the fundamental processes controlling species diversity. |
| Among the most influential concepts addressing the productivity-richness relationship are the humped back model (Grime 1979), the dynamic equilibrium model (Huston 1979), and the resource ratio model (Tilman 1982). |
| Our results suggest a strong context-dependent recovery process as biotic interactions changed with shifting environmental conditions, and support the dynamic equilibrium model of Huston (1979). |
| Huston (1979)s Dynamic Equilibrium Hypothesis predicts that the response of biodiversity to disturbance varies with productivity. Because disturbance is thought to break competitive advantage of dominant species in productive ecosystems, species richness is predicted to increase with disturbance frequency in productive systems. Recovery of plant biomass following disturbance is also predicted to be faster in productive systems. |
| The potential for competitive dominance is thought to be greater in productive environments where the onset of competitive exclusion is rapid (Huston 1979). |
| Since bare space did not appear to be not limiting, greater diversity under H. grandifolius suggests that these communities had developed for longer than those under D. menziesii (Huston 1979). |
| Based on classical models in ecology, e.g., Intermediate disturbance Theory (Huston 1979), the Grazing Generalized Model (Milchunas et al., 1988); Stability of Grazing Systems (Noy-Meir, 1975), and the State and Transition Model (Westoby et al., 1989), Noy-Meir attempted to lay a theoretical ground for ecology-based management of open landscapes in Mediterranean Israel. |
| Overall, our results also underline that changes in diversity are expected to be more pronounced on more productive sites because plant growth and hence population dynamics are faster with high productivity (Huston 1979). |
| A strongly competitive pioneer species may restrict local diversity and impact species composition, especially on productive sites (Huston 1979). |
| High productivity, while increasing growth and survival of individuals, also increases competition, potentially leading to competitive exclusion of species (Huston 1979). Accordingly, the DEM predicts that increasing productivity increases species richness at high levels of disturbance, and that increasing disturbance, by increasing mass mortality, decreases species richness at low levels of productivity. When disturbance is low, however, increasing productivity is expected to decrease species richness through extreme competitive exclusion (Huston 1979); |
| The DEM applies not only to disturbance magnitude or frequency but also to time since the most recent disturbance; therefore, species richness is expected to increase with succession time (time since land emergence) at low levels of productivity but decrease it at high levels of productivity (Huston 1979). |
| Competitive exclusion is more likely in stable, uniform environments, while periodic population reductions through moderate disturbance and environmental fluctuations may promote consistently high levels of diversity (Huston 1979). |

Excerpts of text citing Huston & Smith (1987)

| |
|---|
| Diversity and population dynamics are commonly correlated (Huston & Smith 1987). The preponderance of species invasion from the annual to the perennial-grassland stage seems to be not in agreement with Huston & Smith (1987). They mention species replacement as of high importance and more rapidly in early than in late succession. Only if the above presented successional seres are altogether considered as an early successional situation, or at maximum in a change over to mid-succession, than a rough balance between invasion and extinction can be found. However, species extinction and invasion are at no time really balanced. |
| The property of prolonged coexistence in mid-succession is a widely observed feature of natural succession (Huston & Smith 1987). |
| Above all Calamagrostis epigejos, Elymus repens, Solidago canadensis and Urtica dioica are referred to as species which quickly form dominant stands. They may remain stable for many years and are able to inhibit the further development of woody species. These highly competitive species, the so-called superspecies (sensu Huston & Smith 1987). |
| It is commonly assumed, however, that during the course of succession species are replaced by other species that are better adapted (by their traits) to the new abiotic conditions (Huston & Smith 1987). |
| Among-group differences might be influenced by the relationship between dispersal mode and successional status (Huston & Smith 1987). |
| In our study S. canadensis, a non-native perennial species with capacity for vegetative spread, was the species most frequently present in the seed bank (in many plots with very high seed densities). Such highly competitive species, which have been referred to as superspecies (Huston & Smith 1987), often do not fit the general pattern of functional classification. |
| As depicted in models of succession (Connell and Slatyer, 1977), species richness in frequently disturbed areas is not limited by competition for (or tolerance of) limiting resources; instead, it reflects the ability of species to colonise, grow rapidly and withstand local environmental conditions (Huston & Smith 1987). |
| The ready supply of alien plant propagules close to disturbed areas may lead to seed swamping or earlier arrival of alien propagules post-disturbance, enabling alien species to colonise highly disturbed areas more rapidly than native species (see Huston & Smith 1987). |
| It could be expected that lichen traits related to establishment and performance will differ between lichens occurring on young versus old trees, just as life history traits generally differ between early and late successional vascular plant species (Huston & Smith 1987). |
| The positive spore sizetree age relationship can be viewed as support for a general increase in propagule size during succession, as suggested both for lichens (Topham, 1977) and plants (Huston & Smith 1987). |
| To address these shortcomings, mechanistic mathematical models of succession have been developed (e.g., Pastor and Post 1986; Huston & Smith 1987). Their application to forest secondary succession has led to key insights concerning the importance of life-history traits and competition in succession (Huston & Smith 1987). |
| On the other hand, within community similarity increased following the development of a vertical structure in the canopy, which in turn influences the colonization patterns of mid-successional species due to changes in light availability (Huston & Smith 1987). |
| The core theoretical justification behind recent chronosequence approaches, which suggested the transition from clustering to overdispersion was driven by competitive exclusion (e.g., Table S1), is that: (1) competitive exclusion should be more likely to occur among closely related species with similar traits because of the large niche overlap and (2) such patterns will become more prominent at later successional stages because the strength of the species competition would increase as communities mature (Huston & Smith 1987). |
| Correlations between species groups, observed in studies of environmental gradients or succession, therefore, should be traceable to underlying abiotic control factors. However, there is little consensus as to what this underlying control factor might be (Huston & Smith 1987). |

| |
|---|
| In succession, less competitive species are replaced by more competitive ones through time (Huston & Smith 1987). |
| Empirical studies have shown that early successional species, which have relatively high colonization and mortality rates (Huston & Smith 1987), tend to have more negative feedbacks than later successional species (Van der Putten, Van Dijk & Peters 1993; Kardol, Bezemer & Van der Putten 2006; Middleton & Bever 2012). |
| Taken with the fact that feedbacks have been shown to correlate with successional stage (Van der Putten, Van Dijk & Peters 1993; Kardol, Bezemer & Van der Putten 2006; Middleton & Bever 2012), which in turn correlate with colonization and mortality rates (Huston & Smith 1987), it is possible that the observed correlation between feedback and abundance is actually the result of life-history differences. |
| On the nutrient-rich topsoil, successional pathways followed the pattern of sequential succession (Huston & Smith 1987). |
| Stabilizing mechanisms such as competition or ecological/environmental filtering then steadily become more influential during the succession (Huston & Smith 1987) and may lead to lower spatial heterogeneity in taxonomic and functional composition. |
| As far as the late and early species were concerned (here, sugar maple and paper birch), dynamics were simple and fitted perfectly the descriptions of classical succession theory (Huston & Smith 1987). Under a disturbance regime characterized by long return intervals, a stand had time to mature and the late species progressively replaced the early species in the canopy when the latter died. |
| A commonality of these models is that populations of species reach peak abundances at different stages of succession, resulting in easily recognizable species turnover. While light availability is a key driver of these successional dynamics (Huston & Smith 1987), tree populations may also be particularly susceptible to changes in water and nutrient availability. |
| Given the previously documented impacts of water and nutrient availability on stand- and individual-level processes at our study site and elsewhere, we expected controls over forest succession were potentially more complex than overriding autogenic control through light availability (Huston & Smith 1987). |
| Multiple successional pathways ( Huston & Smith 1987) may not only occur due to different habitat conditions but also due to the impact of rare events and the interaction of multiple successional factors. |
| Species that have a high growth rate and low age at first maturity or short generation time are considered well-suited to ecological dominance immediately following a disturbance, and in ecosystems with frequent disturbances (Huston & Smith 1987). |
| Also, the herbivorous fishes maintain large body sizes and moderate life spans. This is counterintuitive to predictions that organisms occupying lower trophic levels would be composed of smaller and short-lived species (Huston & Smith 1987). |
| Comparative investigations that centre solely on plant demography have the potential to expand our understanding of community dynamics and species coexistence (e.g. Huston & Smith 1987) but are an incomplete endeavour if done in isolation. |
| In our simulation, as time progresses and bare area becomes limited, establishment decreases and colonisers are replaced by competitors, consistent with the real world (e.g. Huston & Smith 1987). |
| In large gaps, the increased light availability promotes the persistence of shade-intolerant species allowing them to recruit to the canopy (Huston & Smith 1987). |
| Huston & Smith (1987) argued that a key for understanding natural systems lies in understanding the life history and physiological traits of a single species; thus individual-based models could explain the complex variety of successional dynamics that community-based models usually fail to explain. |
| Every phase of a plant's life history seed production, dispersal, establishment, growth, mortality, and reproduction is influenced both directly and indirectly by climate (Huston & Smith 1987). |

| |
|---|
| These deterministic models are all based on the idea that tradeoffs between traits promote success in different stages of succession (Huston & Smith 1987). |
| Changes in both light availability and soil N (nitrogen) have been documented in many successional sequences (Huston & Smith 1987). |
| Anemochorous, as well as long-distance dispersed zoochorous species, have been identified as quick colonizers of new sites in a wide range of habitats (e.g., Alday et al., 2011; Hardt and Forman, 1989 ; Martnez-Ruiz and Marrs, 2007), although the role of wind dispersal is often replaced by animal dispersal during succession (Huston & Smith 1987). |
| Succession, or the directional, predictable change in biodiversity over time, has captured the interest of ecologists for over a century (Huston & Smith 1987). |
| There may be a continuum of multiple plant growth strategies should transition toward the native grassland state occur (Huston & Smith 1987). |
| These same factors were also used to answer Question 4, by investigating if successional trajectories differ between rock types, e.g. if assemblages develop more quickly or slowly on a certain rock type or if patterns indicative of sequential succession (Huston & Smith 1987) occur on one rocktype but not the other. |
| The traditional view holds that plant succession is largely deterministic, driven by abiotic and biotic filters such as resource availabilities (Drury & Nisbet 1973), facilitation (Connell & Slatyer 1977) and competition (Huston & Smith 1987). |
| The PnET-Succession (simulation) extension also includes physiological attributes such as light saturation intensity, water use efficiency and drought tolerance that allow the realized niche of species to emerge through time (sensu Huston & Smith 1987). |
| Common milkweed (hereafter milkweed) is a fast-growing competitor with a tall canopy, effective clonal spread and seed dispersal ability, broad leaves with high leaf area and effective chemical defence against herbivory (Bhowmik & Bandeen 1976; Agrawal 2004). These traits are characteristic of super species (after Huston & Smith 1987). Succession is a sequential change in relative abundances of the dominant plant species in a community (Huston & Smith 1987). |

Excerpts of text citing Grime (1977)

| |
|---|
| These conditions favour first, fast-growing ruderal vegetation from buried seeds, and ultimately the establishment of competitive species (sensu Grime 1977). |
| Although sheep-faeces deposits can lead to a partial destruction of the covered vegetation and therefore represent a kind of disturbance (sensu Grime 1977), for the purpose of simplicity, only soil disturbance is referred to as disturbance in the following. |
| Disturbance effects on feedbacks may partly act as a mechanism for observed plant community patterns in a manner consistent with Grime (1977)'s CSR theory. More frequent and intense disturbances over time should reduce negative feedback and favour ruderal plant species that are likely to realize rapid growth with decreased pathogen activity, increased nutrient availability and competitive release. With infrequent and mild disturbance, both negative feedbacks and competition are likely to be more important, such that plant communities are dominated by more pathogen-tolerant, competitive species that are typical of late succession. |
| The leaf economics spectrum (LES; Wright et al. 2004; see also Daz et al. 2004) refers to the continuous variation of leaf traits from thin, nitrogen-rich, short-lived leaves with high photosynthetic rates, also referred to as the exploitative strategy (sensu Grime 1977), to thicker, more fibrous, nitrogen-poor, longer-lived leaves with lower photosynthetic rates, also referred to as the conservative strategy. |
| As an ultimate consequence, the inability to cope with environmental stress may lead to tree death. In this context, the individual stress tolerance can be regarded as an important feature of the life-history strategies of trees (Grime 1977). |
| The commonly held view is that species associated with nutrient poor soils are inherently slow growing stress-tolerators (sensu Grime 1977), characterized by low phenotypic plasticity and long-lived organs that maximize acquisition and conservation of nutrients (Grime 1979; Chapin 1980) at a cost to growth (Aerts<br>Van der Peijl 1993). |
| Responses of species associated with intermediate-age sites (P-depleted and relatively poorly drained soils) were more like those of the stress-tolerators envisaged by Grime (1977): low growth rates, high root:shoot ratios and inherent unresponsiveness to environmental heterogeneity (Lambers<br>Poorter 1992). |
| Grime (1977)s classic work on plant life-history strategies maintains that species are constrained by fundamental compromises between the conflicting selection pressures resulting from particular combinations of competition, stress and disturbance, and this inescapably involves the sacrifice of fitness in one scenario for another. A key question in invasion dynamics is whether exotic species are subject to trade-offs that constrain the life histories of native plants, especially tree species. In conditions which support forest growth, life-history trade-offs largely occur between ruderal and competitor life histories (sensu Grime 1977). |
| Pacala et al. (1996) determined that such interspecific trade-offs are necessary for realistic predictions of successional dynamics in native forests, and like Grime (1977), that design constraints prevent the evolution of a hypothetical super species efficient at utilizing the high-to-low light levels found in the forest. If exotic invasive plants are not constrained by this life-history trade-off, this would offer a powerful explanation of the invasiveness of exotic species in forest ecosystems. |
| Competition for resources, for instance, is most important in mid-late succession (Grime 1977), so declines in diversity from exploitative competition are expected to be concentrated there. |
| Fast growth rates are associated with competitive ability (Grime 1977). |
| Stress resulting from insufficient soil resources (water, nutrients) and severe maintenance disturbance may not allow colonization of diverse plant life forms in the shoulder (Grime 1977). |
| Lusk |

Matus (2000) suggested that fast-growing species should dominate on nutrient-rich soils, because these species capitalize on the nutrients available to them by rapid height growth, giving them substantial advantages over neighbours with whom they compete for light. This idea is similar to that expressed in the plant strategy theory of Grime (1977).

An unresolved question is why competition among the fast-growing light-demanding species does not lead to a few species rising to dominance at the expense of others on the nutrient-rich soils, as appears to occur in grassland systems (Grime 1977).

Plants are generally adapted to their natural habitat through different growth strategies. Grime (1977) defines three main strategies: ruderal, competitive and stress-tolerant plants, with the former inhabiting younger and latter older successional stages.

These three groups are to some extent analogous to the three primary strategies suggested by Grime (1977). Pioneer species such as S. fimbriatum are ruderal plants that occur at sites with low stress but high disturbance.

Particularly, vegetative height is usually considered as a good indicator to predict species competitive ability (Grime 1977).

As in other grasslands (Lep and tursa, 1989 ; Gibson and Brown, 1992) after all types of disturbance, Mediterranean steppe is colonized by weedy and ruderal species (sensu Grime 1977) during the first succession stage.

Under natural conditions, secondary succession may be viewed as a gradual transition from ruderal vegetation, in which resources are allocated to the rapid production of seeds (Barbour et al. 1987), to competitive species that maximize vegetative growth or to stress-tolerant vegetation (Grime 1977).

Disturbed soil provides habitat for annual or short-lived perennials, a specialization clearly adapted to exploit environments intermittently favourable for rapid plant growth (Grime 1977).

Zimdahl et al. (1988) showed experimentally that paddy weeds germinated immediately after soil tillage. Owing to the ruderal strategy defined by Grime (1977), summer annual TPWs can complete their life cycle before rice covers the soil.

Successful establishment occurs where species life-history strategies are compatible with the prevailing establishment or regeneration conditions generated by the disturbance regime (Grime 1977).

s. The absence of an isolation effect for annual species is not so surprising, because these species are known produce a large number of highly dispersive, dormant seeds (Grime 1977).

Grime (1977)s life strategy classification describes the trade-off that organisms face when the resources they gain from the environment are allocated between growth, maintenance or regenerationknown as the universal three-way trade-off (Grime 1977). Plants utilizing the Competitor strategy facilitate their survival by using methods that maximize resource acquisition and resource control in consistently productive niches. Those plants using a stress-tolerant strategy maintain metabolic performance in variable and unproductive niches. Species with a Ruderal strategy accomplish rapid completion of their lifecycle and regenerate in niches where events are frequently lethal to the intermediate strategies.

Competition results when plants share limited resources. Inter-specific competition is clearly influenced by geoclimatic conditions and plants can respond using two different strategies. Firstly, plants may choose faster growth to occupy more space and, as a result, improve their chances of obtaining new resources (Grime 1977).

Disturbances can be defined as temporally discrete events which abruptly kill or displace individuals, or that directly result in a loss of biomass from a system (Grime 1977).

Disturbances that resulted in a loss in biomass from a community (sensu Grime 1977) affected the subtidal macrobenthic community on a temporal scale.

| |
|---|
| Plants are most likely to form associations with and benefit from mycorrhizal fungi under conditions in which availability of one or more soil nutrients, including water, is low (Hoeksema and Schwartz 2003; Jones and Smith 2004). There are at least two primary situations in which this might occur. One is in environments with a low frequency of disturbance and high abiotic stress, where soil nutrients are not sufficient to support high aboveground growth rates and carbon (C) allocation to belowground biomass is usually proportionately greater (Grime 1977). |
| The availability of nutrients could influence plant growth and colonization in two possible ways. First, if nutrient availability is low, plants produce less biomass and hence would colonize over smaller distances. Second, plants in short supply of nutrients may start to explore for better sites and would colonize over relatively larger distances per unit of biomass when compared to nutrient-rich situations (Grime 1977). |
| Eutrophic species put more biomass into shoot production compared to mesotrophic species, eventually resulting in the exclusion of mesotrophic species due their better competitive ability (Grime 1977). |
| First, given the more or less parabolic relationship between nutrient richness and biodiversity, strong eutrophication often results in vegetation with a lower biodiversity (Grime 1977). |
| The relative ability of species to perform (species performance) in different environmental conditions also influences successional dynamics. Resource availability and the ability of populations to capture those resources (Tilman 1986), ecophysiological plant traits (Larcher 1995), stress and species' ability to avoid or tolerate stress (Grime 1977), and trade-offs associated with life-history strategies (Crawley 1997) influence the success or failure of a species. |
| Indeed, competitive ability depends upon plant characteristics that maximize vegetative growth in productive, relatively undisturbed conditions (Grime 1977: 1183). |
| As the three primary strategies are related to the levels of productivity and disturbance at a given site, their change in space and time may serve as a measure of key processes such as succession, eutrophication and other changes in habitat conditions (Grime 1977). |
| The three-strategy (CSR) model proposed by Grime (1977) constitutes one of the most established systems for plant functional types. The primary strategies (competitive ability, adaptation to severe stress and adaptation to disturbance) relate to the productivity and level of disturbance at a given site. |
| Facilitation by micro-site improvement is clear in the cases of A.millefolium and S. jacobea; two species that require a relatively high soil nutrient status (Grime 1977). However, escape from herbivory might be the reason for the increase in Lactuca spp. and H. radicata, two ruderal species with large leaves and rapid growth (sensu Grime 1977), which showed greater cover values under greater shrub volumes, reducing the grazing damage underneath by physical shelter (Baraza et al., 2006). |
| Photosynthetic properties differ between plants that are typical to different successional stages: maximum photosynthetic capacity, dark respiration and light compensation point generally decrease from early- to late-successional species (Bazzaz 1979), while the physiological stress experienced by the plants increases (Grime 1977). |
| To examine photosynthetic strategies of the moss species a posteriori, we classified the species in three categories after Grime (1977): ruderal, competitive, and stress-tolerant, based on their PPFDc and PMAX. |
| The moss species can be classified in the three groups defined by Grime (1977) as (i) ruderal species that show high production and occupy recently disturbed areas, (ii) competitive species that show high production and occur in more stable conditions, and (iii) stress-tolerant species that show lower production but are more adapted to stress or resource scarcity (Table 5). |
| Both undrained and drained spruce swamp forests can be compared with the late-successional stage of forested vascular plant communities, where succession is associated with decreased availability of resources (Grime 1977). |

| |
|---|
| The three species strategies, as defined by Grime (1977), can be placed along the successional gradient: stress-tolerant P. schreberi, S. magellanicum, S. russowii, and S. angustifolium at the late-successional stages, ruderal S. riparium occupying recently disturbed areas and competitive S. girgensohnii during mid-succession. |
| Though the dynamic equilibrium model defines disturbance as any factor that reduces population size (abiotic disturbance, grazing, predation intensity), there is some debate in the literature about whether grazing predation should (e.g. Grime 1977) or should not (McGuinness 1987) be considered a disturbance. |
| In the model, competition for one or more resources exponentially increases in intensity with decreasing environmental severity, potentially reaching a plateau in very low-severity environments. This is supported by both empirical (Goldberg et al. 1999) and other theoretical models (Grime 1977). |
| Physiological and demographic responsiveness to environmental severity also differs dramatically across species (Grime 1977). |
| Mature plant traits are often uncorrelated with seedlings traits (Grime 1977), it is necessary to measure traits at multiple ontogenetic stages if both effects and responses are of interest. |
| According to the Grime (1977)s triangular model for representing plant strategies (CSR theory) (Grime 1977) plant competition is not important in recently disturbed areas with great availability of resources and rapid colonization. |
| While P. tomentosa possesses many characteristics of a classic ruderal species (high fecundity, early reproduction, rapid growth; Grime 1977) its ability to tolerate water stress appears to be critical to its ultimate persistence. |
| Differences among plant species in their ability to compete with other species and utilize resources can dictate community composition and successional patterns after disturbance events (Grime 1977). |
| Reduction of metabolic rate is a widespread mechanism to improve resistance to stress factors, and a low rate of growth and development is a common feature of stress tolerant plant species (Grime 1977). |
| A high rate of growth and development is regarded as a basic feature of ruderal plants adapted to environments with periodically deteriorating conditions (Grime 1977), since rapid seed production ensures restoration of populations after disturbances. The extremely slow diameter and shoot growth of seedlings in undisturbed forests (Figures 6, 7) is indicative of a species with a stress-tolerant strategy (sensu Grime 1977). In general, ruderal species are considered to be more homogeneous in their functional traits compared to competitive species (Grime 1977). |

| Excerpts of text citing Hubbell (2001) |
|---|
| Recent community-level studies analyzing functional traits and species abundances have shown that niche-related processes play a role in community assembly even in tropical forests (Gunatilleke et al. 2006, Engelbrecht et al. 2007, Kraft et al. 2008), where unpredictable chance events are proposed to play a leading role (Hubbell 2001). |
| Community assembly is strongly influenced by propagule dispersal, which produces heterogeneous assemblies by random dispersal (Hubbell 2001). |
| Neutral theory, in contrast, predicts that community composition is determined solely by stochastic events, historical contingencies, and random dispersal events (Hubbell 2001). |
| By examining early life stages in second-growth forests, we adopt a novel approach to investigate the relative importance of niche and neutral processes in determining community reassembly in secondary strands in the absence of long-term data. Early life stages provide a unique opportunity to address this issue because young individuals better reflect the footprint of dispersal events, stochastic demography (Hubbell 2001). |
| Random phylogenetic structure is predicted by Neutral theory (Hubbell 2001). |
| Straight-line distances between plots were ln-transformed, as suggested by Condit et al. (2002) and Jones et al. (2006), and postulated by Hubbell (2001)s Neutral theory. |
| This result extends the general tendency of decelerating rates of community change with time, as described by Anderson (2007), based on presence/absence data, to data sets based on abundance. Our findings are also consistent with Hubbell (2001)s Neutral theory. |
| In effect, these rare, but ecologically important events may decrease the influence of trait-based processes relative to the importance of trait-neutral stochastic processes. In his provocative Neutral theory, Hubbell (2001) even claimed that traits do not matter at all. The probability of colonizing a site would then merely become a lottery in which chances are weighted by the availability of seed sources in the surroundings. |
| Neutral theory (Hubbell 2001) assumes that species are competitive equivalents with the same prospects for survival and reproduction. In this situation, functional differences between species are unimportant and communities are the result of processes such as dispersal, speciation, and extinction with ecological processes such as competition playing a limited role (Hubbell 2001). |
| Null models were used to compare these community phylogenetic distance metrics to expectations of Neutral theory (Hubbell 2001). |
| Stochastic (trait-neutral) processes (Hubbell 2001) are expected to result in species co-occurrence patterns that are random with respect to species traits. |
| Species presence in a community can be caused by a combination of stochastic dispersal and establishment processes (Hubbell 2001) combined with subsequent species interactions such as competition or facilitation (Callaway & Walker 1997), in which case differences in the pool of functional traits among successful species would be emphasized (Valladares et al. 2008). |
| Biodiversity is the result of neutral drift, in which all species of the same trophic level have equal chances of becoming abundant and are functionally equivalent, despite having different functional traits (Hubbell 2001). |
| Stochastic processes have been explored with null models (Hubbell 2001), distance (Bossuyt et al. 2003), contingencies (Bakker & Moore 2007) and climate variation (MacDougall et al. 2008). |
| This empirical study provides a useful test of the current theoretical debate on the underpinnings of community assembly, as it supports the view that dispersal assembly (Hubbell 2001) and niche assembly (Chase & Leibold 2003) are non-exclusive mechanisms that shape the composition of a plant community. |
| If community dynamics are slow, assembly history can have long-lasting effects on plant community composition (Hubbell 2001). However, sooner or later late species will get new opportunities to colonize areas via disturbance or the death of currently established species. |

| |
|---|
| This result contrasts with a neutral expectation that any observed spatial turnover in species composition should occur independent of differences in competitive traits or demographic rates (Hubbell 2001). |
| Dispersal limitation has also been invoked as a major mechanism in neutral models in which community structure is the outcome of stochastic drift in the abundance of species with equal fitness (Hubbell 2001). |
| These successional patterns are in some ways reminiscent of those observed in diverse tropical tree communities, where a small number of species have distinctly ruderal life history strategies but the vast majority of dominant species target similar niches (Hubbell 2001). |
| Seed dispersal and recruitment limitation are integral to plant community development. In combination they shape the context in which populations succeed or fail (Schupp et al. 1989; Howe and Miriti 2004; Norden et al. 2009b), determine success of emigration and immigration (Nathan and Muller-Landau 2000; Nathan 2006), drive ecological succession (Connell and Slatyer 1977; Ronce et al. 2005), determine templates of plant community composition (Schupp 1995; Levine and Murrell 2003), and set boundaries within which stochastic processes shape species-abundance distributions (Hubbell 2001). |
| The importance of this broad category to constraining succession is highlighted by the concept of initial floristics where succession is determined by initial colonists (Egler, 1954) and is critical to Neutral theory, which focuses on stochastic dispersal and colonization events (Hubbell 2001). |
| Large sets of species are a considerable problem for any study. Thus, most multi-species models are extremely laborious to parameterize (e.g. Kimmins et al., 1999) or strongly simplified (Hubbell 2001). |
| Therefore recruitment or seed input should be a function of the surrounding vegetation. However, if the number of seeds or seedlings is correlated with the abundance of the surrounding vegetation then mechanisms which maintain the coexistence of the different species must be considered. Otherwise the system will end in a monodominant state (as shown in Hubbell 2001). |
| As a suite of dispersal and establishment bottlenecks increases the proportion of ecological equivalent species in post-agricultural forest (cf. Verheyen et al. 2003; Vellend et al. 2007; Baeten et al. 2009b), demographical stochasticity may have partly replaced interspecific interactions in structuring local abundances (Hubbell 2001). |
| In contrast, stochastic (trait-neutral) processes (Hubbell 2001) are expected to result in species co-occurrence patterns that are random with respect to species traits. |
| The results have implications for the question whether and to what degree neutral mechanisms (Hubbell 2001) affect community assembly. |
| Competitive similarity, which acts as an equalizing force (sensu Chesson 2000), makes the outcome of interspecific competition sensitive to arrival order owing to neutral drift (Hubbell 2001). |
| Lack of support for the heterogeneitydiversity relationship in some studies has been explained in a number of ways, including dispersal limitation (Questad and Foster 2008), predominance of stochastic processes (Hubbell 2001), and overriding resource limitation (i.e. diversity is more strongly related to mean resource availability than variation in resources; Lundholm 2009). |
| These findings contrast with the perception that interdependent processes among plant species are insignificant over evolutionary time frames, which is an idea underlying both the concept of communities as a mere coincidence of constituent species (Gleason 1926) as well as the development of Neutral theory on biodiversity (Hubbell 2001). |
| Species growth and mortality rates may not covary if tree death is entirely stochastic and unrelated to species traits (Hubbell 2001). |
| A limiting case of a metacommunity is a mainland-island configuration the familiar domain of island biogeography. In this case, there is pronounced asymmetry in dispersal, and a clearly defined external source pool from which colonists are drawn. More broadly, all habitats in a region can in principle provide dispersing propagules that can appear in any other habitat. Metacommunity ecology is thus a generalization not just of metapopulations from one to many species, but of island biogeography (Hubbell 2001). |

| |
|---|
| This means that the lifetime reproductive success (per generation population increase) of either species, introduced at low density to the other, is neither negative nor positive, but exactly zero. This is the characteristic of neutrality, which leads to ecological drift (Hubbell 2001). |
| Indeed, many of our most fundamental ecological models and concepts, such as island biogeography (MacArthur and Wilson 1967), metacommunity dynamics (Leibold et al. 2004), succession (Clements 1916), neutral models (Hubbell 2001), and community stability (Ives and Carpenter 2007) are based on temporally dynamic phenomena, yet they are frequently tested with spatial rather than temporal data. |
| Caused in part by interest in the Neutral theory of community structure and biodiversity (Hubbell 2001), rank-abundance curves and curve-fitting procedures have experienced a resurgence (McGill et al. 2006). |
| It is important to note that any connection between traits and abundance does not preclude an effect of demographic stochasticity on abundance (see McGill et al. 2007). However, ecological equivalence and drift (sensu Hubbell 2001) do not predict any traitabundance connection. |
| Equal chance is a component (but only one) of Hubbell (2001)l's Neutral theory, and almost all tests of the latter have been weak: failure to find departure from null-model predictions. |
| Increases in -diversity can also emerge in fragmented rain forest landscapes, as fragmentation limits interpatch dispersal patterns, and can therefore promote the compositional differentiation between forest patches and landscapes (Hubbell 2001). |
| Neutral community assembly (Hubbell 2001) assumes ecological equivalence of species (identical trait space) within a given ecological guild and ecological drift as the major process of colonization. |
| This type of random sampling complies with predictions from neutral models (Hubbell 2001), we can conclude that species in this subtropical forest community are not assembled randomly. |
| Neutral theory assumes that all species are functionally equivalent and species assemblages are formed by stochastic drift (Hubbell 2001). |
| Nearly all ecological communities are assembled from a small number of common species and a large number of rare species (see Hubbell 2001). |
| Along with competition and the abiotic environment, species availability is another important determinant of plant community composition (Hubbell 2001). Along with presence in the community as reproducing adults, dispersal from outside the focal community is a primary determinant of species availability. |
| According to the Neutral theory, coexistence of many species is possible when species are similar in their competitive abilities and dispersal rates (Hubbell 2001). An extension of Neutral theory involves the selforganization of species into clusters where members of a cluster are similar in their competitive abilities but the average competitive abilities of clusters are sufficiently different to allow coexistence of many clusters (Scheffer and van Nes 2006). |
| It was likely that these assemblages were supersaturated because the coexisting species were very similar in their competitive abilities, that is, neutral coexistence (Hubbell 2001). |
| Colonization success may be dependent on stochastic variation during immigration resulting in a random selection from the species pool (Hubbell 2001). |
| Their relative abundances would be determined by stochastic events during colonization, rather than ecological advantages determined by their phenotypes. This model of community assembly is assumed in the Neutral theory (Hubbell 2001), which has successfully predicted species distribution curves for communities of macroscopic organisms (Hubbell, 2005) as well as microbial communities (Sloan et al., 2006). |
| The neutral models popularized by Hubbell (2001) additionally assume ecological equivalence of species (identical trait space) within a given ecological guild. |

| |
|---|
| Here, metacommunity dynamics refers to species immigration rates from the panmictic species pool (metacommunity) to multiple sites, the multispecies interactions that govern recruitment and extinction rates at local scales, and the properties of the neutral (logseries-like) metacommunity determined by speciation-extinction at large spatial scales (Hubbell 2001). |
| Therefore, to keep the excess of rare species in the metacommunity at speciation-extinction equilibrium, species must preferentially originate at small population sizes (Hubbell 2001). |
| To cite Hubbell (2001) (p. 90): the expected abundance of the ith species at equilibrium in the local community is simply equal to the local community size, J, times the metacommunity relative abundance of the ith species. Once immigration into the local community has occurred, then, since the individual probabilities of survival and reproduction in the local community are assumed equal, the structure of the local community is determined entirely (except for random sampling fluctuations) by the relative abundance of each species in the larger landscape. |
| First, the majority of species in our data base are relatively rare, and rare species are more likely to be lost with fertilization (Suding et al. 2005) and owing to random demographic stochasticity exacerbated by environmental fluctuations (Hubbell 2001). |
| Species will show spatial auto-correlation even in homogeneous environments due to biological processes, such as dispersal limitation (Hubbell 2001). |

Excerpts of text citing Pickett et al (1987)

| |
|---|
| Succession is affected by multiple interacting factors, both intrinsic (autogenic) and extrinsic (allogenic) to plant communities (Pickett et al 1987). |
| Local disturbances has been stressed as important sources of successional change, controlling inhibition mechanisms (Pickett et al 1987). |
| One promising avenue towards resolution of species composition puzzles is to more closely examine the availability of propagules (Pickett et al 1987). |
| The purpose of the original conceptual framework was to organize successional thought in a useful and logical manner (Pickett et al 1987). While there is no specific temporal or spatial scale associated with the original hierarchical view, it can easily be rearranged into a filter model to describe successional dynamics within a single site. In this usage, each level constrains lower levels, but there is also the potential for feedback among levels.Simply, site conditions filter out species able to survive under local conditions from those that reach the site and those are then sorted out via competition or other determinants of performance. |
| La primera fase de la sucesin secundaria, luego de una perturbacin, es la invasin o regeneracin de especies pioneras y el establecimiento de las plantas en el sitio perturbado (Pickett et al 1987). |
| Los procesos sucesionales son esencialmente demogrficos y producen cambios en la estructura de la comunidad y funciones ecosistmicas (Pickett et al 1987). Si las especies colonizadoras no permiten el establecimiento de especies sucesionales tardas por inhibicin, la sucesin quedar detenida o retardada en sus etapas iniciales. |
| Estos tres modelos (tolerancia, facilitacin e inhibicin) pueden actuar durante la sucesin como hiptesis no excluyentes (Pickett et al 1987). |
| En consecuencia, Sphagnum podra actuar como especie facilitadora para el establecimiento de especies como E. coccineum en el matorral sucesional, aunque en el mismo sitio podra inhibir el establecimiento de otras especies. As, los mecanismos de facilitacin e inhibicin actuaran simultneamente en una misma etapa de la sucesin (Pickett et al 1987). |
| Disturbances that damage the local soil surface force the succession processes to recommence from the level of the disturbance (Pickett et al 1987). |
| First outlined by Pickett et al (1987), successional management is based on the three primary causes of succession: site availability, species availability, and species performance. |
| When plant communities are dominated by one functional group, restoring them can be difficult due to reduced species performance (Pickett et al 1987) associated with competitive interactions between seeded and existing vegetation. |
| Following the framework of Pickett et al (1987), competition for growth-limiting resources is conditional upon site and species availability. That is, propagules must be both present within the landscape and capable of dispersing into and establishing in new sites. |
| Mechanisms affecting post disturbance succession are complex, but an important factor is species performance after arrival in the site (Pickett et al 1987). |
| As part of an integrated weed management and restoration program, it may be desirable to manage bird-dispersed weeds and native species simultaneously, through manipulation of species availability, site availability or species performance (sensu Pickett et al 1987). |
| Establishment can be manipulated by altering either site availability or species performance (sensu Pickett et al 1987). |
| Many mechanisms can contribute to temporal turnover in community composition, including plantsoil feedbacks (Kardol et al. 2006), facilitation, inhibition, herbivory, life history traits, and competitioncolonization trade-offs (Pickett et al 1987). |
| The chronosequence approach has a few well-recognized limitations that include the level of detail in which changes can be deduced (Pickett et al 1987). |

| |
|---|
| Applying these reviews to direct succession of conifers on mineral surfaces suggests several mechanisms that may be especially important in controlling conifer colonization of the rock-fall surface. These, listed in their possible order of declining importance, include: (1) active tolerance (sensu Pickett et al 1987) of the physical and chemical environment of the coarse mineral surface; (2) potential facilitation of establishment and growth by nurse plants or nitrogen fixers (Walker and del Moral 2003); (3) tolerance (both active and passive sensu Pickett et al 1987). |
| Pickett et al (1987) provided the theoretical basis for successional management by developing a hierarchical model that includes the general causes of succession, controlling ecological processes, and their modifying factors. The three causes of succession, including site availability, relative species availability, and relative species performance must all be considered in developing an integrated land-management program. |
| In fragments occupied by nonsuccessional pioneer species, disturbances of sufficient intensity, duration, and frequency may relieve competitive inhibition and enhance succession (Pickett et al 1987). |
| On substrates with an existing matrix of bacteria, prostrate and short stalked diatoms, facilitation (Pickett et al 1987) might increase D. geminatas chance to attach (Stevenson 1983). |
| The concept of vegetation dynamics was proposed to account for successional changes on a site at a single species or individual level (Pickett et al 1987). |
| The analysis of aerial photos indicated that about 88.5% of the surface occupied by the dwarf community was replaced by juniper scrub over a 50-year period, and the latter currently shows scarcity of open microhabitats. This finding, together with the information about land-use change, is evidence of a process of secondary succession (Pickett et al 1987). |
| Earlier stages were marked by coexistence of species with different growth forms and life history strategies, consistent with the passive tolerance model [Pickett et al (1987)].The successional mechanism switched to active tolerance in later stages when later immigrants, having better access to resources, suppressed the growth rates of early colonizers in both low and high current regimes. |
| The role of pioneer species favouring or precluding the regeneration of late-successional species is largely documented in the literature (Pickett et al 1987). |
| Pickett et al (1987) identified three general causes of succession: species availability, site availability, and species performance. If recruitment is to occur, propagules of the species must be present, safe sites must be available to the propagules for establishment, and the species must perform well enough to survive over time. |
| Effects of rock type may interact with the stage of assemblage development, indicating a divergence in successional trajectories between rock types (Pickett et al 1987). If this were true, then changes over time would be expected in assemblage structure and individual species abundances on new substrata, and the changes should be consistent among random locations but dissimilar on boulders of different rock type within and/or between different reef types. |
| Currently we know very little about the consequences of such management history on secondary succession as it relates to tree and understory plant diversity. Will diverse hardwood forests come back naturally through succession as those seen in old-field succession in North America (Pickett et al 1987)? |
| Vegetation dynamics during succession were viewed as emerging from properties of component species (Gleason 1926), rather than from a holistic, organismal concept of community development (Pickett et al 1987). |

| |
|---|
| There are multiple potential trajectories for succession, including single or multiple pathways that can be parallel, convergent or divergent, but that also can be cyclic or form complex networks (Fig. 1; Walker<br>del Moral 2003). Single and cyclic pathways are the most easily adapted to space-for-time inferences because they typically have few dominant species and few stages (Watt 1947). |
| The degeneration of dominant plant stands consequently results in patches that are open for recolonisation. As local cycles tend to be out of phase, maturation and degeneration will lead to a mosaic of community states that shift across the habitat. Some well-known examples also describe these shifting mosaics in various ecosystems, for instance grasslands (Watt 1947). |
| In some of these examples the shifting mosaics are driven internally, i.e. by biotic interactions (Watt 1947) whereas others are driven by external disturbances, such as flooding (Kollmann et al., 1999). |
| The lack of a very obvious and significant response in NPP (net primary productivity) to warming at the colder sites in our experiment may be related to the age of the C. vulgaris plants, which are in a mature phase and therefore not accumulating much new biomass (Watt 1947). |
| The rejuvenating process of C. vulgaris (Watt 1947) will determine which species will prevail in the habitat, which again depends on a number of natural disturbances and conservation/management initiatives (Bruggink 1993; Diemont and Linthorst Homan 1989). |
| Disturbance regimes can often induce successional changes (Watt 1947), which lead to waves of colonization potential and extinction risks, effectively shifting the spatial configuration of patches or successional states. |
| A. G. Tansley stated, I have always tried to impress on my students, perhaps sometimes ad nauseam, the essential importance of the study of succession (Tansley 1939). Similarly, a major component of A. S. Watts research explored temporal changes in communities (Watt 1947). Both Watt and Tansley also appreciated that spatial dynamics (e.g. propagule availability) could affect the direction of succession. |
| A major goal of ecology is to link the patterns observed in populations, communities, and ecosystems to the processes that generate them (Watt 1947). |
| The increase-when-rare mechanism here is similar to that of circular interference networks, but the cycle is between vegetation phases, not individual species. Moreover, it can operate with only two phases. Patches are involved, but their environmental differences are autogenic, not allogenic. Watt (1947) excellently reviewed the subject, but unfortunately none of his examples have been supported by subsequent work. |
| It has been thought that more diverse ecosystems are more likely to resist invasion (Yachi & Loreau 1999), and it could be that C. vulgaris and E. nigrum growing together do better at resisting grass and tree invasion, as these species might complement each other's senescent phases in a similar way as was described for C. vulgaris and Arctostaphylos uva-ursi growing together (Watt 1947). |
| The patch dynamics mechanism was first elaborated and applied to several different plant communities in a seminal paper by Watt (1947). A patch-dynamic system consists of a spatial mosaic of patches in which the same cyclical succession of patch states occurs (Fig. 1). |
| Apart from theoretical approaches, patch dynamics have also been shown to be an appropriate description for several different communities, including heathland (Watt 1947), rocky intertidal communities (Levin & Paine 1974), forests (Remmert 1991), grasslands (Coffin & Lauenroth 1990), agricultural communities (Kleyer et al. 2007) and plankton communities (Steele 1978). |
| Since Watt (1947)s first account of patch dynamic ecological systems, patch dynamic modelling frameworks have been developed (Wu<br>Levin 1997; Levin et al. 2001). |
| Most vegetation is in the mature and degenerate phases of the dwarf shrub cycle (Watt 1947) but there is also short vegetation of mown rebreaks. |

| |
|---|
| Previous work suggested a model of cyclical succession (sensu Watt 1947) where the interaction between shrubs and grasses drives the cycle and determines the transitions among successional stages (phases) that coincide with specific structural patches (Soriano et al., 1994, see Aguiar and Sala (1999) for a diagram of the model). |
| A landscape is an assemblage of ecological communities in which the aggregates of similar individuals and species compose a patch; patches of various types in combination create a juxtaposition landscape, the mosaic (Watt 1947). |
| Forest gap models established according to the approach of Botkin et al. (1972) and Shugart West (1977) are based upon the concept of gap phase replacement (Watt 1947). |
| More than 60 years ago, Watt (1947) provided the first comprehensive description of clonal plants forming rings of regular shape. Herbs, shrubs and even trees, during their ontogenetic cycles, can produce clones of circular shape that progressively degenerate in the older inner area, thus producing a weaker or dieback central zone (Fig. 1). |
| However, central dieback has been also reported for some woody shrubs and trees (Electronic Appendix: Table 1) capable of clonal growth from prostrated stems and branches (e.g., Calluna vulgaris in heathland, Watt 1947). |
| "Watt (1947) firstly suggested that environmental abiotic stress may play an important role in the formation of rings, identifying drought as the primary causal factor for central dieback in Agrostis patches in Breckland grassland ecosystem. In particular, Watt (1947) observed that severe summer droughts selectively killed the inner, weaker clump of the perennial grasses Agrostis tenuis and A. canina. " |
| "In his citation classic paper, Watt (1947) firstly reported that the ramets in the inner areas of clones of the perennial grasses Agrostis tenuis and A. canina were primarily and selectively affected, up to senescence, by persistent water shortage after severe summer drought, while those at the periphery were always surviving. " |
| The processes that shape the dynamics and thus the patterns of plant communities have fascinated ecologists for a long time (e.g. Watt 1947). |
| Watt (1947) developed a theory of pattern and process in the plant community, that was influenced by ideas of Gleason (1927, 1939), emphasizing the importance of individuals and species in a cyclic succession process. Both Watt (1947) and Gleason recognized the importance of processes such as population dynamics, competition and tolerance for understanding the mosaic of vegetation patches resulting from forest succession. Since these seminal works, predicting the direction of changes in species composition and stand attributes over time has been a major focus of forest ecological research. |
| Watt (1947) recognized that natural disturbances are an integral component of the development cycle, which may lead to large canopy gaps, providing a distinct microclimate and thus influencing the competitive dynamics in the plant community. |
| Pattern and process are tightly interwoven (Watt 1947). |
| Because disturbance maintained the presence of lesser bryophyte competitors through the constant presence of intermediate succession (Kimmerer and Allen 1982), moss harvesters could be said to serve as third party facilitators (after Glenn-Lewin and van der Maarel 1992) in an artificially accelerated pattern of cyclic development (Watt 1947). |
| Watt (1947) developed the concept of cycle of change in forests or forest cycle (Whitmore 1989) which corresponds to a cyclic patch dynamics, taking place anywhere in the forest in an asynchronous way. |
| " According to Watt (1947), the cycle may be divided into two series: an upgrade series, characterized by a continuous change in age, growth rate and density of dominant tree species and by an increase in primary production and vegetation biomass; a downgrade series associated to an increase in dead and dying trees, the occurrence of typical gap species and the decrease in productivity. " |

| |
|---|
| "These vestiges are a proof of the dynamic relation between species and mosaic patches. A community subjected to this type of dynamics consists of patches differentiated by structure, floristic composition, age of dominant species and habitat (Watt 1947)." |
| Thus, there are in fact two cyclic processes among the populations of the dominant tree species. This fact does not facilitate the identification of the upgrade series and downgrade series (sensu Watt 1947). |
| Most landscapes are characterized by disturbances that create spatial heterogeneity in community structure. Local disturbances generate a mosaic of communities driven by ecological processes of succession (Watt 1947). |
| Our study contributes to understanding the dynamics of plant communities in subtropical coastal dunes through the patternprocess approach, which has proven to be an essential method for understanding community organization (Watt 1947). |
| This stage-dependent pattern of facilitation and competition has the potential to generate a cyclical succession (sensu Watt 1947) of ant species on acacias as host plants grow older. |
| A general principle of forest dynamics proposes that the most shade-tolerant tree species are competitive dominants in late-successional forests, whereas pioneer tree species persist by rapid establishment following disturbance (Watt 1947). |
| Both biotic and abiotic factors influence long-term regeneration dynamics (Watt 1947). First, biotic interactions, such as dispersal mutualisms and facilitation among nearby individuals, strongly shape the initial recruitment stages (Seidler<br>Plotkin 2006; Hampe et al. 2008). |
| Based on the theory of patch dynamics (Watt 1947) tree development (growth), establishment, and mortality are simulated with an annual time step on small areas (patches), while the influence of climate and ecological processes is taken into consideration using a minimum of ecological assumptions. |

Excerpts of text citing Tilman (1987)

| |
|---|
| Species typical for pioneer stages are significantly more rapidly reduced by nitrogen enrichment, while mid-successional species are fostered. Transient cover increases of certain species have been reported from other plant communities (Tilman 1987) |
| Accumulation of nitrogen in the ecosystem is probably one of the main driving variables that determine the rate of succession (Tilman 1987). |
| To assure that N was the only limiting nutrient (Tilman 1987), we also added the same amount of P (10 g P2O5 m2 yr1), S (0.2 mg m2 yr1), and trace elements (Zn: 190 g m2 yr1, Mn: 160 g m2 yr1, B: 31 g m2 yr1) for all treatments with the exception of control based on the soil census data (Inner Mongolia Soil Census Office & Inner Mongolia Soil and Fertilizer Service, 1994). |
| The rapid dominance of annuals following N (nitrogen) addition in other previously degraded sites was short-lived (¡2 years, Tilman 1987) likely because the perennial grasses were more dominant in the long run on these dry-sandy soils as opposed to the more moist soils of the mature grassland of Inner Mongolia. |
| In this case, as with many novel grasslands across the United States, the sites had a legacy of nutrient enrichment that has been shown experimentally to lead to diversity declines (Tilman 1987). |
| The rank clocks from Cedar Creek clearly show the decrease in species richness in response to fertilization relative to the control (Tilman 1987). |
| Both measures provided unique information on the rapid changes in composition and rank abundance in the N fertilization experiment at Cedar Creek (Tilman 1987) |
| Nitrogen is an essential macronutrient needed in large amounts by plants for survival (pik and Rolfe 2005) and low levels of available nitrogen can limit plant growth and species diversity (Tilman 1987). |
| Nutrient limitation is well known to shape plant community structure (Tilman 1987). |
| Quantifying plantsoil feedbacks relevant to the dynamics of invasions in nature requires field experiments incorporating natural climate variation and competitive regimes, both of which strongly influence how plants respond to soil nutrients (Tilman 1987). |
| Competition also structures vegetation during succession, by contributing to species substitution: late successional species displace early successional species by competition (Tilman 1987). |
| Tilman (1987) suggested that prairies may be nitrogen-limited in wet years and water-limited in drought years. |
| Alternatively, Lespedeza, especially at greater abundance, can lower light availability limiting the abundance and distribution of shade-intolerant species, such as other N-fixing species (Tilman 1987). |
| However, the main driving variables for change in species composition may be the lower availability of light caused by a closing stand canopy (Tilman 1985, 1988, Poschlod et al. 1998, Stampf li and Z eiter 1999) and accumulation of nitrogen in the ecosystem (Tilman 1987) |
| Meta-analyses across herbaceous ecosystems find that N enrichment generally increases N cycling and plant production (Gough et al. 2000, Elser et al. 2007), favoring species proficient in rapid aboveground biomass accumulation such as Agropyron repens over Schizachyrium scoparium in Minnesota old fields (Tilman 1987). |
| The experiment is located in an old field (Field C) of the Cedar Creek Ecosystem Science Reserve (CCESR, formerly Cedar Creek Natural History Area), approximately 45 km north of Minneapolis, Minnesota, USA, that was last cultivated to corn 48 years earlier (Tilman 1987). |
| Our observational study and experiments were conducted in a low soil-nitrogen, 4.1 ha old field (Field C) at CCESR that was abandoned from agriculture in 1934 and, via natural succession, came to be dominated by perennial grasses and forbs of the North American tall grass prairie (see Tilman 1987). |
| Little bluestem, Kentucky bluegrass, scribner's panicum (Panicum oligosanthes Schult.), bush clover, rigid goldenrod (Solidago rigida L.), gray goldenrod (Solidago nemoralis Aiton.), sheep sorrel (Rumex acetosella L.), and Pennsylvania sedge (Carex pensylvanica Lam.) are common herbaceous species (Tilman 1987). |

| |
|---|
| Seedlings of species of different successional statuses (light demanding/pioneers, shade tolerant/non-pioneers) may respond differentially to enhanced nutrient availability as they may vary markedly in their physiological and life history traits (Tilman 1987). |
| Despite the fact that P. pratensis is known to benefit from high N and associated N mineralization (Wedin & Tilman 1990) and can even displace tallgrasses under N addition (Tilman 1987), our findings cast doubt on whether the reverse is true. |
| Blackberry has long been an important colonizer of high-elevation forest gaps in GRSM (Crandell 1958), but its rate of colonization, density, persistence, and consequent capacity to inhibit forest reorganization on these sites may be enhanced by atmospheric-N deposition (Tilman 1987). |
| Grasslands are historically limited by nitrogen (N) availability and as a result, native plant species richness typically peaks at relatively low levels of soil N (Tilman 1987). |
| There are strong relationships between plant communities and environmental gradients such as climate (Weltzin et al. 2003), nutrient availability (Tilman 1987), topography/soil type (Whittaker 1960; Jones et al. 2008) and disturbance (Rao et al. 1990; Brockway & Lewis 1997; Ohmann & Spies 1998). |
| In nitrogen-limited terrestrial systems such as the Minnesota, USA, grasslands studied by Inouye et al. (1987) and Tilman (1987), early-successional annual forbs actually decline in abundance as old fields age and nitrogen accumulates in the soil (Tilman 1987). |

| Excerpts of text citing Huston and DeAngelis (1994) |
| --- |
| The overall similarity of the dynamics and life-history trade-offs indicates that native/exotic status has little influence on the performance of species in communities. These findings support the argument that native and exotic species are essentially drawing from the same pool of traits and therefore function within communities based on the same ecological rules (Huston and DeAngelis 1994). |
| Disturbances can contribute to resource heterogeneity by altering patch resource levels in contrast to the surrounding undisturbed matrix habitat (Huston and DeAngelis 1994). |
| Diversity patterns over the course of succession are known to vary widely and are strongly influenced by environmental conditions and the life-history characteristics of the specific organisms in question Huston and DeAngelis (1994). |
| Nutrient availability can influence plant performance and plant community composition during succession (Huston and DeAngelis 1994). |
| Study of the distribution patterns of living organisms and of the factors that they are driven by is a central theme that has stimulated a profusion of scientific research during recent decades (Huston and DeAngelis 1994). The large number of paradigms and theories proposed to explain spatio-temporal patterns of biodiversity at different scales attests to this interest, while it also underlines the complexity of the ecological processes involved. |
| Diversity is commonly related to disturbance severity (Huston and DeAngelis 1994). |
| Marquet et al. (2004) proposed the diversity deconstructive approach for detecting multiple response patterns that might otherwise be hidden by total richness (see also Huston and DeAngelis (1994)). |
| Stemwood production in forests appears to be more relevant than total plant production for evaluating tree growth rates and competition interactions as it is through the investment in the physical structure of wood (and also roots) that plants compete with one another (Huston and DeAngelis 1994). |
| Different hypotheses included equilibrium mechanisms through niche partitioning (Tilman 1994); non-equilibrium coexistence dynamics (Huston and DeAngelis 1994) related to disturbance (Connell 1978), predation and biotic interactions (Janzen 1970, Wills et al. 1997); fluctuations of environmental conditions (Chesson 2000); and balance between immigration/speciation and extinction (McArthur and Wilson 1967, Hubbell 2001). |
| Despite the fact that mature grassland is the dominant natural community type of 24% of the Earth's vegetated land area (Huston and DeAngelis 1994), most evidence of insect impacts on mature herbaceous communities is from areas generally dominated by forbs (e.g. Fraser & Grime 1997; Maron & Jefferies 1999; Carson Root 2000; Schdler et al. 2003). |
| Low-production successional environments, with low rates of competitive displacement and high diversity, should have high turnover while productive mature environments should have low turnover because their species are longer-lived and difficult to invade given their high rates of competitive displacement leading to dominance by only a few species and low diversity (Huston and DeAngelis 1994). |
| Spatial heterogeneity is thought to promote diverse communities (Huston and DeAngelis 1994). In many forest ecosystems, a few key species, referred to here as framework species, dominate the physical structure and function of the ecosystem. Framework species, also referred to as foundation species (Ellison et al. 2005) and structural species (Huston and DeAngelis 1994), provide the resources and microclimatic conditions used by interstitial species. They also restore functional characteristics suitable for the local environmental conditions and disturbance regime. |
| Species richness (total number of species occurring per unit area 1 m2), was used as an indicator of macrophyte species diversity (Huston and DeAngelis 1994). |
| This pattern may be explained following Huston and DeAngelis (1994)s suggestion that low levels of richness may be found in highly disturbed sites or in communities where strong competitive interactions result in species exclusion. |

| |
|---|
| The long-term maintenance and self-perpetuation of jack pine in the Moisie site is a direct illustration of an effective ecosystem process of recurrence dynamics over a period of several thousand years where classical successional trajectories (Huston and DeAngelis 1994) changing the vegetation assemblage were not operative. |
| Protected sites with P. juliflora having more richness may be due to localized site disturbance, which could enhance richness (Huston and DeAngelis 1994). |
| Light availability influences epiphytic establishment (Huston and DeAngelis 1994). |
| Such periodic flooding can maintain the resilience of these ecosystems by redistributing resources and inhibiting competitive exclusion (Huston and DeAngelis 1994). |
| A change in H was supposed to measure the effect of precipitation variability on the forest's community structure in different climate change scenarios, where pi is the proportion of individuals belonging to the ith species in the dataset (Huston and DeAngelis 1994):<br>$H = Ri = 1pi \times ln \times pi$ Equation(3)<br>Generally, the higher the index, the better the equipartition of species (Huston and DeAngelis 1994). |
| The term gradient means gradual unidirectional change of the parameter or change in discrete steps over that space (Huston and DeAngelis 1994). |
| Species diversity is measurable attribute of vegetation resulting from the combined effect of processes such as immigration, resource partitioning, competition, adaptation, speciation and extinction. The trade-offs in physiological and life-history characteristics cause the spatial patterns of species composition in response to environmental gradients (Huston and DeAngelis 1994). |
| Determining the causes of diversity gradient is extremely difficult (Huston and DeAngelis, 1994). |
| Although statistical analyses can never show causal relationship, they can often show strong correlations of diversity with factors that are marginally responsible for the diversity gradients (Huston and DeAngelis 1994). |
| Elevation and habitat moisture are examples of complex gradients. They have no direct effect on plant growth, but they are correlated with various factors that influence growth directly such as precipitation, temperature and solar radiation intensity (Huston and DeAngelis 1994). |
| A specific type of gradients are classified as complex because they influence the variation of other resource and regulator gradients (Huston and DeAngelis 1994). |
| There is evidence that species diversity of different growth forms can vary in different (opposite) manner along the productivity gradient for example, decreasing of tree diversity versus increasing grass and herb diversity (Huston and DeAngelis 1994). |

Excerpts of text citing Grime et al (1988)

| |
|---|
| At the CZ site, two sown species that successfully established and became abundant in plots where they were sown were Trisetum flavescens with very light seeds (0.18 mg Grime et al 1988) and Lathyrus pratensis with heavy seeds (12.85 mg). |
| However, in this study anemochory and zoochory (Grime et al 1988) were the most frequent dispersal mechanisms even in the early stages of succession, similar to results found in secondary successions on semi-arid Mediterranean old-fields (Bonet & Pausas 2004), although there are a number of species that do not possess specific dispersal modes (Grime et al 1988). |
| Persistent seeds are present in the seed bank for more than one year versus transient seeds that are present for less than one year (Grime et al 1988). |
| Molinia also has a wide pH tolerance (Taylor et al. 2001) and is classified as a stress tolerant competitor (Grime et al 1988). |
| Groups were formed on the basis of the strategies outlined by Grime et al (1988), i.e. ruderals (R), stress tolerators (S) and competitors (C), which is a general benchmark for plant strategy types and plant functional types (van der Maarel 2005). |
| All strategies were identied using Grime et al (1988)s determina-tion key (Grime et al 1988). |
| Furthermore they were classified to CSR-strategy according to Grime et al (1988): c=competitor, s=stress-tolerant, r=ruderal (cr, cs, sr and csr are intermediate strategies). |
| The types of strategies were determined chiefly on the basis of the list by Hunt et al. (2004), for species absent on this list the key given by Grime et al (1988) was used. Several factors affect the buoyancy of seeds, such as seed shape (defined as length/width: Grime et al 1988). |
| Cover weighted averages of the indicator values given by Ellenberg et al. (1992) and calibrated C-S-R strategy types by Grime et al (1988) were calculated for each sample. Calibration of unbalanced C-S-R radii for species was performed according to Hlzel (2003). Only species categorized by Grime et al (1988) were included in the analysis; these comprise about 70% of the entire species pool and 90% of the most frequent and abundant species. The generation of C-S-R values for species not mentioned in Grime et al (1988) was explicitly avoided. Instead, species without C-S-R values were made passive in the calculation |
| In the course of the vegetation development, agrestal and ruderal species with a high production of small seeds were increasingly suppressed by grassland species with lower seed production but larger seeds (Grime et al 1988). |
| The long-term monitoring of vegetation in permanent plots on Surtsey showed that it was mainly ruderal species (sensu Grime et al 1988). |
| Reactions to disturbance differ since species are adapted in distinct ways to these impacts. Species groups can be differentiated by their C-S-R strategies. The triangular model of primary ecological strategies (Grime et al 1988) discriminates strategies of competitiveness, stress tolerance and ruderality using resource availability and disturbance as two orthogonal dimensions for plant classication. |
| Abundance-weighted (i.e. weighted for seed density) calibrated C-S-R strategy types by Grime et al (1988) were calculated for each plot based on the species found in the soil seed bank. |
| Only species categorized by Grime et al (1988) were included in the analysis when considering C-S-R as dependent variable. |
| Meadows and meadow-pas-tures contained a number of indicator species characteristic of relatively nutrient poor site conditions such as Luzula campestris and Potentilla erecta which are considered as stress tolerators (Grime et al 1988). |

| |
|---|
| In contrast to silage meadows and permanent pastures, hay meadows and meadow pastures are characterized by a higher proportion of the stress strategy, which probably reects a lower degree of disturbance and a lower nutrient availability. This is best exemplied by the occurrence of certain stress tolerant grassland species such as Luzula campestris, Pimpinella saxifraga, Festuca rubra and Potentilla erecta (Grime et al 1988). |
| Plant species with very small seeds were possibly lost during the washing process. In our locality J. effusus is such a species (seeds 0.01 mg, 0.50.3 mm2 large; Grime et al 1988). |
| This is probably a consequence of heavy herbivore pressure, as L. pratensis is a relatively large seeded species (seed weight 12.85 mg; Grime et al 1988) and indeed, we have often found half eaten pods on Lathyrus plants in our site. |
| Again, species as sedges, T. dubium or A. reptans have generally round shaped seeds, their size varying from 0.32 mg for T. dubium up to 1.88 mg for C. panicea ( Grime et al 1988). |
| Several authors already have mentioned the broad range of soil conditions tolerated by F. japonica. In the UK, it has been found on soil with pH ranging from 3.0 to 8.0 (Grime et al 1988). |
| MAVIS was also used to indicate the environmental conditions characterising each community via Ellenberg's indicator values (light, moisture, pH and fertility), modified for British conditions (Hill et al., 1999) and the type of plants and their life strategy through the functional strategy theory (Competitor, Stress-tolerator, Ruderal characterisation (CSR)) (Grime et al 1988). |
| F. sylvatica does not establish from seed on open ground (Rackham 1980; Willoughby et al. 2004), but is tolerant to shade (Grime et al 1988). |
| Ecological strategy follows the C (competitive species)-S (stress-tolerant species)-R (ruderal species) strategy types by Grime et al (1988). |
| Ecological strategies are defined according to Grime et al (1988). C = competitive species, R = ruderal species, S = stress-tolerant species. |
| Less aggressive colonizers such as Quercus (Quercus petraea Liebl. and Quercus robur L.) and Fagus sylvatica L. do not produce seed until the tree is at least c. 40 and c. 28 years of age, respectively (Grime et al 1988). Filipov and Krahulec (2006) considered A. alpinum as a S strategist (sensu Grime et al 1988) but its flexible response to fertilisation is a trait rather than a characteristic of other strategies. |
| Herbs that expanded most after clear-felling had a functional strategy with an important C-coordinate, indicating that they are adapted to a relatively low stress level (Grime et al 1988). |
| Many forest herbs require a certain minimum quantity of light or a minimum temperature to germinate (Grime et al 1988). |
| Twelve species of flowering plants were selected whose seeds were classified as showing a small proportion of seeds as persistent in the soil (type 3 of Grime et al 1988). |
| There were clear differences in the behaviour between species. As might have been expected from the classification in Grime et al 1988), the number of viable seeds of Anthoxanthum odoratum declined rapidly. |
| The C-S-R signature (Grime et al 1988) for each plot was calculated by means of C, S and R values. |
| Maple is a relatively common species in the area; it usually grows at borders, in hedgerows and along roads. It is also anemochorous, and its seed can be blown for long distances. It recruits very well in different habitats (Grime et al 1988). |
| Seedlings of mountain maple were more abundant in younger growths, which grew fast; in other words, the stand was more rich in nutrients. The ecological preferences of the species correspond with this finding (Grime et al 1988). |
| The better retention of sown species in mown plots may result from the selection of more stress tolerant species compared to competitors (sensu Grime et al 1988). |

| |
|---|
| Few common perennials were significantly affected by mowing during the establishment phase, but after 13 years the most competitive species (sensu Grime et al 1988) were more abundant in the absence of mowing, while most of the sown species, typical of semi-natural grassland, fared better, often by degree, when mown. |
| Strategy details the CSR functional strategy: competitive (C), stress-tolerant (S), ruderal (R) or a combination of them, following Grime et al (1988). |
| The most effective seed response to wild boar disturbance was found in Agrostis capillaris and Festuca gr. rubra, which can be considered as polyvalent in terms of the functional strategy they adopt, since they can behave as competitors, stress-tolerant or ruderal depending on environmental conditions (Grime et al 1988). |
| The positive correlation of the ruderal ecological strategy, whose particular distinctive features include a short ontogeny or a high reproductive effort (Grime et al 1988), with the avoidance of mycorrhizal symbiosis has been discussed previously (Grime et al (1988); Francis and Read, 1995; Cornelissen et al., 2001; Betekhtina and Veselkin, 2011). |
| The grass Festuca rubra, a species associated with disturbance (Grime et al 1988), was more abundant near burrows where it may have benefited from vole-derived nutrients and the suppression of Molinia (Herben and others 1994; Lep 1999). |
| The positive response of Festuca rubra (see above) and rushes, species associated with disturbance (Grime et al 1988), to water vole occupancy, and the overall declines in the density of disturbance intolerant grass in more burrowed areas hint at the importance of the physical process such as burrowing and soil deposition. |
| First, germination of Juncus effusus is strongly light dependent and therefore suppressed by dense plant cover (Agnew 1961). It is frequently amongst the first species to establish on peaty soil bared by disturbance (Grime et al 1988). Second, Juncus prospers under wet conditions but requires aeration for at least part of the year despite its aerenchyma (Grime et al 1988). |
| Raunkiaer's life form spectra (RLFS) (Raunkir, 1934) places species into groups based on the plant characteristics such as morphology, namely the architectural trait height of perennial buds.[...]Since RFLS groups are based upon characteristics associated with considered survival strategies (Grime et al 1988), the proportion of each life-form present in a community indicates not only how species adapt to climatic conditions but also how strong the possible stressors are and how they impact the community. |
| Some common and dominant wetland species, such as cattails (Typha sp. div.) and common reed (Phragmites australis), produce light seeds, which are easily wind-dispersed (Grime et al 1988). |
| Vegetation functional traits were also analysed (Table 2), including species of ancient woodland habitats in the UK (Kimberley et al. 2013), established plant strategies (Grime et al 1988), life forms, clonality and Ellenberg indicator values based on their proportionate cover values (Hill et al. 2004). |

Excerpts of text citing Noble and Slatyer (1980)

| |
|---|
| The identification of habitat resources that develop over many decades, and the documentation of associated developmental patterns, can inform fire management by identifying minimum and maximum fire intervals for fauna, as undertaken using plant species attributes (e.g. age at first seed set, longevity: Noble and Slatyer 1980). These different spatial and temporal patterns of disturbance are likely to have acted as profound selective forces resulting in organisms with life-history attributes that enable them to cope with a particular range of disturbances, e.g. fire regimes, floods and droughts (Noble and Slatyer 1980). |
| Nevertheless, the premise that an organisms capacity to regenerate or re-colonise following fire will be limited by the organisms own life-history attributes (e.g. age at sexual maturity, reproductive rate, 210 dispersal ability) seems inescapable. Therefore, much effort has been made, to identify what Noble and Slatyer (1980) labelled the Vital Attributes of organisms, largely plants. |
| This may make analysis numerically possible in the absence of data for certain communities, but whether the recommendations arising from this application of Noble and Slatyer (1980)s method have any ecological validity is debateable. |
| Second, data on the length of time that seeds remain viable within a soil seed bank is often lacking in the vital attributes data-base for many species (Cheal 2004), despite Noble and Slatyer (1980) recommending that soil seed-bank longevity data were an essential component of the minimal data set needed to apply their method. |
| Generalities across ecosystems, or mechanisms underlying differences, have remained elusive. However, one promising tool is the analysis of plant functional traits (or vital attributes) autecological qualities with common response to the environment across taxa. Noble and Slatyer (1980) described a qualitative scheme for predicting shifts in plant communities subject to recurrent disturbance, using functional traits relating to regenerative strategy (e.g. seed longevity, dispersal capacity, sprouting ability) and competitive relations (e.g. growth rate, shade tolerance). |
| By assessing logically determined interactions between functional groups and fire interval, this approach provides a broadly applicable framework for understanding the role of disturbance frequencies in the origin of different successional pathways (Noble and Slatyer 1980). |
| To elucidate mechanisms and improve predictive capability across ecosystems, we explored the question: What plant functional traits are associated with the different fire histories (SI fire, LI fire, mature/old-growth (M/OG) stands with no recent fire)? We focused on broadly applicable traits (related to those in Noble and Slatyer 1980) including regenerative strategy, life form, and dispersal strategy. |
| Fire Effects Information System http://www.fs.fed.us/database/feis, which summarizes functional traits relevant to post-fire regeneration and development for each species. These traits overlap broadly with the vital attributes described by Noble and Slatyer (1980). |
| The mechanisms for increases in early seral species and total richness in the SI burn could be abiotic, such as changes to soil properties that favour such species, or biotic, as in the development of a propagule bank for early seral species during the 15 years between fires (Noble and Slatyer 1980). |
| Assessing Noble and Slatyer (1980) vital attributes against the 15-year fire interval, forbs and low shrubs were G types (rapid maturation time and stored soil seed banks), hardwoods and shrubs were S and/or V types (vegetative sprouting ability and/or long-lived soil seed banks), and conifers were D types (well-dispersed propagules from surrounding live tree sources). |
| Thus, species investing more resources in early reproduction, and less in long-lived leaves and secondary tissues (wood), may be expected to be relatively dominant immediately following recurrent stand-replacing fires (Noble and Slatyer 1980). |

| |
|---|
| Post-fire succession can be divided into two phases (Noble and Slatyer 1980): the first, immediately post-fire, when competition for resources is low and species abundance is driven primarily by regenerative processes (the focus of this study); and the second, after this initial pulse, when resource competition becomes progressively important. Species availability encompasses the processes that determine the ability of species to disperse into a recently disturbed site or survive a disturbance as propagules (Noble and Slatyer 1980). |
| A number of classification systems for plant adaptions exist, including some specifically related to fire. Noble and Slatyer (1980) have presented a general model of plant strategies in respect to fire, focusing on regeneration or colonization attributes; application of this model to a community, however, requires detailed information on life cycle, persistance and dispersal abilities of the species present. |
| An ability to predict the occurrence of species is critical in regions prone to frequent disturbance, particularly when such disturbances are amenable to management (Noble and Slatyer 1980). |
| A number of studies have highlighted strong relationships between plant life history attributes and sequences of fires (e.g., Noble and Slatyer 1980). |
| Outcomes are often site specific, and it is possible there may not be a set of vital attributes (sensu Noble and Slatyer 1980) that determines responses for animals as there has been hypothesized for plants. |
| To analyse trait responses on any environmental gradient, a classification of traits into the fundamental stages in the life-cycle (ecological performance) of plant species is helpful. Weiher et al. (1999) classified the traits of their core list into dispersal, establishment and persistence. Other classifications are (1) the vital attributes scheme of Noble and Slatyer (1980), (2) competitive ability (Gaudet and Keddy 1988) and dispersal ability (Knevel et al. 2003, Rmermann et al. 2005) or into (3) adult traits and seed traits (Fenner and Thompson 2005). |
| Sprouting from stems and roots has been reported to be common after disturbance (Uhl et al.1988; Putz and Brokaw 1989; Kauffman 1991) but the regrowing vegetation depends on various regeneration sources (Noble and Slatyer 1980). |
| Some of the variability in successional models and theory is due to a failure to define the temporal phase or context (i.e., within versus among life forms) of succession (Grace 1991). For this purpose, Noble and Slatyer (1980) recognized two temporal phases of succession: (1) the immediate post-disturbance phase (initial state) when species abundance is determined by regenerative processes, and (2) later succession when species interactions and resource competition determine abundance. |
| Initial and relay floristics models of succession (Egler, 1954) are useful for understanding early succession in managed forests because these models (1) focus on patterns of initial regeneration that may be evident in the immediate post-disturbance phase of succession (Noble and Slatyer 1980), (2) predict measurable shifts in diversity and abundance, and (3) are sufficiently descriptive to be useful for developing forest management strategies (Egler 1954; Kimmins 1997). |
| Community assembly immediately after fire is primarily governed by fire survival and post-fire plant regeneration strategies (reflected in species traits) in a relatively competition-free environment (Mallik and Gimingham 1985). In time, plant interaction becomes a dominant force, as available space for regenerating species is depleted (Noble and Slatyer 1980). |
| Other modelling approaches also use suites of morphological and regenerative traits to simulate the assembly of communities. A classical model is the vital attributes model from Noble and Slatyer (1980) with several derivates (Moore and Noble, 1990 ; Lavorel, 2001). Conceptual models of succession following fire have been derived for vascular plants. These models incorporate the adaptive traits that plants have to survive fire (Gill 1977, 1981) into a small set of attributes vital to the reproduction and recovery of a species following fire (Noble and Slatyer 1980). |

| |
|---|
| The objectives for this study were to (i) record the species of macrofungi occurring in recently burnt karri regrowth forest (based on the presence of sporophores); (ii) monitor the change in species abundance, diversity and composition over time; and (iii) compare the post-fire mycoflora in recently burnt regrowth forest with that on similar aged regrowth forest that was unburnt since establishment. We also consider the application of vital attributes (Noble and Slatyer 1980) to macrofungi and assess their reliability in predicting the successional sequence in a macrofungal community following fire, and how they may also be utilized in determining appropriate fire regimes for continued survival of species. |
| A third qualitative model by Noble and Slatyer (1980) appears even more applicable, as it concerns succession in plant communities that are susceptible to recurrent disturbances, and emphasizes the properties of individual species. The three key features that the model aims to consider are: (1) the method of species arrival, or persistence, after a disturbance; (2) the ability of species to establish and grow in the postdisturbance community; and (3) the time taken by species to reach critical life stages. |
| The boreal forest is highly dynamic and canopy change, although being predictable by tree life history strategies (Noble and Slatyer 1980), is greatly influenced by disturbances that occur throughout the course of succession (Bergeron and Charron 1994, Kneeshaw and Bergeron 1998). |
| A shift to an ecophysiological view in ecology, driven by Hutchison and Eugene Odum, took a functional approach to succession (Drury and Nisbet, 1973), with Noble and Slatyer (1980) attributing vital attributes to species, wherein species-environment interactions, rather than the developmental anatomy of the community per se, drove community change. |
| The variation in species characteristics along the succession is important because they change the systems function. Stages in successions have long been described through the biological characteristics of the dominant species (Noble and Slatyer 1980). |
| Fire affects vegetation via the attributes of individual fire events, such as fire severity or time since fire; as well as by the sequence of fire events through time, the fire regime (Gill 1975, Gill et al. 2002). For example, the severity of fire events determines the degree to which vegetation structure is altered (Smucker et al. 2005). Following fire, vegetation then undergoes successional changes (Noble and Slatyer 1980). |
| The length of time between fires (fire interval) can influence the persistence of individual plant species by affecting whether reproductive maturity has been reached at the time of fire (Noble and Slatyer 1980) and cause major and long-lasting change to vegetation structure and composition (Zedler et al. 1983). |
| The wide variation in species response to Time likely reflects variation in species relative growth rate, longevity and dispersal mechanism. These are important mechanisms driving secondary succession of vascular plant communities in favourable environments (Noble and Slatyer 1980) and we assume the same is true for biocrust communities. |
| The observed declines in abundance of all ruderal species in early stages of succession are likely due to species competition for space and light (Noble and Slatyer 1980) and resources such as soil nutrients (Tilman 1985). |
| Therefore, the secondary succession among the shrub and herb species was determined by the vital attributes of the species, i.e., the attributes of a species that are vital to its role in a vegetation replacement sequence (Noble and Slatyer 1980). The three most important vital attributes recognized are: (1) the method of arrival or persistence of the species at the site during and after a disturbance; (2) the ability to establish or grow to maturity in the developing community; (3) the time taken for the species to reach critical life stages. |

Excerpts of text citing Tilman (1985)

| |
|---|
| This finding provides support for the resource ratio hypothesis of plant succession, i.e. species that can grow at the lowest resource level tend to dominate resource-limited sites throughout succession (Tilman 1985). |
| Species performance contains all of the mechanisms by which species interact and sort themselves within a community. This class has received the most attention and therefore contains the greatest diversity of potential successional drivers. It covers Tilman (1985)'s competition-based resource ratio hypothesis (Tilman 1985), the interaction-based views of Connell<br>Slatyer (1977) and even Clements (1916), as well as the majority of trait-based sorting suggested by Grime (2001) |
| Tilman (1985)s formulation of the resource ratio hypothesis focuses on two plant resources: light availability at the soil surface and nutrient concentrations in the soil. Tilman (1985) viewed successional changes as a gradient from high availability of light and resource-poor soils in the beginning, to nutrient-rich soils and low availability of light later in succession. |
| We addressed the following questions: (1) do our findings support the resource ratio hypothesis of Tilman (1985) and if so (2) how quickly do resources change over time? (3) do our findings confirm the initial floristic composition model by Egler (1954) and thus (4) does the initial treatment of the plots, through differences in plant species composition, have lasting effects on nutrient supply, soil organic matter or light availability? |
| Our study plots changed from open herbaceous to closed pioneer forest within relatively short time periods. Therewith light supply on the ground dramatically decreased over time. Nitrogen showed a clear accumulation in the upper soil but not in total, the C/N ratio significantly increased over time, and phosphorus showed a slight decreasing rate. Thus our results do not completely fit the resource ratio hypothesis proposed by Tilman (1985), but rather emphasise light as being a main influencing factor for vegetation development during succession. |
| An important result was that the total species richness peaked in the early stages of succession, 1015 years after reclamation started. This early peak in total species richness is similar to those documented in other studies of old field successions in Mediterranean climates (Tatoni et al. 1994; Bonet and Pausas 2004) and is consistent with the resource-ratio hypothesis (Tilman 1985). |
| Tilman (1985)s model, which was initially developed to explain the course of plant succession and, in particular, the dominance of species (Tilman 1985), establishes the link between alien and native species invasions. |
| Competition for nutrients plays a major role in governing grassland plant community dynamics (Tilman 1982). Adding a single nutrient alters the absolute abundance of that nutrient, as well as its abundance relative to other nutrients (e.g., N:P), often shifting the identity of the limiting nutrients and ultimately plant community composition (Tilman 1985). For instance, enriching soils with N favors N-limited species, and increases plant productivity at the expense of diversity (Tilman 1987, Foster and Gross 1998). |
| Fundamental theories such as those that link succession and the use of resources by plants (e.g., the resource-ratio theory; Tilman 1985), though extensively tested in laboratory or microcosms studies, still lack empirical validation from field experiments (Miller et al. 2005). |
| In the context of succession, resource-ratio theory (Tilman 1985) predicts that resource supply determines whether competing species can coexist and, if not, which species will exclude others. It also predicts that the highest diversity of competing species should occur at intermediate resource availability. |
| That aboveground competition was important for biogeochemically mediated feedbacks is not surprising given that plant neighbors can switch the limiting resource from soil nutrients to light (Tilman 1985). |
| Clarkson (1990) and Walker and del Moral (2003) present reviews of the application of various models (sensu Connell and Slatyer 1977) and mechanisms (sensu Finegan 1984) of succession in the development of montane plant communities on volcanic terrains similar to that studied here, and Clarkson (1990) has also framed the development of vegetation on mineral surfaces in relation to Tilman (1985)s resource ratio model of succession as a gradient of declining ratios of solar radiance to nutrient availability. |

| |
|---|
| In general, successional changes on old fields have been interpreted in terms of competitive ability mediated by resource availability, in particular light and nutrients (Tilman 1985). |
| The evolutionary consequence of nutrient limitation on shade-tolerance is predicted by two hypotheses that are mutually exclusive: the resource ratio hypothesis (Tilman 1985) and the growth-survival trade-off hypothesis (Fine et al. 2006, Russo et al. 2007). Tilman (1985) hypothesized that species with an ability to use one resource efficiently use the other resources inefficiently. If so, plants adapted to the environments with severe nutrient limitation inherently show lower shade-tolerance because they cannot enhance abilities to utilize both nutrients and light efficiently. |
| We determined the responses of shade-tolerance (defined as survivorship at 5% GSF) to different levels of nutrient limitation. The resource ratio hypothesis (Tilman 1985) and the growth-survival trade-off hypothesis (Fine et al. 2006, Russo et al. 2007) predict the contrasting patterns of shade-tolerance in relation to nutrient availability. All species in the severely nutrient-poor site (tropical heath forest) showed greater shade-tolerance (over 91% survival for 8 mo) than species in the other sites (5487% survival for 8 mo) (Figure 1, Appendix 4), which supports the growth-survival trade-off hypothesis. |
| The nutrient limitation hypothesis (Tilman 1985) predicts that whichever of these is in shortest supply within a given environment is the limiting compound. When this limiting element or compound is added, it would logically increase the amount of CH4 oxidation that can be performed, until saturation. If the ratio of existing available inorganic N to available CH4 to which methanotrophs have access determines the rate of CH4 oxidation of the system, then when this ratio is low any addition of N may stimulate increased CH4 oxidation. |
| Many models have been developed to understand and predict population dynamics, but with the exception of several models of mutualisms (e.g. Holland, DeAngelis<br>Schultz 2004) most have focused on negative interactions such as predation (e.g. Polis<br>Strong 1996) and competition (e.g. Tilman 1985) |
| By building on the work of Tilman (1985), Norberg et al. (2001) and Enquist et al. (2015), we predict that changes in resource availability during succession would select for traits associated with different physiological and life-history strategies along the fast-slow trait continuum. |
| Building on the work of Tilman (1985), Norberg et al. (2001), and Enquist et al. (2015), we hypothesized that for TDFs (tropical dry forests) community functional composition during succession will shift from more xeric and filtered slow strategies towards more mesic fast strategies. |
| These results may be attributed to the species sequence observed in the secondary succession in rich soils, which include the initial dominance by early successional species, with a replacement of these species by late successional species (Tilman 1985). The change in dominant species likely resulted in changes in the growth patterns of other species and their resource acquisition strategy (Tilman 1985). |
| Although community assembly studies typically focus on individual species (e.g. Tilman 1985) at relatively small scales (Leibold et al. 2004), the same suite of processes that affect biodiversity can influence patterns of vegetation structure, quality, or chemical composition (McGill et al. 2006), which can be estimated spatially via remote sensing. |
| According to the resource-ratio hypothesis (Tilman 1985), one of the species should be a superior competitor for a particular proportion of the limiting resources and this balance shall predict the community composition. |
| Furthermore, species responded differently to N and/or water addition, indicating differential resource requirement or limitation among species. For example, biomass of L. chinensis increased after N addition, but did not respond to water addition, while biomass of A. frigida was more responsive to water addition than to N addition (Fig. 2), indicating that L. chinensis was mainly limited by N, while A. frigida was limited more by water availability. Our results support the resource ratio hypothesis (Tilman 1985) that steppe species have different resource demands and show differential responses to water and N additions (Xia and Wan 2008, Yang et al. 2011). |

| |
|---|
| According to resource-ratio hypothesis (Tilman 1985), if the most limiting resource is added, species will have greater competitive abilities. Our results suggest that, early successional species were limited most by water while late-successional species were limited most by N. |
| Fields and forest gaps developing from an open, disturbed state to closed-canopy forest tend to increase in tree richness and diversity, while decreasing in ground-layer cover and diversity (Tilman 1985). |
| Low root-shoot ratios of stem cuttings of E. nigrum and J. communis in organic matter mulch and control may also suggest higher allocation of energy to lateral shoot increment. In contrast, higher root-shoot ratio of the cuttings in hay-mat mulch may suggest that these cuttings allocated more energy to root elongation in search of moisture and nutrients (Tilman 1985). |
| The observed declines in abundance of all ruderal species in early stages of succession are likely due to species competition for space and light (Noble & Slatyer 1980) and resources such as soil nutrients (Tilman 1985). |
| Although the grasslands in our study system represent a succession that is truncated by grazing management, our results are nevertheless in agreement with the predictions of general models of succession, such as the resource-ratio hypothesis (Tilman 1985) and the tolerance model (Connell & Slatyer, 1977). If the typical grassland species that have their highest frequencies in the mid-successional grasslands have a somewhat higher requirement for nutrients than other typical grassland species, then the models predict that these species should be relatively less frequent in the nutrient-poor, old grasslands. |
| Tilman (1985) presented the resource-ratio hypothesis, which assumes that each plant species is a superior competitor for a particular proportion of the limiting resources. According to the hypothesis, species composition in a community should change whenever the relative availability of limiting resources changes. Because limiting resources such as light and soil nutrients change through succession in general, the hypothesis predicts the replacement of species over time. The relative availability of limiting resources likely changed in the study forest during secondary succession. |

Excerpts of text citing Shugart (1987)

| |
|---|
| Regarding basic research, species-specific growthmortality relationships could be implemented into forest succession models, e.g. gap models (Shugart 1984) |
| Regeneration filters (sensu Shugart 1984) include winter temperature, light, browsing, degree-days, and immigration. |
| The year 1999 had slightly higher values for all forest variables (records prior to 1981 are not available), and could be due to the forest being in a dynamic equilibrium and the non-equilibrium state of vegetation (Shugart 1984), and all percent errors comparing 1999 observed data and modeled results were high. |
| Forest succession models from the family of gap models (Shugart 1984) have been used widely in applied research to examine |
| Gap models typically consist of sub-models describing tree growth, allometry, mortality, and recruitment as driven by individual tree characteristics, competition for light or other resources, and local environmental conditions. They have long proven successful in representing population dynamics and disturbance impacts in complex, multi-species forest stands (Shugart 1984), and recently have been extended to predict geographic distributions and succession at regional scales (Purves et al. 2008, Vanderwel et al. 2013a). |
| Forest gap models have been used widely to simulate the succession of mixed-species forests in a changing environment (Porte and Bartelink, 2002; Shugart, 2002) and their structure is well described (Shugart 1984). |
| Capacity to project changes in species composition in naturally regenerating forests following anthropogenic disturbances has been a key area of interest in the application of gap-based models to forest succession (Shugart 1984). |
| The field of forest modeling was for a long time focused on niche-based, resource-mediated interactions among individuals (e.g., Shugart (1984) and derivatives, Botkin, 1993). |
| Phipps (1979) assumed that all species had the same mortality response to low growth rate (a common assumption shared by most gap models, e.g., Shugart 1984). |
| The concept of forest gap models (Shugart 1984) deviates strongly from that of forest growth models in the sense that they are formulated more generally and usually do not depend on site-specific parameterizations. |
| Forest gap models have been used widely to examine forest succession as a complement to field observations and experiments ( Shugart 1984). However, being adapted to a relatively broad spectrum of environmental conditions, these models are most appropriate for simulating forest succession under general temperate and boreal conditions, for which they were originally developed. |
| Gap models account for competition among individuals of multiple tree species for light and other resources, with the outcome determining the composition and structure of the forest through aggregation of homogenous mosaic patches through time (Shugart 1984). |
| In forest succession models that are based on the gap dynamics paradigm (Botkin et al. 1972; Shugart 1984), the life history strategies of tree species are captured via sets of parameters that are either species specific (e.g., longevity or maximum diameter increment rate) or relate to functional groups (e.g., shade tolerance). |
| The fundamental growth/survival tradeoff that allows tree species to either establish in gaps or persist under deeply shaded understories is a basic tenet of the shade-tolerance paradigm (e.g. Shugart 1984). |
| Ecological process models are important tools for studying past and present ecosystem dynamics and are a key source of information about possible future changes. Of these, forest gap models simulate individual tree establishment, growth and mortality in response to environmental conditions, disturbances and competition on a large number of small, separate patches (e.g., Shugart 1984). Gap models have recognized limitations (e.g., Bugmann et al., 1996; Bugmann, 2001; Nalder, 2002) but are particularly well-suited to investigating individual species responses to local conditions and the influence of these on community and landscape level dynamics (e.g., Shugart and Smith, 1996; Waring and Running, 1998; Smith et al., 2001). |

| |
|---|
| Tree mortality in the EDS (Ecosystem Dynamics simulator) model occurs as a combination of age-related and stress-induced death rates. Each tree has an inherent risk of natural death likely to occur to any healthy tree from lightning strikes, storms, fire and other random causes. Small trees are subject to high mortality due to strong competition for resources and old trees to high mortality due to diminishing growth vigour. Trees growing poorly due to adverse environmental conditions are subjected to stress-induced mortality (Shugart 1984). |
| Height estimation in most gap models is based on a relationship between dbh and total tree height that uses species specific parameters b1 and b2 (Shugart 1984):<br>$H(height) = datum + b1dbhb2(dbh)2$<br>where datum is the height (cm) at which tree diameter (dbh, cm) is measured. |
| In contrast to gap models which track succession dynamics for individual trees (e.g., Shugart 1984), LANDIS (model) treats a landscape as a grid of sites (or cells) with vegetation information stored as attributes for each cell. |
| Individual-based forest successional models (i.e. gap models) provide a tool to explore thresholds at which a forest enters a certain successional stage. Gap models simulate the fate of single trees on the basis of species life-history traits and limited resource availability (e.g. light, Shugart 1984), thereby facilitating detailed analyses of forest successional trajectories. |
| The model simulates forest dynamics as a mosaic of interacting forest patches of 20 m 20 m, which is the approximate crown size of a large mature tree. Within these patches, forest dynamics is driven by tree competition for light and space following the gap model approach (Shugart 1984). The competition for light is modeled by dividing each patch vertically into height layers. In each height layer the leaf area is summed up and the light climate in the forest interior is calculated via a light extinction law. |
| Age-related or natural death of tree species was estimated using the natural mortality equation found by (Shugart 1984). |
| The tree height was calculated from the relationship $H = 137 + b2DBHb3DBH2$ (Shugart 1984). |
| This result clearly shows that the often-considered biomass (Shugart 1984) is not sufficient to characterize spatio-temporal patterns of forest structure resulting from disturbance regimes. |
| Regenerating gaps would have a particular role in terms of providing seeds for newly created patches according to existing species communities and stand structures (Shugart 1984). |
| Gap models are based on the forest dynamics involved in the competitive aftermath of the death of a large, dominant tree (Shugart 1984) and are able to simulate small-scale tree responses to their environment, climate, and disturbances, tree to tree competition, as well as larger-scale successional dynamics. |

| Excerpts of text citing Guariguata and Ostertag (2001) |
| --- |
| In young forests, before density-dependent forces become strong enough to regulate tree distributions, a high density of stems may recruit and establish (Guariguata and Ostertag 2001). |
| Fast increments in height and crown cover compared to basal area agree with processes of early allocation to resource acquisition followed by a later shift toward structural materials (Guariguata and Ostertag 2001). |
| In TRF (tropical rain forest), this second phase of pioneer trees is replaced, within 1030 yr, by a group of long-lived pioneers that dominate the forest during 75150 yr before mature forest species do so (Guariguata and Ostertag 2001). |
| A small set of effective pioneer species without long life spans leaves less room for compositional variation and thus may lead to earlier convergence between secondary and mature forests. Such convergence would not take place before 40 yr of regrowth but is possible for it to occur earlier than the 75150 yr estimated for TRF (Finegan 1996, Guariguata and Ostertag 2001). |
| Species number recorded on the same plot size as in our study (900 m2) increased from an average of 45 tree species (¿1 cm dbh) after 12 years since the start of succession to 71 species after 80 years. Similar values have also been reported from Neotropical secondary forests (Guariguata and Ostertag 2001) or from Japan (Aiba et al. 2001). |
| Basal area and soil litter, which change more gradually than vegetation cover (Guariguata and Ostertag (2001), Lebrija-Trejos et al. 2008), also modify the understorey environment. |
| Along these seral stages, pioneer plants are replaced by shade-tolerant trees and the tree assemblage gradually accumulates biomass and becomes increasingly more diverse in both taxonomic and ecological terms (i.e. higher diversity of plant life-history traits irrespective of their taxonomic composition; Guariguata and Ostertag 2001). |
| Increase in light availability favor shade intolerant species and decrease the number of understorey species in secondary forests (Guariguata and Ostertag 2001). |
| During the development of a plant community following a secondary succession there is an increase in the amount of species with larger individuals, which in turn leads to stratification (Guariguata and Ostertag 2001) and structural complexity. |
| There, basal area peaks early or midway during succession because early suppression of regeneration produces a time-lag between the mortality of the dominant pioneer cohort and the regeneration of late colonizers (Guariguata and Ostertag 2001). |
| We recognize the important role of seed dispersal and dispersal limitation in community assembly during secondary forest succession (Guariguata and Ostertag 2001). |
| Our estimated time required for above-ground biomass to reach approximately 85% of undisturbed forest levels is similar to suggested rates for basal area recovery in the neotropics [46]. While our results and previous observations [46] suggest that forest biomass approaches that of undisturbed forest within a century, full recovery may take substantially longer. |
| Secondary tropical forest tree communities are initially dominated by short-lived pioneer tree species, and these are sequentially replaced by longer-lived species [46]. Some secondary forests may be isolated from seed sources, leading to an impeded recovery of richness, but our results, and the observations of others [46], suggest that this is relatively rare. By contrast, epiphyte dispersal is largely local and propagation is often restricted to individual trees [58]. In addition, epiphytes seem to occur more commonly on large trees [59]. These factors may lead to relatively poor recovery of epiphyte species because many secondary forests are fragmented and tend to consist of smaller-stemmed trees [46]. |
| From chronosequence studies, it appears that regenerating forests can attain many of the structural characteristics of OG and equivalent woody plant species richness within 2030 yrs (Guariguata & Ostertag 2001), but carbon storage may not attain old-growth levels until many decades later (Fearnside & Guimares 1996; Denslow & Guzman 2000; Mascaro et al. 2012). |

| |
|---|
| Rapid recovery of species richness depends on unlimited seed availability and dispersal (Guariguata and Ostertag 2001), and the configuration of the BCNM allows for high connectivity and seed dispersal among forest patches. |
| Newly germinating colonizers must tolerate high soil temperatures and dry conditions in upper soil layers (Pineda-Garca, Paz & Meinzer 2013), potentially compounded by decreases in soil water-holding capacity post-deforestation (Guariguata and Ostertag 2001). |
| Slash and burn conversion of forests into pastures can homogenize the environment through soil compaction, applied fertilization, and post-disturbance cultivation of non-native grasses (Guariguata and Ostertag 2001). |
| Changes in habitat structure due to human activities can alter environmental conditions such as exposure to light, humidity, and quantity of leaf litterfall (Guariguata and Ostertag 2001). |
| Three of these habitats are of focal dispersal interest for conservation and/or restoration efforts: forest remnants that harbor the majority of remaining biodiversity and function as sources of seeds for restoration; secondary forests and fallows that are increasingly valued for providing ecological services (Guariguata and Ostertag 2001); and extensive areas of abandoned pasture. |
| The establishment of tree seedlings, however, was more dependent on the total number of slash-and-burn cycles imposed on the fallows than the number of years since the last abandonment. This is in line with other studies showing that intensity of previous land use affect recruitment (see Guariguata and Ostertag 2001). |
| A higher number of slash-and-burn cycles implicates more fire and soil disturbance, which subsequently remove more of the seedbank and potentially resprouting roots and stumps (Guariguata and Ostertag 2001). The decrease in seedling species richness and abundance with increased number of slash-and-burn cycles clearly suggests that reducing the number of slash-and-burn cycles before the land is left to recover, may shorten the establishment period for new tree seedlings. |
| Tree planting may be another tool to speed up the slow process of natural succession, and studies have shown that planted trees may have large effects on the recruitment of other tree species (see Guariguata and Ostertag 2001). Moreover, by selecting the most appropriate species for planting, both species richness and site quality can be restored (Guariguata and Ostertag 2001). |
| It is, however, expected that seed dispersal from the forest will be more important than resprout later in succession (Guariguata and Ostertag 2001). |
| After high disturbance intensity, succession may deviate from this general pattern (Guariguata and Ostertag 2001), but it is not clear how the increasing intensity of anthropogenic disturbances will affect these factors, and consequently, the resilience of tropical Sfs (secondary forests). |
| Given the importance of initial colonization in determining further successional pathways (Finegan 1996; Guariguata and Ostertag 2001) and the increasing extent of young SFs in agricultural landscapes (Metzger 2002; van Breugel et al. 2013), we focused on the first 5 years of forest succession. |
| Variation in both agricultural practices and the ecological matrix make it difficult to draw generalizations about forest recovery processes although some general patterns are apparent (Guariguata and Ostertag 2001). |
| Therefore, forest recovery at the landscape scale may depend on an interaction between land use history and underlying soil fertility (Guariguata and Ostertag 2001). |
| Guariguata and Ostertag (2001) point out that microhabitat differences and soil nutrients can strongly affect composition and growth of colonizing species during tropical secondary forest succession. |
| Height, basal area and species density increased logarithmically with stand age, as reported both for TWF (Guariguata and Ostertag 2001, Chazdon et al. 2007) and for TDF (Quesada et al. 2009). |
| SOM (soil organic matter) stabilizes soil aggregates, contributes to soil fertility (by holding on to organic forms of nutrients, and due to its high cation exchange capacity [CEC] that facilitates plant nutrient uptake), and increases the water-holding capacity of soils (Guariguata and Ostertag 2001), which may be more relevant for TDF (tropical dry forest) species distribution than stand age. |

| |
|---|
| EC (electric conductivity) also influenced community structure and is also related to CEC (Brady 1990), hence to soil fertility, which greatly influences rates of recovery of tropical forest structure (Guariguata and Ostertag 2001). |
| Chronosequence studies across the tropics demonstrate rapid increases in aboveground biomass during the first 4050 years of secondary forest succession, followed by either a slow, asymptotic increase toward old-growth values (Guariguata and Ostertag 2001), or a peak in biomass at intermediate stand ages (e.g., Marn-Spiotta et al. 2007, Letcher and Chazdon 2009, Mascaro et al. 2012). |
| Dominance of short-lived pioneer species has been hypothesized to occur in second-growth forests up to 20 years old, whereas long-lived pioneers are expected to dominate until a stand age of 100150 years (Guariguata and Ostertag 2001). |
| After planting exotic species on degraded lands, some native species are naturally recruited through seed dispersal, producing changes in plant diversity, composition and structure (Guariguata and Ostertag 2001). |
| In a tropical successional dynamic, the early successional plant community is characterized by grasses, shrubs and forbs that are later shaded by light-demanding pioneer tree species characterized by a short life cycle and a very fast growth rate. In more advanced stages, the canopy is dominated by tree species with a fast growth rate, a longer life cycle and taller stature but still requiring high light availability (Guariguata and Ostertag 2001). |
| Most of these trees species are unable to grow and reproduce under their own shade (Saldarriaga et al. 1988) and in consequence are replaced by shade-tolerant tree species in later successional stages (Guariguata and Ostertag 2001). |
| The increase in soil fertility over the succession may be attributable to the biomass accumulation and decomposition that leads to increased soil organic matter content and nutrients that are mineralized from organic matter turnover (Guariguata and Ostertag 2001). |
| Plant recolonization and species turnover are mainly determined by factors related to the type and intensity of previous land-use, such as soil conditions for germination, the presence of soil-stored seed and opportunities for seeds to disperse to the site (Guariguata and Ostertag 2001). |
| As intensity of previous land-use increases, the potential to regenerate from stored seed diminishes (Guariguata and Ostertag 2001). |
| The seed bank is likely to play a significant role in secondary succession at sites where land-use intensity before abandonment was low to moderate (Guariguata and Ostertag 2001). |
| Although secondary forests can achieve structural characteristics of old-growth forests in as little as 20 years (Guariguata and Ostertag 2001), less is known about long-term controls on soil C dynamics during succession. |
| These results are consistent with other studies that suggest shade-tolerant species tend to be dominant in secondary forests after 100400 years since abandonment (e.g. Guariguata and Ostertag 2001). |
| In diverse tropical forests, existing research has been more successful at predicting structural aspects of change (e.g. biomass accumulation, stem density) than in characterizing the dynamics of species composition during succession (Guariguata and Ostertag 2001). |
| Both successional and soil filtering processes thus contribute to maintaining species diversity in tropical forests, yet the mechanisms underlying these processes have mostly been studied independently (Guariguata and Ostertag 2001) |
| Soil properties change during forest succession (Guariguata and Ostertag 2001) |
| Many major changes in soil properties occur after disturbance and throughout the successional trajectory (Guariguata and Ostertag 2001). |
| From a functional perspective, ecosystems may recover functions long before they recover, if any, floristic similarity to previous condition (Guariguata and Ostertag 2001). |

| |
|---|
| Unlike tree fall gaps and other naturally disturbed areas, the regeneration of secondary forests on anthropogenically disturbed lands does not always follow predictable pathways (Guariguata and Ostertag 2001). |
| Once the forest reaches maximum basal area (20100 years) mangrove forests may not continue into stages characterizing old growth forests (¿100 years), which include enhanced plant diversity and the dominance of very large, shade tolerant trees (Guariguata and Ostertag 2001). |
| Forest regeneration depends on seed arrival, successful seedling establishment and growth. Seed arrival is typically related to proximity to old-growth or secondary forests (Finegan 1996, Guariguata and Ostertag 2001). However, even in cases with adequate seed sources, conditions in regenerating secondary forest may prevent seedling establishment or persistence (Guariguata and Ostertag 2001). |
| The last phase corresponds to the emergence of long-lived mature forest trees that germinate and establish under the cover of the previous species and develop following the disappearance of second stage species (Guariguata and Ostertag 2001). |
